# Identifying Differential Methylation in Cancer Epigenetics via a Bayesian Functional Regression Model

**DOI:** 10.1101/2021.03.21.436232

**Authors:** Farhad Shokoohi, David A. Stephens, Celia M.T. Greenwood

## Abstract

DNA methylation plays an essential role in regulating gene activity, modulating disease risk, and determining treatment response. Researchers can obtain insight into methylation patterns at a single nucleotide level utilizing next-generation sequencing technologies. However, complex features inherent in the data obtained via these technologies pose challenges beyond the typical big data problems. Identifying differentially methylated cytosines (dmc) or regions is one of such challenges. Current methodologies for identifying dmcs fall short in handling low read-depth data and missing values, capturing functional data patterns, granting multiple covariates (categorical, continuous, or combination), and multiple group comparisons. We have developed an efficient method to identify dmcs based on a Bayesian functional regression approach, termed `DMCFB`, that tackles these shortcomings. Through simulation studies, we establish that `DMCFB` outperforms current methods and results in better smoothing, and efficient imputation. We apply the proposed method to analyze a dataset containing patients with acute promyelocytic leukemia and control samples. With `DMCFB`, we discovered many new dmcs, and more importantly, exhibited enhanced consistency of differential methylation within islands and at their adjacent shores. Furthermore, we detected differential methylation at more of the binding sites of the fused gene involved in this cancer.

## 1. Introduction

DNA methylation at the fifth position in cytosine (5mC), an epigenetic mark found in many living organisms, has a variety of regulatory roles in disease and normal biology. Specifically, it regulates or causes transcriptional activity during embryonic development (Messerschmidt, Knowles and Solter, 2014), tissue development (Zhang et al., 2013), cell differentiation (Meissner et al., 2008), genomic imprinting (Li, Beard and Jaenisch, 1993), X-chromosome inactivation (Singer-Sam and Riggs, 1993), etc.

In this paper, we consider in particular the role of differential methylation of processes leading to the development of cancer; see, for example, Arteaga et al. (2015), Beggs et al. (2013). There is increasing evidence that epigenetic variation has a considerable influence on the regulatory processes that modulate cancer growth. The principal focus of our analysis is Acute Promyelocytic Leukemia (APL) (Liquori et al., 2020; Schoofs et al., 2013). APL is an aggressive subtype of acute myeloid leukemia (AML) that accounts for 10% of AML cases. APL is highly fatal in a short time with a severe bleeding tendency. A single mutation is known to cause APL, specifically a translocation involving two genes, the pml gene on Chromosome 15 and the rara gene on Chromosome 17 (Wang and Chen, 2008). Although the genetic cause is clear, how the genetic changes lead to dysregulation has been the subject of ongoing research (Liquori et al., 2020). For example, Schoofs et al. (2013) observed broad DNA hypermethylation in APL cells compared with control samples, and they argued that this may be a consequence of a loss of transcription factor binding. Epigenetic variation is also strongly believed to influence the choice of therapeutic intervention: the success of all-trans retinoic acid (ATRA) treatments as an alternative to chemotherapy has been marked (Arteaga et al., 2015), but such treatments are also known to vary in efficacy, with resistance to treatment being observed, due to mechanisms related to methylation. Of specific interest is the PML-RARalpha fusion protein (Jing, Xia and Waxman, 2002; Jing, 2004) which, when over-expressed due to the translocation, has been determined to have a leukemia-generating action. Therefore, understanding the modulation of regulation of expression of this protein is of key importance.

In light of the growing understanding the role of epigenetic variation in processes influencing tumorigenesis, there is a demand for both fundamental and clinical research on DNA methylation, and also a demand for analytical and statistical tools. Several assays have been developed to collect DNA methylation data (Yong, Hsu and Chen, 2016). Bisulfite sequencing (bs-seq) (Frommer et al., 1992) of DNA has become a popular technique that provides positive identification of 5mC residues in genomic DNA, particularly at CpG sites (where a cytosine nucleotide is followed by a guanine nucleotide in 5’–3’ direction). The technique benefits from the fact that bisulfite treatment will not affect 5mC, but converts unmethylated cytosines to uracils. Hence, after polymerase chain reaction and sequencing, the combined experiment leads to single base resolution information on methylation status by simply counting the number of times a sequencing read at a single genomic position appears as methylated versus unmethylated. Coupled with next-generation sequencing (ngs) (Behjati and Tarpey, 2013), bs-seq has become an effective tool to obtain single-nucleotide resolution data from the whole genome. Recently, a rapid decline in ngs costs made the whole-genome bisulfite sequencing (wgbs) (Lister et al., 2009) more accessible for research. Other direct assessment techniques include bc-seq (Hodges et al., 2009), bspp (Ball et al., 2009), and rrbs (Meissner et al., 2008). Although bs-seq is one of the most accurate technique to retrieve methylation data, it has several pitfalls that include false-positive methylation calls due to incomplete conversion, DNA degradation during bisulfite treatment, methylation in pseudogenes, and inability to discriminate between different methylated states such as 5mC and 5hmC (Wreczycka et al., 2017).

Much research has been conducted to provide efficient methods for quantitative analysis of DNA methylation data. One specific goal is to identify *differentially methylated cytosines* (dmc) or *regions* (dmr), as methylation is known to vary as a function of various biological and epigenetic factors. Hansen, Langmead and Irizarry (2012) (`BSmooth, bsseq`) used a binomial model with a local linear regression to smooth data and retrieve dmrs. Wu, Wang and Wu (2013) (`DSS`) utilized a Bayesian hierarchical model (Poisson, Gamma and log-normal) followed by the Wald test to capture dmrs. Wu et al. (2015) (`DSS-single`) and Feng, Conneely and Wu (2014) (`DSS`) followed a similar approach with different hierarchical models (Binomial, Beta and log-normal). Dolzhenko and Smith (2014) (`RADMeth`) combined a beta-binomial regression model with Stouffer-Liptak tests for dmr identification. Akalin et al. (2012) (`methylKit`) suggested using either logistic regression (capable of adding many covariates) or Fisher’s exact test. In Hebestreit, Dugas and Klein (2013) (`BiSeq`), a weighted local likelihood with a triangular kernel assuming a binomial probability function for methylation data is used. Jaffe et al. (2012) (`bumphunter`) applied linear mixed models to model methylation levels with the possibility of adding confounders. Several others (e.g., Hodges et al. (2011), Molaro et al. (2011), Song et al. (2013) (`MethPipe`), Saito, Tsuji and Mituyama (2014) (`Bisulfighter`), Saito and Mituyama (2015) (`ComMet`), Sun and Yu (2016) (`HHMFisher`), Yu and Sun (2016) (`HMM-DM`), and Shokoohi et al. (2019) (`DMCHMM`)) have focused on using versions of hidden Markov models to profile methylation data and to identify dmcs/dmrs. Lee and Morris (2015) (`WFMM`) proposed a wavelet-based functional linear mixed model that accommodates spatial correlations across the genome and correlations between samples through functional random effects. For more on the existing methods refer to the reviews by Bock (2012), Robinson et al. (2014), Klein and Hebestreit (2015), Chen, Lin and Fann (2016), Wreczycka et al. (2017), and Shafi et al. (2017), among others.

Several features common in the data that arise from assays such as bs-seq and rrbs lead to analytic and computational challenges. To illustrate some of these features, we explore a dataset containing samples from patients with acute promyelocytic leukemia (APL) and also samples from several different types of cells or tissues for comparison (Section 2.2). In these data, as in many other sequencing-derived methylation datasets, read-depths – the numbers of individual methylated and unmethylated counts recorded via the sequencing platform – vary appreciably and often unsystematically across cytosine positions; CpG sites are unevenly distributed (Lövkvist et al., 2016); both methylation autocorrelation between samples (Gallego-Fabrega et al., 2015) and between cytosines (Eckhardt et al., 2006) within a profile change irregularly across chromosomes. High heterogeneity has been observed in DNA methylation in cells from APL patients (Schoofs et al., 2013). Also, a large percentage of positions with missing values exists in the data, and the magnitude of the data leads to computational challenges for any kind of analysis.

In this work, we propose a new method, called `DMCFB`, built on a Bayesian functional regression model for the analysis of sequencing-based measures of DNA methylation. Despite a handful of existing methods for the analysis of DNA methylation data from sequencing experiments, `DMCFB` is more inclusive and addresses more of the data and computational challenges than any other method, as laid out below by comparing several existing methods. (i) *Missing values and imputation*: Sequencing data often contain many missing values; for example, in the apl data, 63% of the CpGs have missing values across samples (see Section 2.2). Almost all methods, except `DMCHMM` and `DMCFB`, remove all or most of the CpGs with missing values and then impute the rest. In `DMCFB`, we set the methylation read and read-depth to zero (i.e., *y* = 0*, n* = 0) for missing values in Binomial distribution, and impute methylation level *β* using the information from *p* neighboring points using a functional regression model. This approach gives a more efficient imputation than other methods including `DMCHMM` where an HMM and a Binomial distribution are utilized. (ii) *Read-depth*: Many measurements in sequencing data are based on only one or two reads, and in contrast, a few have unrealistically high read-depths. Several methods (e.g., `BiSeq`) filter CpGs with low and very high read-depths, whereas `DMCFB` keeps all available data and uses the read-depth information to adjust their contribution in model fitting. Also, since there is a systematic relationship between read-depths and methylation levels (i.e., CpGs with high read-depth tend to be more hypermethylated; see Supplementary Figure S3), `DMCFB` adds the extra covariate log(*n* + 1) in the model to account for such relationship. (iii) *Raw methylation level versus counts*: The aforementioned methods tend to model either methylation counts (*y, n*) or raw methylation levels *β*^raw^ = *y/n*. Modeling raw methylation levels may not fully capture the information (of read-depth) in the data. In `DMCFB`, we model (*y, n*) through logit(*β*) link with a Binomial(*y*; *n, β*) distribution to account for read-depth information. (iv) *Transformation*: Those methods that model *β*^raw^ often tend to use a transformation (e.g., log(*β*^raw^), or logit(*β*^raw^)). However, such transformations become undefined when dealing with CpGs that are fully methylated (*β*^raw^ = 1) or non-methylated (*β*^raw^ = 0), and either these data points must be removed prior to analysis, or a small constant must be subtracted/added to avoid NaN or ∞ values. `DMCFB` does not require a transformation of the raw data since it is built on a binomial emission model for the underlying proportion rather than the observed proportion. (v) *Functional pattern*: Methylation proportions are known to be highly correlated across nearby positions (Eckhardt et al., 2006) and samples (Gallego-Fabrega et al., 2015). The most efficient methods tend to use some type of smoothing to capture these correlations. Techniques range from weighted local likelihood (e.g., `BiSeq`), hidden Markov models (e.g., `DMCHMM`), to functional regressions (e.g., wavelet-based functional linear mixed model in `WFMM`). Here, `DMCFB` also builds on functional regression concepts. (vi) *Distance between CpGs*: CpGs are unevenly distributed across the genome (Lövkvist et al., 2016), and correlations between methylation levels decrease quickly with distance. `DMCHMM` and `WFMM` assume all positions are equally spaced which may under/overestimate autocorrelation; however, `DMCFB` incorporates the distance explicitly. (vii) *Sample characteristics*: In addition to these characteristics of the methylation data, existing methods take different approaches to test for the association between methylation levels and covariates. Some use a two-stage approach, first smoothing each sample and then testing on the smoothed data (e.g., `BiSeq`). Others perform two-group comparisons, i.e. testing for differential methylation between cases and controls (e.g., `HMMFisher`). `DMCFB`, on the other hand, does smoothing and dmc calling in one run. (viii) *Biological replicates*: A few methods (e.g., `DMRcaller`) tend to ignore biological variation across replicates which may increase type 1 error (Shafi et al., 2017). Recent methods (such as `WFMM`, `BiSeq`, and now `DMCFB`, etc.) can utilize such replicates. (ix) *Multiple covariates*: Usually, additional features including clinical information are collected about subjects in addition to the DNA methylation data. Several of the most widely used methods (e.g., `BiSeq`) are not capable of including these covariates in their statistical models. Most of those that do are incapable of accounting for multiple covariate types (that is, both categorical and continuous).

Furthermore, we have addressed several computational challenges including memory management, parallel computing, etc.; see Discussion (Section 7).

The rest of the paper is organized as follows. In Section 2, we provide brief descriptions of two motivating methylation datasets focussing on (i) data on three cell types extracted from whole blood and (ii) data from a study of acute promyelocytic leukemia. Our proposed Bayesian functional regression method to identify dmcs is given in Section 3. Section 4 covers a comparative simulation study inspired by a real-data structure. We provide analyses of the two real-datasets in Sections 5 and 6. Finally, Section 7 contains some concluding remarks.

## 2. Data

We use two publicly available datasets. One, which we refer to as the wgbs dataset, is a proof-of-principle dataset containing clear signals, and is used to develop our method and validate performance through simulation studies; the other, the apl dataset, contains data from patients with leukemia and will be extensively analyzed using `DMCFB`, `BiSeq`, and `DMCHMM`, and the results will be compared.

### 2.1. BLK DATA: WGBS *data on separated whole blood*

To develop our method we used a small part of a publicly available wgbs dataset derived from peripheral blood samples. Cell types were separated, and methylation profiles were estimated with whole-genome bisulfite sequencing (wgbs) in CD4^+^ T-cells, CD14^+^ monocytes, and CD19^+^ B-cells. We extracted data for a small region near the blk gene on human Chromosome 8, which is known to be hypomethylated in B-cells (Kulis et al., 2015; Shokoohi et al., 2019). The selected region contains 30,440 CpGs spanning 2 MB (10,352,236 - 12,422,082). The methylation status of 23.39% of positions in these data is missing. We have previously studied this dataset extensively – see Shokoohi et al. (2019). For more information see Supplementary Material S1.1.

### 2.2. APL DATA: RRBS *data on acute promyelocytic leukemia*

The second dataset that we analyze here was collected from patients with Acute Promyelocytic Leukemia (APL) (Schoofs et al., 2013) and is publicly available on the Gene Expression Omnibus (accession no. GSE42119). These rrbs data contain information on 18 APL patients and 16 control samples, with eight bone marrow samples from patients in remission (RBM), four profiles of healthy CD34^+^ cells (CD34), and four profiles from promyelocytes (PMC). Questions of interest include identifying regions where methylation is different among patients versus controls and also looking at different cell types and/or disease status (“active” or “in remission”).

The dataset is challenging to analyze. It includes 9,335,693 genomic positions for each of 34 samples. Approximately 63% of CpGs have missing values across samples. Almost all existing methods will remove either all 63% CpGs or may impute a proportion of them. This approach leads to unreliable results due to the loss of much useful information as a result of the removal of many positions from the analysis. Three out of 16 samples in the control group and 12 out of 18 samples in the APL group are females. Age ranges between 20 and 83 years. On Chromosome 15, there are 285,437 positions available, and 64.27% of CpGs across the samples have at least one missing value. Furthermore, nearly 6.19% of CpGs in the control group and about 32.31% of CpGs in the APL group have missing values in all samples. On Chromosome 17, there are 510,386 positions available, and among these, there are 59.81% of positions with at least one missing value, and all samples are missing in 5.55% of CpGs in the control group and 28.91% of CpGs in the APL group. Refer to Supplementary Material S1.2 for more on the data.

A specific focus in our analysis will be the locations of binding sites of the protein PML-RARalpha, for which (Schoofs et al., 2013) lists some 225 locations on Chromosome 17. Epigenetic variation in the vicinity of these sites might indicate modulation of the expression of this protein, and so detection of differential methylation in these sites may lead to key insights.

## 3. Method

We propose a Bayesian functional regression profiling method to identify dmcs. In the following, we describe the main aspects of the proposed method including the introduction of a functional representation of the methylation profiles incorporated into a generalized linear model, the possible statistical inference approaches, several numerical challenges in fitting the model and obtaining parameter values, and the procedure by which a position is classified as a dmc.

We regard the methylation counts as realizations of a *Binomial*(*n*(*t*), *β*(*t*)) process observed on the interval [0, *T*] which represents some large genomic region, where the read-depth *n*(*t*), *t* ∈ [0, *T*] may vary as the result of the sequencing process or the underlying propensity for methylation and may be platform dependent. In this paper, we condition on the read-depth data, thereby ignoring their stochastic properties. We specify that

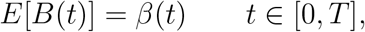

where *B*(*t*) = *Y* (*t*)*/n*(*t*), and essentially consider a (functional) model for {*B*(*t*)}. We have access to replicate data {*Y*_*i*_(*t*), *t* ∈ [0, *T*]}, *i* = 1, …, *m*, and potentially fixed covariates. We consider a functional representation of *β*(*t*) using a natural cubic spline basis in *t*; other spline bases may also be used. Our representation allows a further decomposition into group-specific effects when a grouping structure is present.

### 3.1. A spline model for methylation data

We respect the discrete nature of the observation grid (given by nucleotide position) and regard the observation locations as positive integers *t* = 1, …, *T*. We denote the methylation data of a given sample profile by {(*y*_*t*_, *n*_*t*_), *t* = 1, …, *T*} where *y*_*t*_ and *n*_*t*_ respectively represent the methylation read-count and read-depth at the *t*^*th*^ genomic position. A reasonable, if imperfect, model assumes that

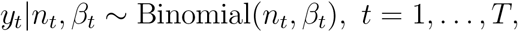

where *β*_*t*_ is the propensity for each site *t* to be methylated at each read instance. In this model, each read is assumed to yield a binary outcome that is conditionally independent of the other reads. The methylation level at position *t*, *β*_*t*_ changes due to different sources of variation. In the blk data, for example, there are three cell-types (B-cell, T-cell, and monocyte). In the apl data, there are four types of samples (CD34, PMC, RBM, and APL), as well as the individuals’ age and sex. Thus there is individual-level variation, group variation, and possible variation influenced by additional covariates, and these can be incorporated in a binary regression model

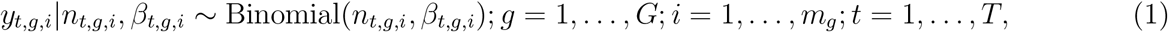

where *G* is the number of groups in the variable of interest, *m*_*g*_ is the number of samples (individuals) in the *g*^*th*^ group, and *T* is the number of cytosines (CpGs) in a sample.

Due to the DNA methylation autocorrelation structure and the unknown functional form of methylation patterns in different regions, we propose a spline-based functional (logistic) regression model. Assume the vector **x** is formed from a *p*-dimensional (natural cubic) spline basis derived from methylation positions.

We then decompose *β*_*t,g,i*_ in (1) as

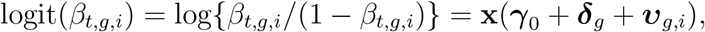

where ***γ***_0_ represents the baseline group-level parameter, ***δ***_2_, …, ***δ***_*G*_ are the group-specific contrasts by setting ***δ***_1_ ≡ **0**, and ***υ***_*g,i*_ are the individual-level variations. Note that ***γ***_0_, ***δ***_*g*_, and ***υ***_*g,i*_ are all *p* × 1 vectors. Accordingly, the design matrix for any individual is denoted as **X**_*g,i*_ which is a *T* × 3*p* block matrix. The entire design matrix **X** that includes all groups (*G*) and all individuals 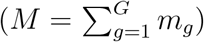 is of dimension *MT* × (*G* + *M*)*p*. The dimension of the spline basis, *p*, is termed the *resolution* and is equivalent to a *band-width* parameter. Note that other splines bases will produce fairly similar results. Finally, the model can be augmented to

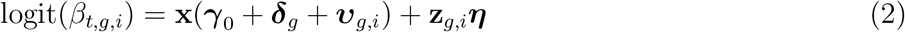

for some non-positional and non-spline-related fixed effects, where ***η*** is a *q* × 1 vector.

### 3.2. Modelling the impact of read-depth

In this work, we emphasize the importance of adding an extra covariate based on the read-depth at each position, as we have observed that read-depth varies systematically with methylation levels and that higher read-depths tend to result in larger methylation levels (Supplementary Figure S3). For instance, a plausible model for the blk data is given as

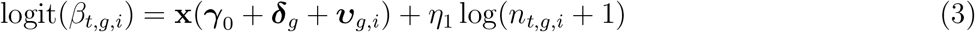

which allows for the effect of read-depth in an additive fashion on the linear predictor scale. As described above, in this analysis we treat read-depth as a deterministic quantity and perform a conditional analysis, even though the stochastic properties of the read-depth process might also be of interest in other settings.

### 3.3. Inference

To estimate the model parameters either the maximum likelihood (ML) or the Bayesian estimation approach may be used. Maximum likelihood estimates are useful in a fully Bayesian analysis as they allow for the Bayesian computational strategy to be implemented more efficiently by providing initial estimates for the Bayesian procedure.

#### 3.3.1. Maximum likelihood estimation

The log-likelihood function is given by

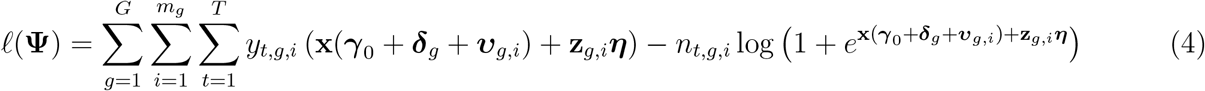

where 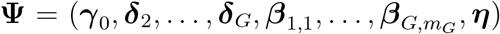 and 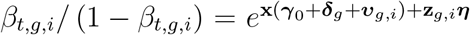. By setting the first partial derivatives of (4) to zero, and verifying that the matrix of second partial derivatives is negative definite and that the solution is the global maximum, one can derive a numerical solution as 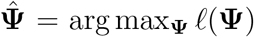. For generalized linear models (GLM), even at a large scale, the ML approach can be implemented effectively using R packages such as `fastglm` (Huling, 2019). However, we prefer a fully Bayesian approach as it yields a more complete representation of associated uncertainties in light of the data and model specification.

#### 3.3.2. Bayesian estimation

The full Bayesian model is given as follows. Denote the estimated logistic scale methylation levels by ***β***\* = [*β_t,g,i_*]. We propose the model represented by the factorization

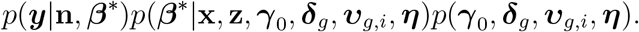

We allow for a deterministic relationship between (**x**, **z**, ***γ***_0_, ***δ***_*g*_, ***υ**_g,i_*, ***η***) and ***β***\*, specifically that at each site, logit(*β_t,g,i_*) follows the linear model in (2). By choosing suitable priors, inference for the collection of unobserved quantities (***γ***_0_, ***δ**_g_*, ***υ**_g,i_*, ***η***) and ***β**\** is attained using computational methods based on Markov Chain Monte Carlo (MCMC).

### 3.4. Numerical methods and techniques

We briefly describe the numerical methods required for inference in the following sections.

#### 3.4.1. Iteratively re-weighted least squares

Although the maximum of (4) can be drawn numerically, the cumbersome process involves solving a system of nonlinear equations, and as is usual for GLMs we adopt an iteratively re-weighted least squares (IRLS) approach: beginning with a tentative solution we apply the recursion

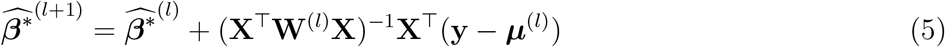

until there is no appreciable change to the estimates.

#### 3.4.2. Quasi-IRLS

Due to the high-dimensionality of the calculation of **X**^*T*^**W**^(*l*)^**X** in (5), there are typically issues with required memory, however, one can utilize the block-structure of **X** to speed up the process, or resort to minorization-maximization (MM) procedures or quasi-IRLS which simplifies the iterative estimation procedure by setting **W**^(*l*)^ equal to the identity for each *l* in (5), that is, without the requirement to recompute the weight matrix **W** and the inverse Hessian (**X**^*T*^**W**^(*l*)^**X**)^−1^; in a GLM it is known that the MM procedure converges to the maximizer of the log-likelihood. These procedures are appreciably faster to compute but can be slower to converge to the ML estimate.

#### 3.4.3. Bayesian inference

For fully Bayesian inference we adopt an MCMC approach and perform block updates of the effect-specific parameters. The Gibbs sampler is used to collect MCMC samples; we devise a Gibbs sampler algorithm by introducing independent Gaussian priors on the model parameters, although this can be easily relaxed. The priors on all parameters are chosen to be independent 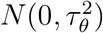 priors. The hyper-parameter *τ_θ_* can be specified subjectively, but it is also common to adopt an empirical Bayes approach, and choose *τ_θ_* after inspection of the data. We describe one such empirical approach below. In general, it is straightforward using simulation to study the impact of different prior specifications; elicitation can be readily carried out by simulation from the prior, and reconstructing the prior for the profiles by computing the *β* values in (3).

The Gibbs sampler updates are structured as block updates for the term-specific parameters, ***γ***_0_, ***δ**_g_*, ***υ**_g,i_*, ***η***, conditional on the other parameters. In each update, the full conditional posterior distribution corresponds to a posterior from a Bayesian binary regression model, and this update can be achieved efficiently using `bayesglm` in the `arm` library in R (Gelman and Su, 2020). There are three possible strategies: a full update for one parameter block may involve

1. multiple ‘inner’ accept-reject Metropolis-Hastings steps to sample the full conditional posterior; this provides an exact update of the block which should lead to faster convergence in the ‘outer’ chain;
2. a single ‘inner’ accept-reject Metropolis-Hastings steps to sample the full conditional posterior; this also provides an exact update of the block but may lead to slower convergence in the ‘outer’ chain;
3. an exact update using the approximate full conditional distribution given by the Normal approximation to the non-Normal exact version, that is, from the standard approximation 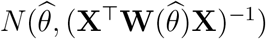 inspired by the quadratic approximation of the log-likelihood function.

The third approach essentially appeals to a Gaussian approximation to the full posterior distribution, and we have found that in fact, the Gaussian approximation works well in most cases. The Gibbs sampler approach – rather than a joint update of all parameters simultaneously – is usually more feasible in large-scale problems. Finally, it is generally not feasible to build a complete functional regression genome-wide. Instead, we use a partitioning approach that fits the model in smaller genomic segments, tailored to the computational resources available (specifically, the available physical RAM and number of cores). The R package we have developed optimizes the segment size on an automatic basis. For the details of the implementation of our method refer to Algorithms 1-9 in Supplementary Material S3.

### 3.5. *Identifying* DMC*s*

Having gathered MCMC samples for parameters at position *t*, we compute the 100(1 − *α*)% credible interval for each (multiple) comparison in the categorical variable of interest. If at least one of the intervals does not contain zero at position *t*, the position is classified as a dmc. The process is repeated for all positions. In addition to the (overall) dmc, the pairwise dmcs are also reported when there are more than two categories in the variable of interest.

## 4. Simulation Study

We have assessed the performance of the proposed method in simulations inspired by the cell-separated data described in Section 2.1. In what follows we provide simulation scenarios and settings, selected methods for comparison, simulation results, and sensitivity analysis.

### 4.1. Scenarios and settings

We use the identical scenarios and settings for simulation proposed in Shokoohi et al. (2019). The methylation information available in the BC and monocyte samples were chosen to generate simulated data as follows: (a) First, read-depths then methylation counts of all BC samples were aggregated. (b) Next, the missing information was imputed using nearby count information. (c) Using ‘`lowess`’ in `R`, we fitted a smooth curve (span = 0.05) denoted G1; this curve was used to specify a baseline profile. (d) Using Supplementary Table S2, we generated the second group curve called G2 by adding effect sizes to the chosen regions of G1. (e) By fitting a Normal distribution to the methylation difference between G1 and all BC samples we obtained *μ* ≈ 0 and *σ* ≈ 0.18. We used these estimates to add extra variation to G1 and G2 for all samples. (f) Read-depths of all BC and monocyte samples were chosen to generate 500 simulated datasets, where within each dataset we generated 8 samples from G1 and 13 samples from G2 by doing the following steps: (i) Additional variation at each site was generated using *N* (0, 0.18) random errors which were added to the methylation levels in G1 and G2. (ii) All values are truncated to fall in [0, 1]. (iii) Methylation counts were generated by multiplying the generated methylation levels in the previous step. The integer parts of these data were chosen as the final methylation counts. Supplementary Figure S4 presents the graph of the dmrs, one simulated dataset that compares it to the methylation in the real dataset.

Several of the most prominent and widely-used analysis tools were chosen for comparison including bsseq (Hansen, Langmead and Irizarry, 2012), `bumphunter` (Jaffe et al., 2012), `BiSeq` (Hebestreit, Dugas and Klein, 2013), `DSS` (Feng, Conneely and Wu, 2014), `DMRcaller` (Zabet and Tsang, 2015), `HMM-DM` (Yu and Sun, 2016), `HMM-Fisher` (Sun and Yu, 2016), `WFMM` (Lee and Morris, 2015) and `DMCHMM` (both non-weighted and weighted approaches) (Shokoohi et al., 2019). Refer to Supplementary Material S4 for a brief comparison of the chosen methods.

### 4.2. Simulation results

Quantification of performance of each method is based on the following definitions: True Positive (TP, CpG is correctly identified as dmc); True Negative (TN, CpG is correctly identified as non-dmc); False Positive (FP, CpG is incorrectly identified as dmc); False Negative (FP, CpG is incorrectly identified as non-dmc). ‘Sensitivity’ (SE) of dmrs and ‘Specificity’ (SP) of non-dmrs for each simulated dataset *r* ∈ {1, …, *R*} are defined as

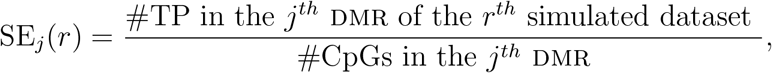

for *j* = 1, …, 10, and

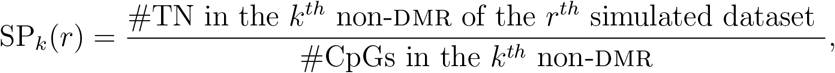

for *k* = 1, …, 11, where *R* = 500. The average ‘Sensitivity’ of the *j^th^* dmr and the average ‘Specificity’ of the *k^th^* non-DMR are respectively calculated as 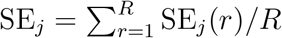 and 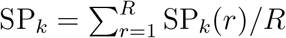. The accuracy is calculated as ACC = {#TP + #TN} #CpGs for each method. We also report the modified accuracy (MACC) by assuming that dmr1 through dmr8 are the only real dmrs as given in Shokoohi et al. (2019).

We set the nominal false discovery (FDR) threshold at 5% for any applicable method. We modified the default parameters in each tool to match simulation settings and to increase their efficiency. `WFMM` does not impute missing data; instead of removing the positions with missing values, we used a naive approach based on neighboring information to impute them in the simulated datasets to get higher performance. The results of `DMCFB` are based on normal priors with mean zero and precision 10 for all parameters, and choosing *p* = 30 as the resolution (band-width). We set *α* = 5 × 10^*−*2^ for the Bayesian credible intervals in `DMCFB`.

Figure 1 depicts the ACC and MACC results for different methods. `DMCFB` performs better than the existing methods. Figure 2 illustrates the average ‘Sensitivity’ and the average ‘1-Specificity’, separated for each dmr and non-dmr. `DMCFB` is superior in terms of ‘Sensitivity’. The results for ‘1-Specificity’ show that `DMCFB` is either better than or comparable to other methods. One advantage of `DMCHMM` is the ability to detect small differences in methylation profiles (dmr1, dmr8), and large differences with high precision (dmr2-7) (Shokoohi et al., 2019). Our proposed method outperforms `DMCHMM` in this regard. Supplementary Figure S6 presents Cohen’s Kappa. Our new method outperforms the competing methods. Supplementary Figure S7 shows the proportions of the times that the start and end of a dmr are identified as dmcs. `DMCFB`’s results are higher than those of other methods. The empirical FDR (eFDR) in Supplementary Figure S8 shows that `DMCFB` attains the nominal FDR.

**Fig 1:**
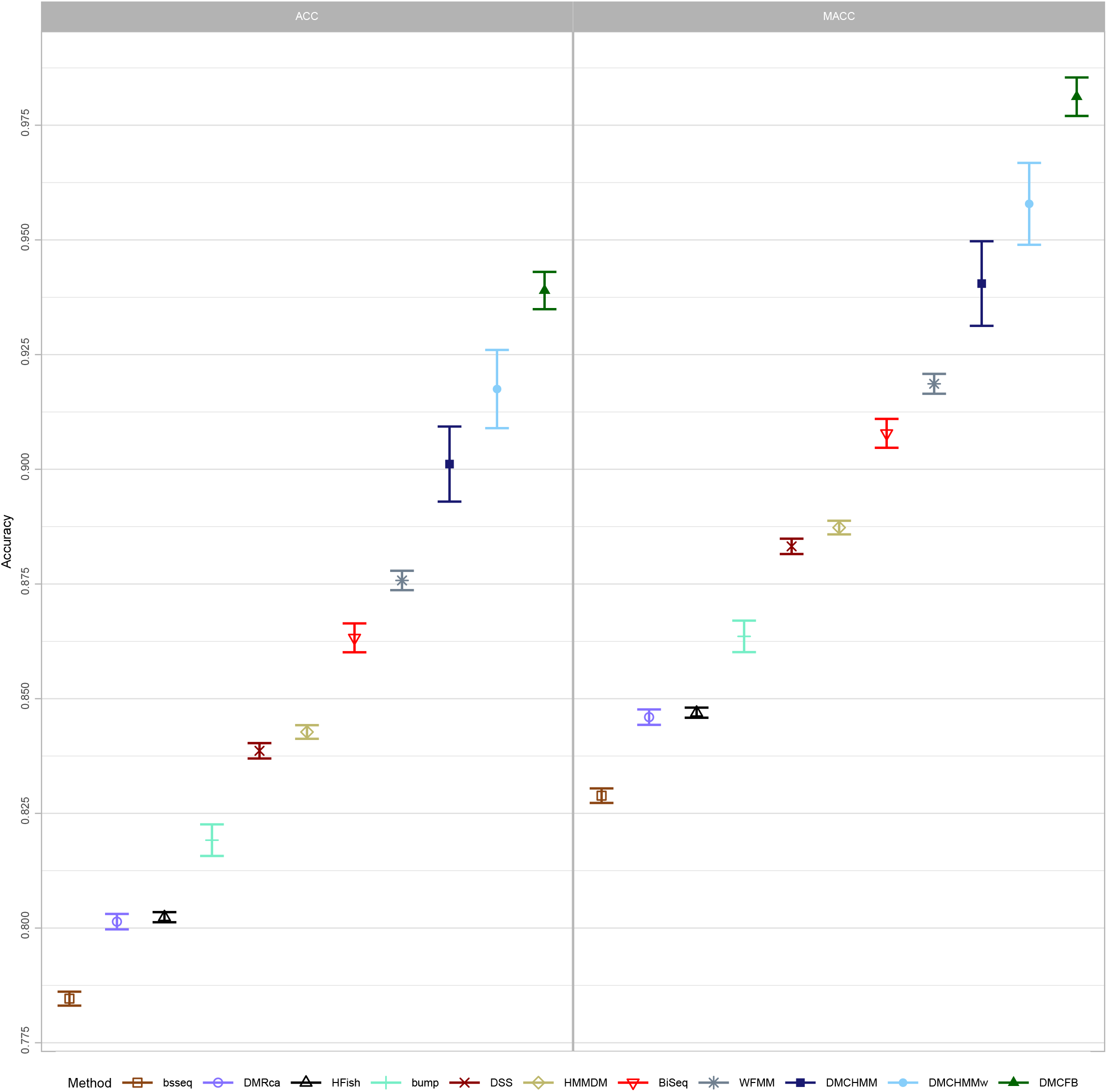
The average overall accuracy (ACC) and average overall modified accuracy (MACC) for different methods in simulated data for the first scenario; Errors are generated from *N* (*μ* = 0, *σ* = 0.18). Relevant results are reproduced from Shokoohi et al. (2019). (sd error bars are added.)

**Fig 2:**
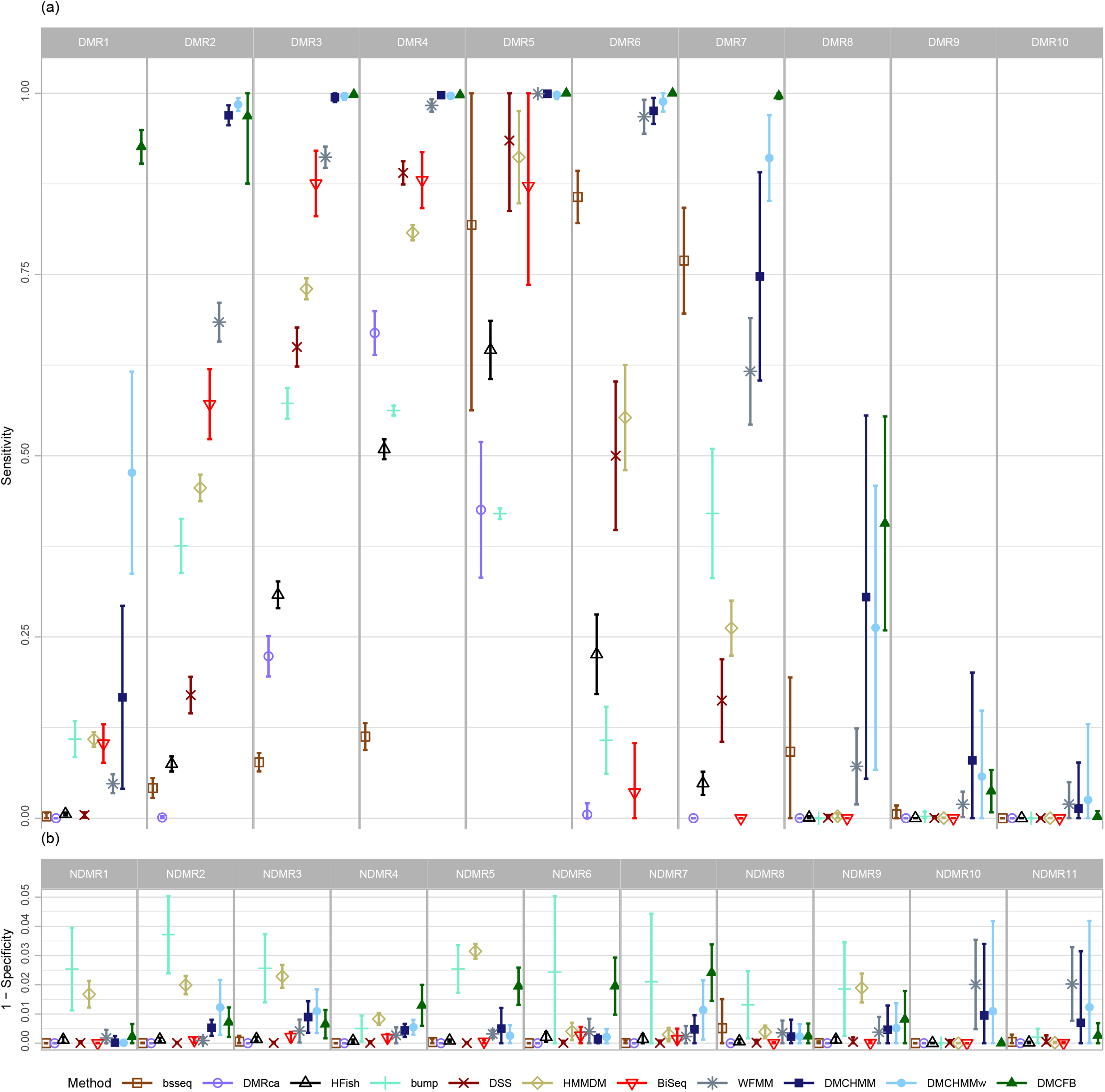
(a) The average proportion of correctly identified dmcs (‘Sensitivity’) for each method separated by dmrs; (b) The average proportion of incorrectly identified dmcs (‘1-Specificity’) for each method separated by ndmrs (The axis is truncated) in simulated data for the first scenario; Errors are generated from *N* (*μ* = 0, *σ* = 0.18). Relevant results are reproduced from Shokoohi et al. (2019). (sd error bars are added.)

#### 4.2.1. Sensitivity Analysis

Since our proposed Bayesian approach depends on the choices of band-width and prior, we performed sensitivity analyses concerning both specifications. First, in terms of band-width, we have chosen a wide range of values in {20, 25, …, 60} to study its impact. Figure 3 compares ‘Sensitivity’ and ‘1-Specificity’ of `DMCFB` using different choices of band-width. From this figure and Supplementary Figures S19-S22, we can conclude that the results are fairly robust for the band-width selection, specifically when the effective size is large. Secondly, concerning prior precision, we examine Gaussian priors with different precisions, 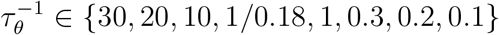, for sensitivity analysis: it transpires that, overall, better results are observed with respect to eFDR if the precision is estimated from the data using an empirical Bayes approach. In this case, the average empirical site-specific standard deviation proves to provide a reasonable scaling, leading to the specification *τ_θ_* = 0.18. Figure 4 and Supplementary Figures S28-S31 show the robustness of the results.

**Fig 3:**
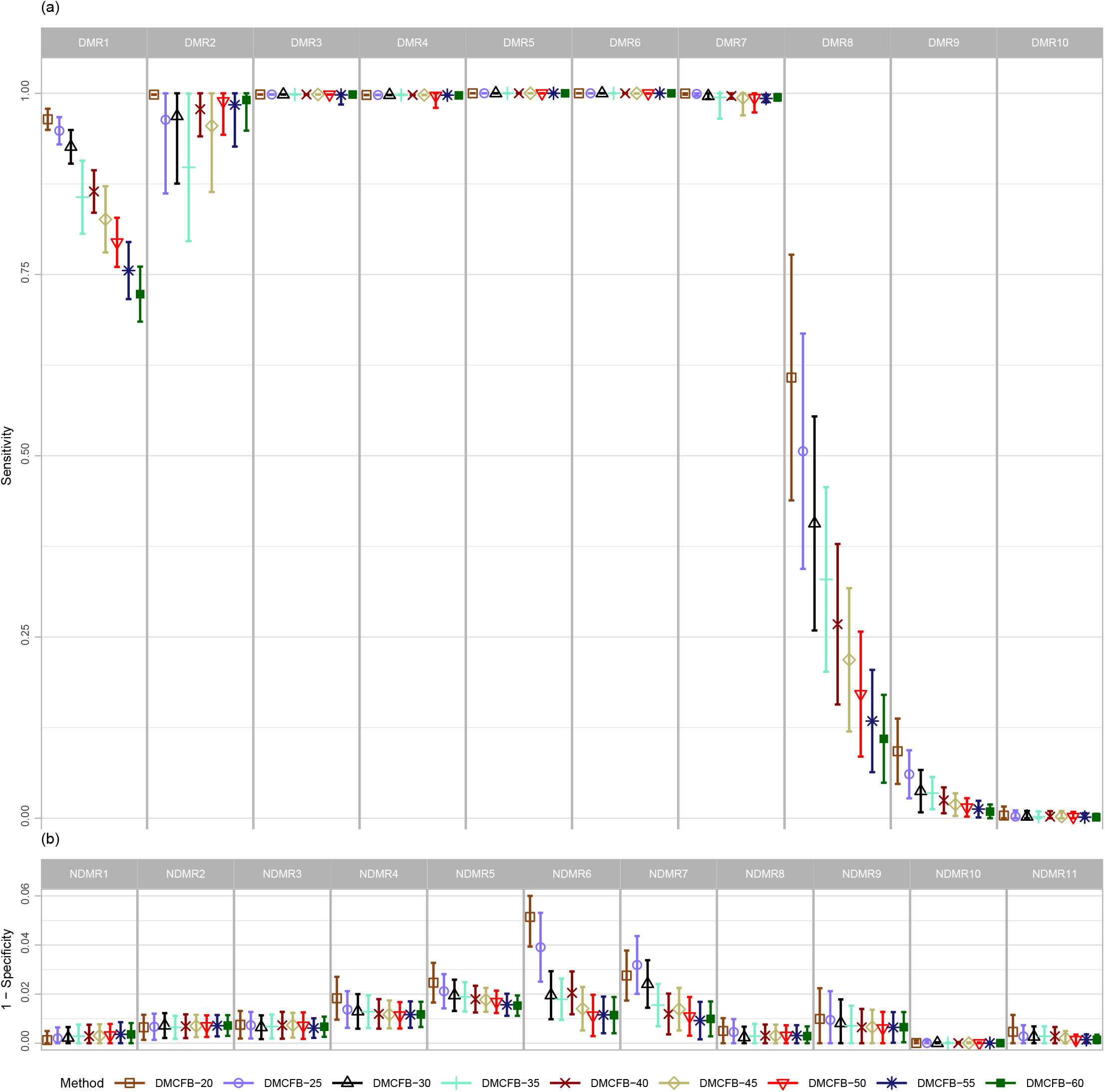
(a) The average proportion of correctly identified dmcs (‘Sensitivity’) for each choice of band-width separated by dmrs `DMCFB`; (b) The average proportion of incorrectly identified dmcs (‘1-Specificity’) for each choice of band-width separated by ndmrs in `DMCFB` (The axis is truncated) in simulated data for the first scenario; Errors are generated from *N* (*μ* = 0, *σ* = 0.18). (sd error bars are added.)

**Fig 4:**
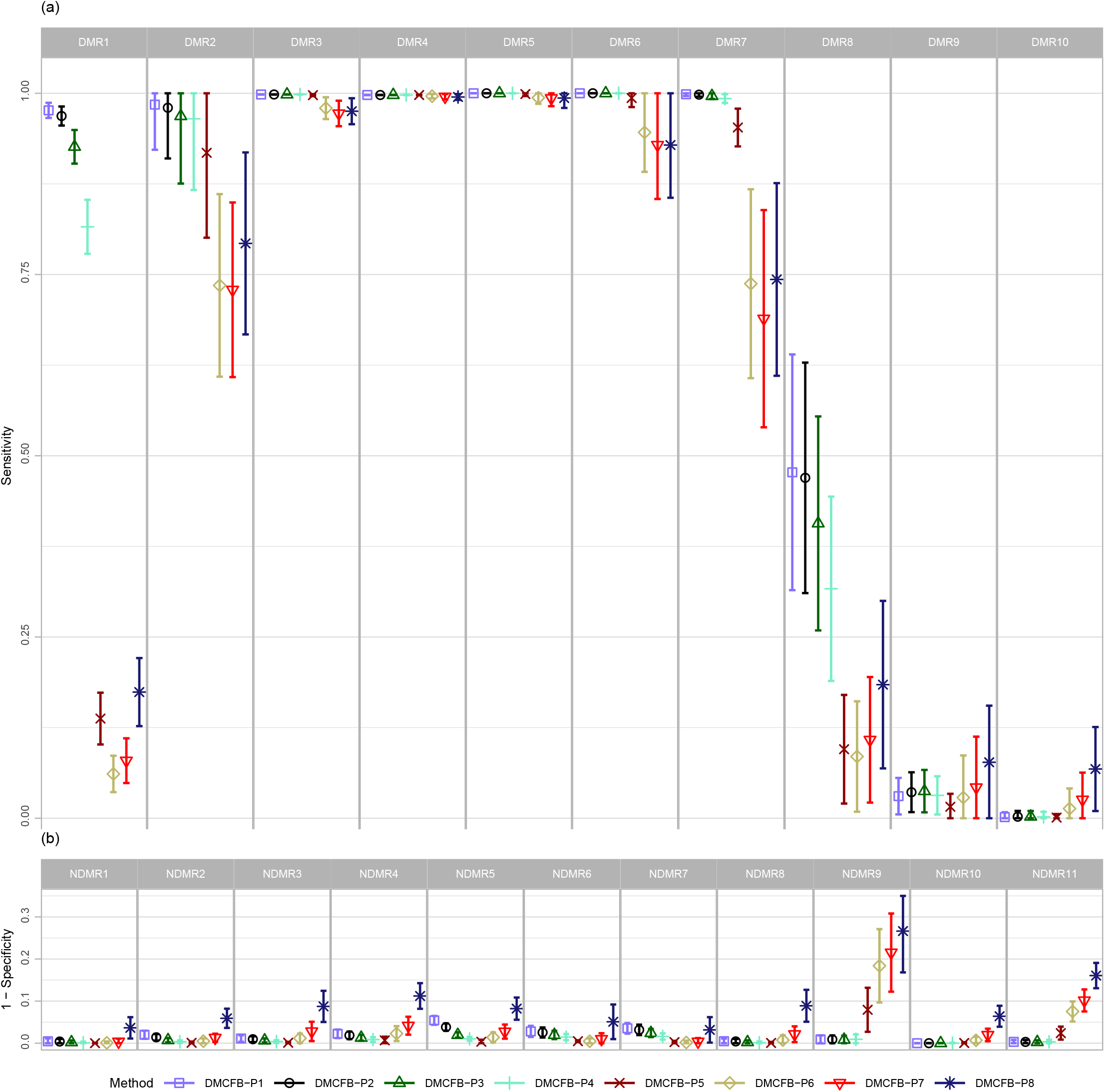
(a) The average proportion of correctly identified dmcs (‘Sensitivity’) for each choice of precision of prior separated by dmrs in `DMCFB`; (b) The average proportion of incorrectly identified as dmcs (‘1-Specificity’) for each choices of precision of prior separated by ndmrs in `DMCFB` (The axis is truncated) in simulated data for the first scenario. Errors are generated from *N* (*μ* = 0, *σ* = 0.18). The precision is chosen, in order, in {30, 20, 10, 1/0.18, 1, 0.3, 0.2, 0.1}. (sd error bars are added.)

A summary of conclusions drawn from several simulation studies is as follows. (i) `DMCFB` outperforms all considered methods in almost all criteria including accuracy, Cohen’s Kappa, ‘Sensitivity’, and the percentage of detecting start and end of dmrs. (ii) In terms of ‘1-Specificity’, the `DMCFB` method is either comparable or better than that of other methods. (iii) Apart from bumphunter, `HMMDM`, and `BiSeq`, all other methods including `DMCFB` attained the nominal FDR at 5%. (iv) Similar to `DMCHMM`, `DMCFB`’s performance is excellent in dmrs with large effect sizes (say 0.2) and does not depend on any other features. (v) For small effect sizes (say 0.1), our method’s performance is superior to other methods including `DMCHMM`. (vi) `DMCFB` does a better job in smoothing the data followed by `DMCHMM` and `WFMM`. Specifically, `DMCFB` is robust concerning the length of dmrs and the size differences; see dmrs 2-7. (vii) Similar results are observed using larger variability for noise (*σ* = 0.24) (see Supplementary Material S5.3). The decline in performance using our method is negligible. This result shows that `DMCFB` is robust with regard to variation in the data while other methods have noticeable performance loss. (viii) Using a different simulation setting with dmrs of different lengths, we observed trends similar to those described above (see Supplementary Material S5.4). (ix) Sensitivity analysis shows `DMCFB` is robust for band-width selection. (x) A sensitivity analysis is done by re-assembling regions to see how a regulatory landscape is altered by the grouping factor. The results are observed to be robust (see Supplementary Material S6.4). (xi) Sensitivity analysis concerning the precision of prior distribution shows the robustness of the results. More specifically, better eFDR is observed when the prior precision is estimated from data using an empirical Bayes approach.

## 5. Proof of Principle Analysis of Sorted Cells

The blk data resembles a dataset in which several challenges such as multiple groups, variable read-depths, and missing values, exist jointly. Almost none of the existing methods can efficiently handle such complexity by addressing all the known challenges. To this end, we have compared the methylation profiles between cell types near the blk gene described in Section 2.1 using `DMCFB`. The blk data include 8 BC, 13 monocyte, and 19 TC samples in which methylation counts and read-depths for 30,440 CpGs are available. We used normal priors with *τ_θ_* ≈ 0.18. The value of *τ_θ_* is computed by obtaining the standard deviation (SD) of methylation levels among samples at each position, and then averaging the SD values over all the positions. The band-width is set to 30. The fitted model is identical to (3), where BC is the baseline, *n_t,g,i_* is the read-depth at position *t* of sample *i* in group *g* where *t* = 1*, …, T* = 30, 440, *g* = 1, 2, 3 and *i* = 1, …, *m_g_* with *m*_1_ = 8, *m*_2_ = 13 and *m*_3_ = 19.

A set of 1000 MCMC samples is collected after a burn-in of 1000, which is more than enough to achieve convergence. We set *α* = 5 × 10^−2^, 1 × 10^−5^ and 1 × 10^−8^ in constructing the credible intervals (Supplementary Table S5); hence, the CpG site at position *t* is classified as a dmc if at least one of the (1 − *α*)% credible intervals for group comparisons does not include zero.

For *α* = 1 × 0^*−*8^, we observed in total 36.14% of CpGs were identified as dmc (Supplementary Table S5). Among these, approximately, 18.46%, 14.37%, and 24.17% of CpGs were differentially methylated respectively for pairwise comparisons BC vs monocyte, BC vs TC, and monocyte vs TC. Figure 5 depicts the result of this analysis for the region near the blk gene promoter. From this figure, we can observe that the three cell-types are differentially methylated specifically at the promoter of the blk gene. These results are in concordance with the literature (Kulis et al., 2015). It is worth mentioning that most analytical tools are incapable of imputing missing values efficiently and analyzing all three groups in the blk data at once, i.e. performing three simultaneous comparisons.

**Fig 5:**
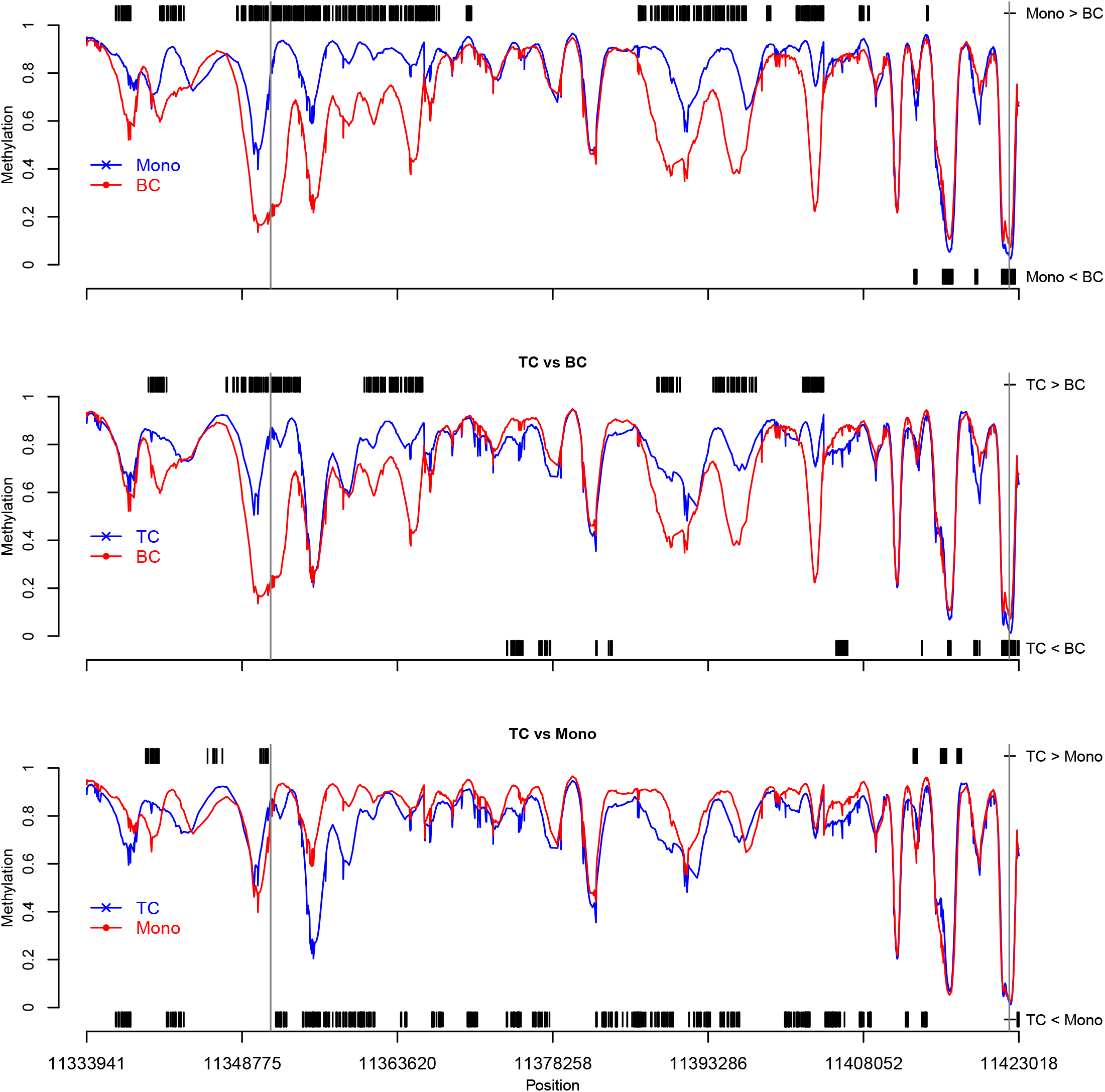
(a)-(c) Identified dmcs for three pairwise comparisons of cell-type specific methylation data near the blk gene promoter. Short black vertical lines indicate CpGs where one cell type was significantly different from the other, at credible interval level *α* = 1 × 10^−8^. The average methylation level for each cell-type is also plotted.

## 6. Analysis of Patients with Acute Promyelocytic Leukemia Versus Control Cell Types

Schoofs et al. (2013) described differences in methylation between patients with APL and three types of normal cells: mononuclear cells from remission bone marrow, CD34^+^ cells from healthy donors, and promyelocytes derived in vitro from CD34^+^ cells. APL is known to be caused by a Chromosome 15-17 translocation affecting promyelocytic leukemia-retinoic acid receptor *α*, and these authors explored, in rrbs data, how methylation patterns differed across the genome. Using rrbs profiling, Schoofs et al. (2013) showed widespread hypermethylation in APL samples, not only restricted to the location of the translocation.

Here, we use `DMCFB` to reanalyze the data on Chromosome 15 and 17, to see if additional sensitivity can be achieved. From the point of view of building an analytic approach for these data, the apl data present several significant challenges: (i) A large proportion of the data points is missing. Most other methods cannot handle such complexity and will remove either all or big chunks of positions. (ii) Additional covariates such as age and sex are of interest, and this may force users to choose among a few methods that can handle multiple covariates. (iii) Read-depths vary dramatically in different regions; few methods account for such information. (iv) It can be seen from Supplementary Figure S2 that the methylation pattern changes dramatically in different regions; simple smoothing techniques cannot efficiently capture such complexity.

We can conclude that the approaches that are taken in Schoofs et al. (2013) (i.e., removal of most positions with missing values and low read-depth and not including additional sources of variation in age and sex) have probably led to a set of weak or unreliable results. Hence, we reanalyzed the apl dataset using `DMCFB` and compared the results with those of `BiSeq`.

We do two reanalyses of the data: First, we perform a simple case-control comparison without considering any additional covariates and grouping the three types of controls (CD34, PMC, and RBM) together. This allows a direct comparison of the results by `DMCFB` with the results by `BiSeq` presented by Schoofs et al. (2013) (Section 6.1). The second analysis compares the four groups of samples and includes covariates (Section 6.2).

### 6.1. *Reanalysis of* APL *data by comparing DMCFB and BiSeq*

The fitted model in `DMCFB` is set to

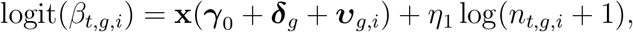

where ***γ***_0_ is the baseline group level (Control), ***δ***_2_ (APL) is the group-specific contrast by setting ***δ***_1_ ≡ **0**, ***v**_g,i_* are the individual-level variations, **x** is the (natural cubic spline transformation) design matrix, *n_t,g,i_* is the read-depth at position *t* of individual *i* in group *g* for *t* = 1, …, *T* (*T* = 285, 437 and 510, 386 for Chromosome 15 and 17, respectively), *g* = 1, 2 and *i* = 1, …, *m_g_* with *m*_1_ = 16 control samples and *m*_2_ = 18 APL patients.

We used Gaussian priors with *τ_θ_* ≈ 0.12. The value of *τ_θ_* is computed by obtaining the SD of methylation levels among samples at each position, and then averaging the SD values over all the positions. The band-width is set to 30.

We collected 1000 MCMC samples after a burn-in of 1000 runs, which is more than enough for convergence. For calling genome-wide significance in genetics, the thresholds of *α* = 1 × 10^−5^ or *α* = 1 × 10^*−*8^ are commonly used (Lander and Kruglyak, 1995). These thresholds were derived to control the family-wise error rate at 5% for genetic data. Although an appropriate threshold for wgbs data is not known, here we choose a threshold of *α* = 1 × 10^−8^ and note that very similar numbers of dmcs were identified if we used the more liberal threshold. For `BiSeq`, we first used the significance level *α* = 1 × 10^−8^, but this identified only a very small number of dmrs (and consequently a very small number of dmcs). Therefore, for our `BiSeq` analysis, we used significance thresholds of *α* = 1 × 10^−1^ and *α* = 5 × 10^−2^ for testing and trimming clusters, respectively.

Tables 1(A, B) show how and when `DMCFB` and `BiSeq` agree or disagree for Chromosomes 15 and 17, respectively. The tables show the numbers of dmcs called for the two methods, along with the percentages conditional on an annotation (Islands, Shores, and Deserts). Firstly, in Table 1(A), we observe a much smaller number of sites are called dmc by `BiSeq` relative to `DMCFB`. The significance thresholds used for `BiSeq` will affect this proportion, but the level we have chosen is already quite liberal (*α* = 5 × 10^*−*2^). Secondly, we can see that if `BiSeq` calls a dmc, then `DMCFB` will call the same site as a dmc 80.6% of the time. This proportion is fairly consistent across the three site annotations, although it is a little lower for Desert regions than those of Islands and Shores. In contrast, we can see substantial differences in performance among the sites that `BiSeq` does not call as dmcs. Overall, the `DMCFB` method calls 33% of these sites as dmcs. But this percentage ranges from 25.6% for CpG Deserts to 51.9% for the CpG Islands. In Table 1(B), similar findings are obtained. The agreement with `BiSeq` calls is a little higher 83.4% instead of 80.6%, and a similar trend is seen among sites not called by `BiSeq`, such that `DMCFB` is calling more dmcs in Islands and Shores than in Desert regions. **Figures 6(A,B)** look at coherence within Islands. If an Island is identified as differentially methylated by `DMCFB` it is more likely that most of the CpGs inside the Island are identified as dmc compared to `BiSeq`, and this finding indicates better smoothing by `DMCFB` as we saw in simulation studies.

**Fig 6:**
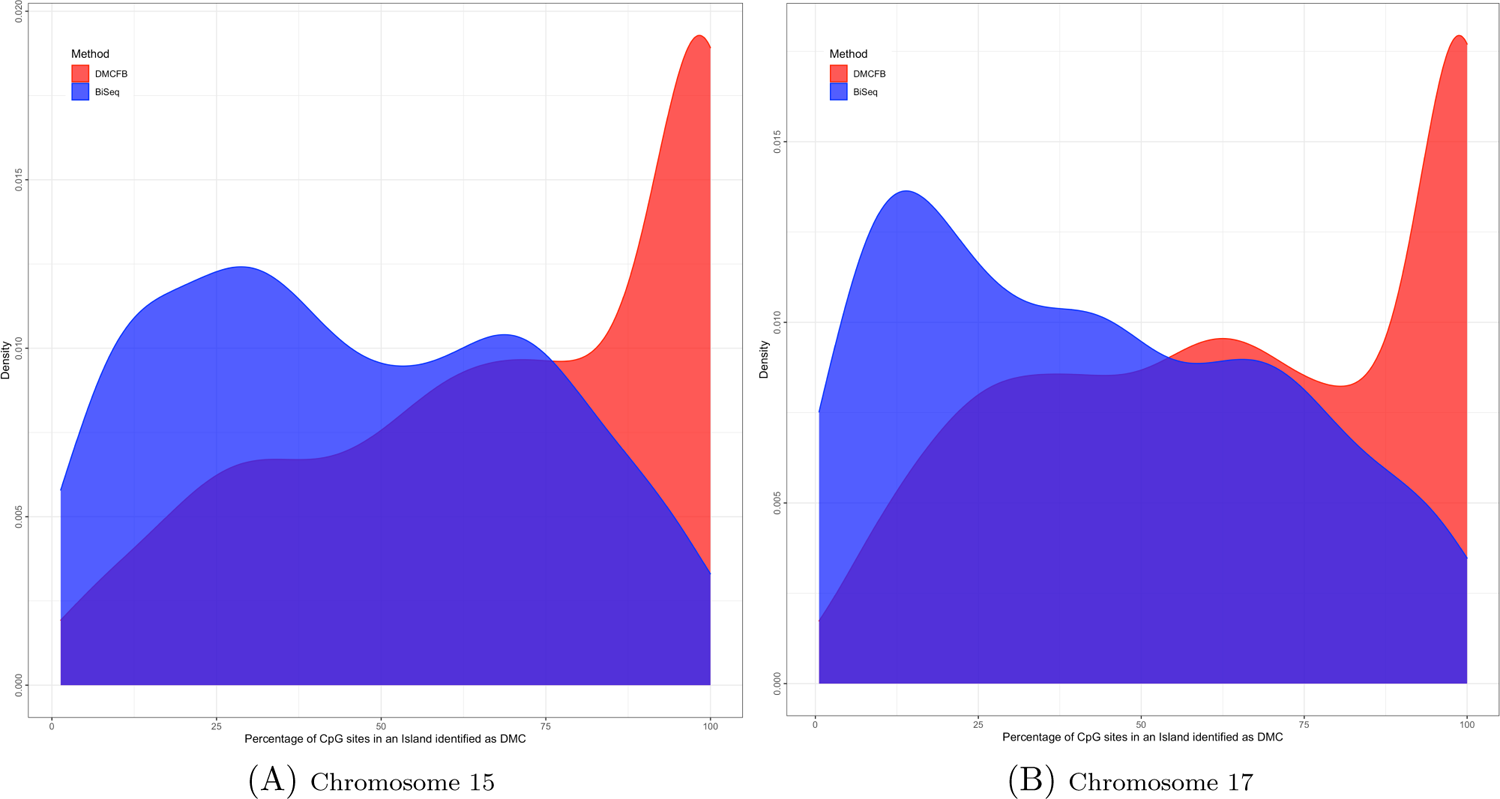
Density of percentage of CpGs in Island identified as DMC by DMCFB (red) and BiSeq (blue)

**Table 1.**
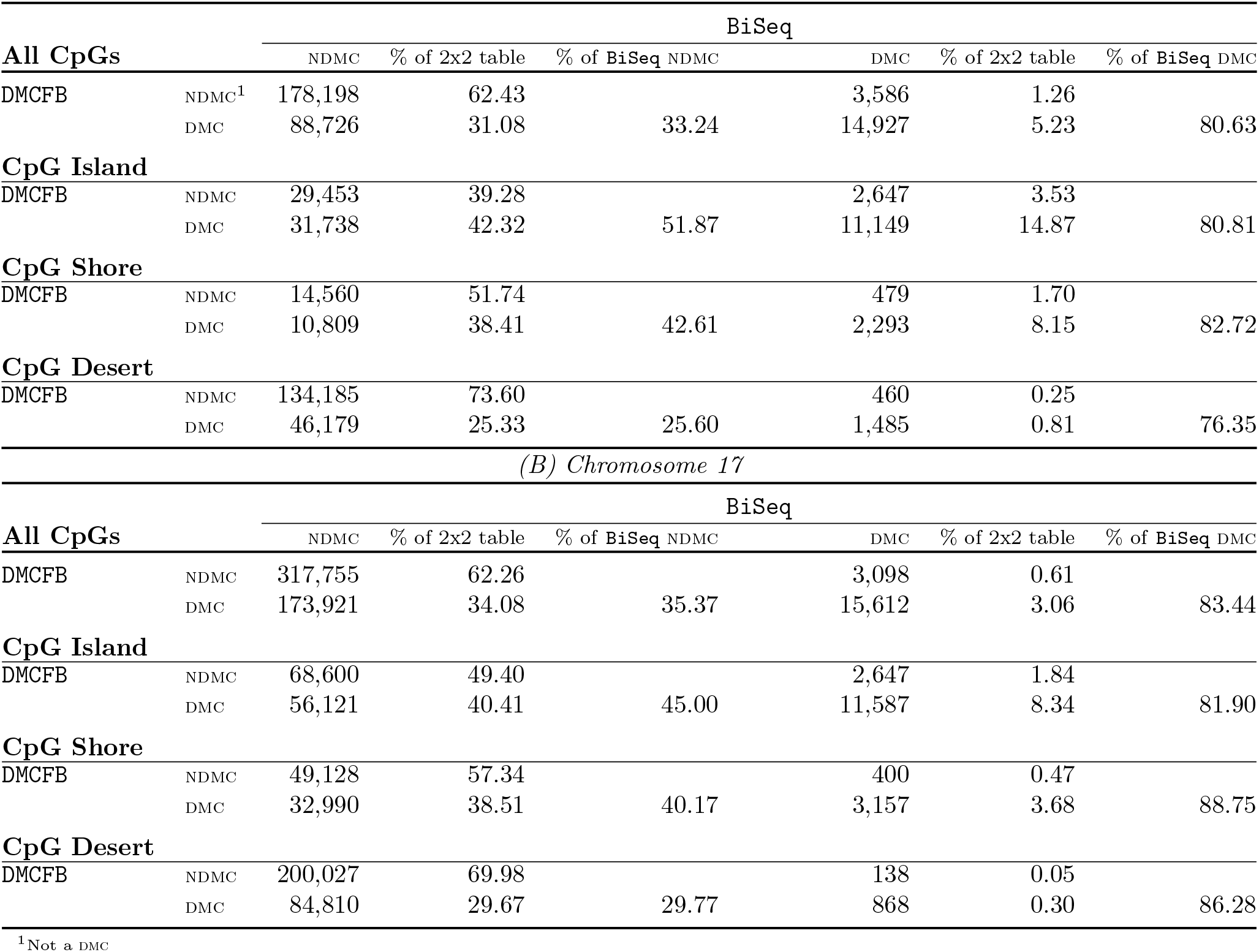
*Comparing `DMCFB` and `BiSeq` in identifying* dmc*s in the* APL *data (A) Chromosome 15*

We now address the detection of differential methylation in the vicinity of PML-RARalpha binding sites. Supplementary Table S8 presents which of the known binding sites listed in Martens et al. (2010) are identified as differentially methylated, when comparing APL and controls, by either method. We observe that `DMCFB` identified more binding sites as differentially methylated than did `BiSeq`. On Chromosome 15, `DMCFB` identified 14 sites, while `BiSeq` identified only 2 sites as differentially methylated; on Chromosome 17, `DMCFB` identified 43 sites, while `BiSeq` identified only 2 sites. We also display the read-depth of the known binding sites that are not identified as differentially methylated by either method or identified by only `DMCFB` in Supplementary Figures S38(A, B) on Chromosome 15 and Supplementary Figures S39(A, B) on Chromosome 17. From these figures, we observe that when neither method detects a dmc (at known binding sites), often the read-depth is very low, and so the capability to detect dmcs is compromised. However, when the read-depth is moderate or high, `DMCFB` identifies appreciably more binding sites as differentially methylated. In addition, `DMCFB` exhibits more sensitivity for capturing the documented widespread dysregulation.

Additional analytic results are found in Supplementary Material S8.1, and here we highlight some features of these additional results:

- Supplementary Table S9 examines the agreement between adjacent Islands and Shores and shows whether they agree. On both analyzed chromosomes, if an Island contains at least one dmc, its adjacent Shore is more likely to be identified as differentially methylated by `DMCFB` than `BiSeq`, suggesting increased sensitivity of `DMCFB`. For example, on Chromosome 15 with `DMCFB`, 61.9% of sites with at least one dmc in an Island also identified at least one dmc in the adjacent Shore. In contrast for `BiSeq`, this percentage was only 32.6%. Similarly, if an Island was not differentially methylated, its adjacent Shore is also more likely to be not called differentially methylated by `DMCFB` than `BiSeq`.
- Supplementary Table S10 addresses the direction of the methylation changes, overall and by annotation. When both methods identify a CpG as dmc, they mostly agree on the direction of the methylation, and if they do not agree on the direction, `BiSeq` tends toward hypermethylation. If a CpG site is detected as dmc by only `DMCFB`, the direction is mostly hypomethylation in Islands and hypermethylation in Shores and Deserts. If a CpG site is detected as dmc by only `BiSeq`, the direction is mostly hypomethylation in both Islands and Shores. These findings are similar for both Chromosomes 15 and 17.
- Supplementary Figures S40 and S41 represent read-depth for dmcs captured by `DMCFB`, `BiSeq`, or both. Clearly, median read-depth is lower for `DMCFB`, which speaks to our method’s efficient use of information. When both methods call a dmc, read-depth tends to be very high in the APL samples - i.e., a strong clear signal for differential methylation.
- Supplementary Figures S42-S45 illustrate the difference between the raw data and the smoothed data, and also show how APL’s methylation levels are often much higher than among controls. The differences between patients and controls become far more apparent after smoothing, and overall, `DMCFB` captures far more sites as differentially methylated than `BiSeq`.

We present a comparison between `DMCFB` and `DMCHMM` in Supplementary Material S8.3.

### 6.2. *Reanalysis of* APL *data by accounting for all the information using DMCFB*

In this section, we reanalyze the apl data using `DMCFB` but account for the additional information of sex, age, and the four groups (CD34, PMC, RBM, and APL). The full model is set to

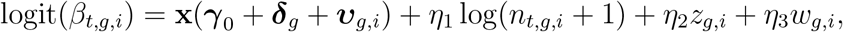

where *γ*_0_ is the baseline group level (CD34), ***δ***_1_ ≡ **0**, ***δ***_2_ (PMC), ***δ***_3_ (RBM), and ***δ***_4_ (APL) are the group-specific contrasts, ***v**_g,i_* are the individual-level variations, **x** is the (natural cubic spline transformation) design matrix, *n_t,g,i_* is the read-depth at position *t* of individual *i* in group *g* with its effect *η*_1_, *z_g,i_* is the sex of individual *i* in group *g* with its effect *η*_2_, and *w_g,i_* is the age of individual *i* in group *g* with its effect *η*_3_. Recall *t* = 1*, …, T* (*T* = 285, 437 and 510, 386 for Chromosome 15 and 17, respectively), *g* = 1,...,4 and *i* = 1*, …, m_g_* with *m*_1_ = *m*_2_ = 4, *m*_3_ = 8, and *m*_4_ = 18 for the four groups CD34, PMC, RBM, and APL, respectively.

A set of 1000 MCMC samples is collected after a burn-in of 1000, which is more than enough to achieve convergence. We set *α* = 5 × 10^−2^, 1 × 10^−5^ and 1 × 10^−8^ in constructing the credible intervals. The results are presented in Tables 2(A, B) for Chromosome 15 and 17, respectively. The patterns in both chromosomes and all three *α* levels are almost similar, specifically for *α* = 1 × 10^−5^ and *α* = 1 × 10^−8^. This analysis, therefore, leads to six age- and sex-adjusted pairwise comparisons across the four sample types. The most number of dmcs are found for the contrast APL versus CD34, followed by APL versus PMC and APL versus RBM. Overall, 3.626% of CpGs identified as dmc using *α* = 5 × 10^−2^.

**Table 2.** *Reanalysis of the* APL *data using `DMCFB` by accounting for all available informations; Chromosome 15 and 17.*

(A) Chromosome 15
| $\alpha$ | Pairwise contrasts | | NDMC<sup>1</sup> % | DMC % | Hyper % | Hypo % |
| --- | --- | --- | --- | --- | --- | --- |
| $\alpha = 5 \times 10^{-2}$ | PMC | CD34 | 98.814 | 1.186 | 0.651 | 0.535 |
|  | RBM | CD34 | 99.722 | 0.277 | 0.165 | 0.113 |
|  | APL | CD34 | 98.371 | 1.629 | 1.142 | 0.487 |
|  | RBM | PMC | 99.847 | 0.153 | 0.066 | 0.087 |
|  | APL | PMC | 99.612 | 0.388 | 0.135 | 0.253 |
|  | APL | RBM | 99.364 | 0.635 | 0.369 | 0.266 |
|  | All positions |  | 96.374 | 3.626 | — | — |
| $\alpha = 1 \times 10^{-5}$ | PMC | CD34 | 99.977 | 0.023 | 0.022 | 0.001 |
|  | RBM | CD34 | 99.983 | 0.017 | 0.016 | 0.001 |
|  | APL | CD34 | 99.832 | 0.168 | 0.139 | 0.029 |
|  | RBM | PMC | 99.979 | 0.021 | 0.015 | 0.006 |
|  | APL | PMC | 99.976 | 0.024 | 0.009 | 0.015 |
|  | APL | RBM | 99.975 | 0.025 | 0.018 | 0.007 |
|  | All positions |  | 99.768 | 0.232 | — | — |
| $\alpha = 1 \times 10^{-8}$ | PMC | CD34 | 99.977 | 0.023 | 0.022 | 0.001 |
|  | RBM | CD34 | 99.983 | 0.017 | 0.016 | 0.001 |
|  | APL | CD34 | 99.832 | 0.167 | 0.139 | 0.029 |
|  | RBM | PMC | 99.979 | 0.021 | 0.015 | 0.006 |
|  | APL | PMC | 99.976 | 0.024 | 0.009 | 0.015 |
|  | APL | RBM | 99.975 | 0.025 | 0.018 | 0.007 |
|  | All positions |  | 99.769 | 0.231 | — | — |

(B) Chromosome 17
| $\alpha$ | Pairwise contrasts | | NDMC<sup>1</sup> % | DMC % | Hyper % | Hypo % |
| --- | --- | --- | --- | --- | --- | --- |
| $\alpha = 5 \times 10^{-2}$ | PMC | CD34 | 98.930 | 1.070 | 0.607 | 0.462 |
|  | RBM | CD34 | 99.597 | 0.403 | 0.185 | 0.218 |
|  | APL | CD34 | 97.897 | 2.103 | 1.555 | 0.548 |
|  | RBM | PMC | 99.650 | 0.350 | 0.159 | 0.191 |
|  | APL | PMC | 99.320 | 0.681 | 0.377 | 0.303 |
|  | APL | RBM | 98.985 | 1.015 | 0.554 | 0.461 |
|  | All positions |  | 95.611 | 4.389 | — | — |
| $\alpha = 1 \times 10^{-5}$ | PMC | CD34 | 99.941 | 0.059 | 0.007 | 0.051 |
|  | RBM | CD34 | 99.998 | 0.002 | 0.002 | 0.001 |
|  | APL | CD34 | 99.765 | 0.235 | 0.215 | 0.020 |
|  | RBM | PMC | 99.976 | 0.024 | 0.024 | 0.000 |
|  | APL | PMC | 99.891 | 0.109 | 0.093 | 0.016 |
|  | APL | RBM | 99.892 | 0.108 | 0.093 | 0.015 |
|  | All positions |  | 99.691 | 0.309 | — | — |
| $\alpha = 1 \times 10^{-8}$ | PMC | CD34 | 99.941 | 0.059 | 0.007 | 0.051 |
|  | RBM | CD34 | 99.998 | 0.002 | 0.002 | 0.001 |
|  | APL | CD34 | 99.765 | 0.235 | 0.214 | 0.020 |
|  | RBM | PMC | 99.976 | 0.023 | 0.023 | 0.000 |
|  | APL | PMC | 99.891 | 0.109 | 0.093 | 0.016 |
|  | APL | RBM | 99.892 | 0.108 | 0.093 | 0.015 |
|  | All positions |  | 99.691 | 0.309 | — | — |

<sup>1</sup>Not a DMC

It is worth noting that almost all the analytical tools are incapable of analyzing the apl data by comparing all six contrasts in one run, and considering all additional information on subjects while keeping all the CpGs in the analysis and imputing the missing values.

## 7. Discussion

In this article, we have proposed an efficient DNA methylation approach, `DMCFB`, for dmc identification based on a functional data model and a Bayesian estimation and inference procedure. We have demonstrated the superiority of this method over existing methods through exhaustive simulation studies as well as robustness via sensitivity analysis concerning both band-width and prior selection. The proposed method is flexible in terms of adding any source of variation and is robust with respect to the true underlying methylation pattern. Missing values are automatically imputed. The method is capable of adding discrete or continuous covariates or combinations. Most existing methods ignore read-depth information in the analysis; by adding the read-depth as an extra source of variation in the model, our proposed method adjusts the methylation levels for better estimation. Similar to `DMCHMM`, our proposed method shows consistent behavior in the sense that the results depend on the difference in methylation between groups and not other aspects of dmrs such as length, location, autocorrelation pattern, read-depth, etc. One drawback of `DMCHMM` is that the efficiency deteriorates if the distances between CpGs are explicitly included in the HMM model. Our proposed method is not restricted in this sense.

While reanalyzing the apl data we noted that `BiSeq`’s default significance threshold maintains a very stringent control over the false discovery rate, and only a small number of dmcs are identified; in fact, `DMCFB` identifies overall 6 (Chromosome 15) or 10 (Chromosome 17) times more dmcs than `BiSeq`. However, it is possible to adjust the `BiSeq` package options slightly, and we ran another series of analyses where the `BiSeq`’s call was 7.5% (Supplementary Table S11(A)) instead of 6.5% (Table 1(A)) on Chromosome 15. However, the results of this sensitivity analysis showed less agreement with `DMCFB`, i.e. among dmcs called by `BiSeq`, the percentage also called by `DMCFB` dropped from 81.63% (Table 1(A)) to 68.79% (Supplementary Table S11(A)) on Chromosome 15. In contrast, among CpGs not called by `BiSeq`, the percentage of dmcs called by our method remained essentially unchanged from Table 1. This suggests that when `BiSeq` is less confident in a call, that the two methods are using nearby reads and methylation patterns differently. Furthermore, the ability to capture more dmcs in CpG Islands reflects `DMCFB`’s ability to use information from adjacent CpGs via our functional smoothing. Given the density of CpGs in the Islands, we obtain much more sensitivity in these regions. `DMCFB`, due to the functional regression and efficient imputation, seems to capture better a signal that persists across several CpGs. `DMCFB` often identifies all CpGs in an Island as differentially methylated, whereas `BiSeq` tends to capture less than half. Similarly, we have shown that if an Island displays differential methylation, that `DMCFB` more often also finds differential methylation in the adjacent Shores. Similarly, the additional sensitivity of `DMCFB` is also visible when examining the known binding sites of PML-RARalpha; performance of our approach was particularly notable when the read-depths were moderate. The two methods are both smoothing, but they are using the information differently.

We now elaborate on several computing challenges and the techniques implemented in `DMCFB` as in which we addressed them.

- *Partitioning*: We partition data for two reasons: one, to avoid using a very large dimensional design matrix which results in very high-dimensional computation and memory problems; two, to utilize the multi-core facility provided in many computing machines which increases the speed of the process. Note that by partitioning the data into smaller segments, a smaller resolution can be used. We recommended partitioning the data into regions of size 500 CpGs and using a single resolution of 30 in each partition for all parameters.
- *Parallel computing*: Implementing parallel computing results in faster computation. Users can use two approaches for parallel computing to speed up the process: (1) running the computation on several partitions simultaneously; (2) using further parallel computing while estimating *β_t,g,i_* parameters, both of which are implemented in `DMCFB`.
- *Multi-resolution modeling*: To speed up the process one can use a multi-resolution model choosing different band-widths for the model parameters (***γ***_0_, ***δ**_g_*, ***υ**_g,i_*). For example, one may choose a larger band-width for ***υ**_g,i_* and a smaller one for (***γ***_0_, ***δ**_g_*).
- *Software*: We provide a universal R package optimized by using several fast libraries and codes in R and C by following the Bioconductor guidelines.

In summary, our proposed method provides an improved, robust, and flexible method for DMC/DMR identification.

## Supplementary Material

Web supplement including tables, figures and extra data analyses referenced here are available with this paper at the Annals of Applied Statistics on the *ims* website. Our open-source package `DMCFB` is available at http://bioconductor.org/packages/DMCFB/.

## Acknowledgements

This work is partially supported by a grant awarded to the first author by the Center of Biomedical Research Excellence (COBRE) under COBRE grant number P20GM121325. The second author was funded by a Discovery Grant from the Natural Sciences and Engineering Research Council of Canada (NSERC).

