## Supplementary material for "Identifying Differential Methylation in Cancer Epigenetics via a Bayesian Functional Regression Model": Web-based Supplement: Identifying Differential Methylation in Cancer Epigenetics via a Bayesian Functional Regression Model (from another version of this preprint: version 1, 2021-03-22)

**Farhad Shokoohi**[1], **David A. Stephens**[2] **and Celia M.T. Greenwood**[3]

These Supplementary Materials contain eight sections as follows:

- Section S1 gives brief information about the two methylation data sets discussed in the main paper.
  - We plot the methylation patterns in both data sets and we show that the patterns are too complex for almost all methods to handle.
- Section S2 displays a figure related to the BLK data set.
  - We show that, in general, methylation level has a systematic relationship with read-depth.
- Section S3 presents details of algorithms developed for the `DMCFB` method.
  - The algorithms presented here explain in detail the computational techniques/approaches we adopted to implement our proposed statistical method, and also show that how we managed to optimize the memory and CPU of different computing systems to be able to handle high-dimensional data.
- Section S4 compares the selected methods in the main paper.
  - We presented a table in which a few of the capabilities of existing methods are compared.
- Section S5 presents the results of the simulation study discussed in the main paper.
  - We show the simulation setups for different scenarios and plot the actual DMRs along with simulated data and real data for one sample (Section S5.1).
  - We present the rest of the results for simulation study Scenario 1 with $\sigma = 0.18$ (Section S5.2). We show that the results indicate that our proposed method, i.e. `DMCFB`, is superior to the existing methods by using different criteria.
  - We present all the results related to Scenario 1 with $\sigma = 0.24$ (Section S5.3) which indicate results similar to the previous scenario. Although there is a drop in the performance in all methods, the decline in our method is negligible.
  - We present all the results for simulation study Scenario 2 (Section S5.4). We show that similar to previous results, `DMCFB` outperforms all the existing methods.
- Section S6 gives the sensitivity analysis for `DMCFB`.
  - We show that the results are generally robust to the choice of bandwidth (Section S6.1 and S6.2), although precision decreases slightly as the bandwidth increases.

  - We show that the results are generally robust to the choice of prior (Section S6.3), although an empirical approach gives better performance.
  - We show that the results are generally robust to the simultaneous choice of band-with and partitioning (Section S6.4).
- Section S7 gives the rest of the details of the BLK data analysis discussed in the main paper.
  - We show that there is widespread hypomethylation in B-cells compared with T-cells and monocytes.
- Section S8 provides details of the APL data analysis and comparisons between `DMCFB`, `BiSeq`, and `DMCHMM`.
  - In Section S8.1, we give additional results from the reanalysis of the APL data in the main paper.
  - In Section S8.2, another comparison between `DMCFB` and `BiSeq` using the APL data is given which confirms the results in the main paper.
  - In Section S8.3, a comparison between `DMCFB` and `DMCHMM` using the APL data is given which shows less agreement between the two methods in some criteria and more agreement in some other criteria compared with those of `BiSeq`.

#### Notation and Acronyms

The following notation and acronyms are used in the main paper and the supplementary materials.

- Plat. : Platform based on which methylation data are generated;
- r.NA : Missing values are removed;
- filter : Data are filtered to remove positions with low read-depth;
- Clustering : Clustering sites prior to analysis;
- Smoothing : Smoothing methylation levels;
- Parallel : Parallel Computing;
- Stats. : Providing Statistics such as p-values, FDRs, etc;
- BSS : BS-Seq technology for retrieving methylation status;
- BiSeq : The `BiSeq` method
- bsseq : The `bsseq` method
- bump : The `bumphunter` method
- DMCFB : The `DMCFB` method
- DMCHMM : The `DMCHMM` method without weights
- DMCHMMw : The `DMCHMM` method with weights
- DMRca : The `DMRcaller` method
- DSS : The `DSS` method
- HFish : The `HMM-Fisher` method
- HMMDM : The `HMM-DM` method
- bed : A file type for storing methylation data
- NDMR, NonDMR : Not a DMR
- NDMC : Not a DMC
- STR, END : Start and end position of a DMR

#### S1. Two methylation data sets

Two publicly available data sets are used in the main paper. The BLK data set is used to develop our method (`DMCFB`) and to perform simulation studies. For further assessment of the proposed method, we use APL data. The data sets are described in detail in the following.

##### *S1.1.* BLK *data*

A WGBS data set from whole blood was collected on a cohort of healthy individuals from Sweden. The data set is obtained from the Genome Québec Innovation Centre (Cheung et al., 2017). There are 19 T-cell samples, 13 monocyte samples and 8 B-cell samples in the data. The Illumina HiSeq2000/2500 system was used to sequence the data. Shokoohi et al. (2019) used these data as motivation and to run simulation studies for comparison of DNA methylation analytical pipelines. We follow Shokoohi et al. (2019) and use part of the data as a motivation. We examine the data near the BLK gene on human chromosome 8. The region contains 30,440 CpG sites spanning 2 MB (10,352,236 - 12,422,082) in which approximately 23.39% of the positions have missing information in at least one of the samples and 21% have at least 20 missing values across all samples. An illustration of the data is given in Figure S1. For each cell type in addition to showing the variability in methylation (yellow dots), smooth curves were obtained by: first, aggregating the data (summing read-depths and summing methylation counts over cell type samples at each position), second, imputing the missing values (using a naive method), and third, fitting a smooth curve.

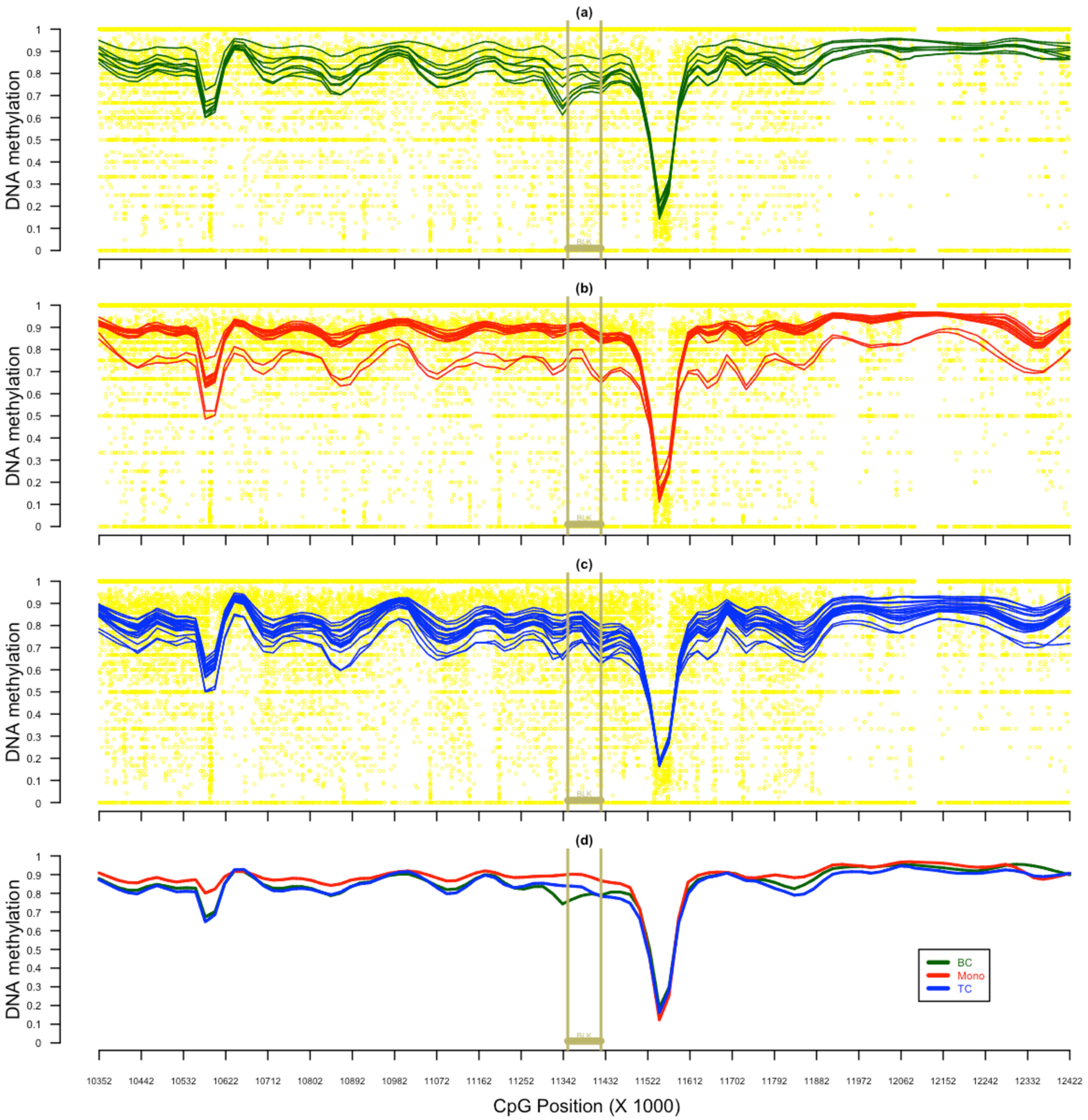


FIGURE S1: *Lowess smoothed profiles of methylation levels near the* BLK *gene for 3 cell types are shown for (a) 8 B-cell samples, (b) 13 monocyte samples and (c) 19 T-cell samples. In each panel methylation proportions at each probe are shown using yellow dots for one sample to illustrate the variability. (d) Smooth aggregated profiles for each cell type are overlaid. The smooth curves were obtained by first aggregating (summing) the read-depths and aggregating methylation counts for each cell type at each position, imputing the missing values, obtaining the methylation levels and finally smoothing using lowess (span = 0.05).*

##### *S1.2.* APL *data*

An RRBS data set is collected from patients with Acute Promyelocytic Leukemia (APL). The data set is publicly available on the Gene Expression Omnibus (accession no. GSE42119) Schoofs et al. (2013). These RRBS data, from the Infinium HumanMethylation450 platform, contain 18 APL patients, and 16 control samples. The control samples include 8 bone marrow samples from patients in remission, four healthy $CD34^+$ cells and four promyelocytes. These data have been analyzed in several articles including Hebestreit, Dugas and Klein (2013). An illustration of the data on chromosome 17 is given in Figure S2. In addition to methylation levels for each position and sample, we have plotted smooth curves of each sample in both APL and control groups. We also plotted the smooth average of methylation levels for groups using lowess (span = 0.00001).

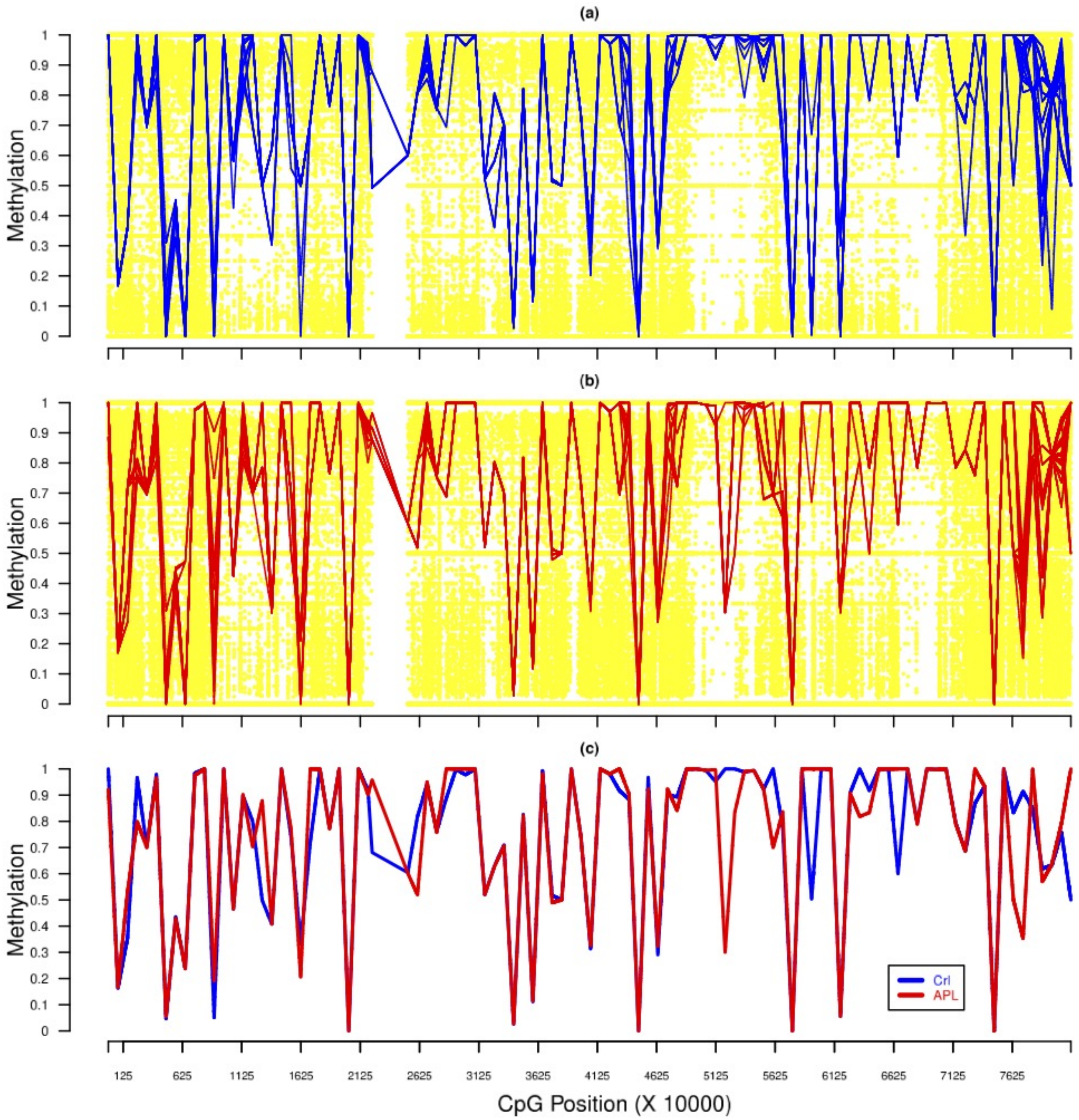


FIGURE S2: *Lowess smoothed profiles of methylation levels for chromosome 17 for two groups are shown for (a) 16 control samples, and (b) 18 APL patients. In each panel methylation proportions at each probe are shown using yellow dots for one sample to illustrate the variability. (c) Smooth aggregated profiles for each group are overlaid. The smooth curves were obtained by first aggregating (summing) the read-depths and aggregating methylation counts for each group at each position, imputing the missing values, obtaining the methylation levels and finally smoothing using lowess (span = 0.00001).*

#### S2. Plots

In this section we provide some preliminary plots to examine the data.

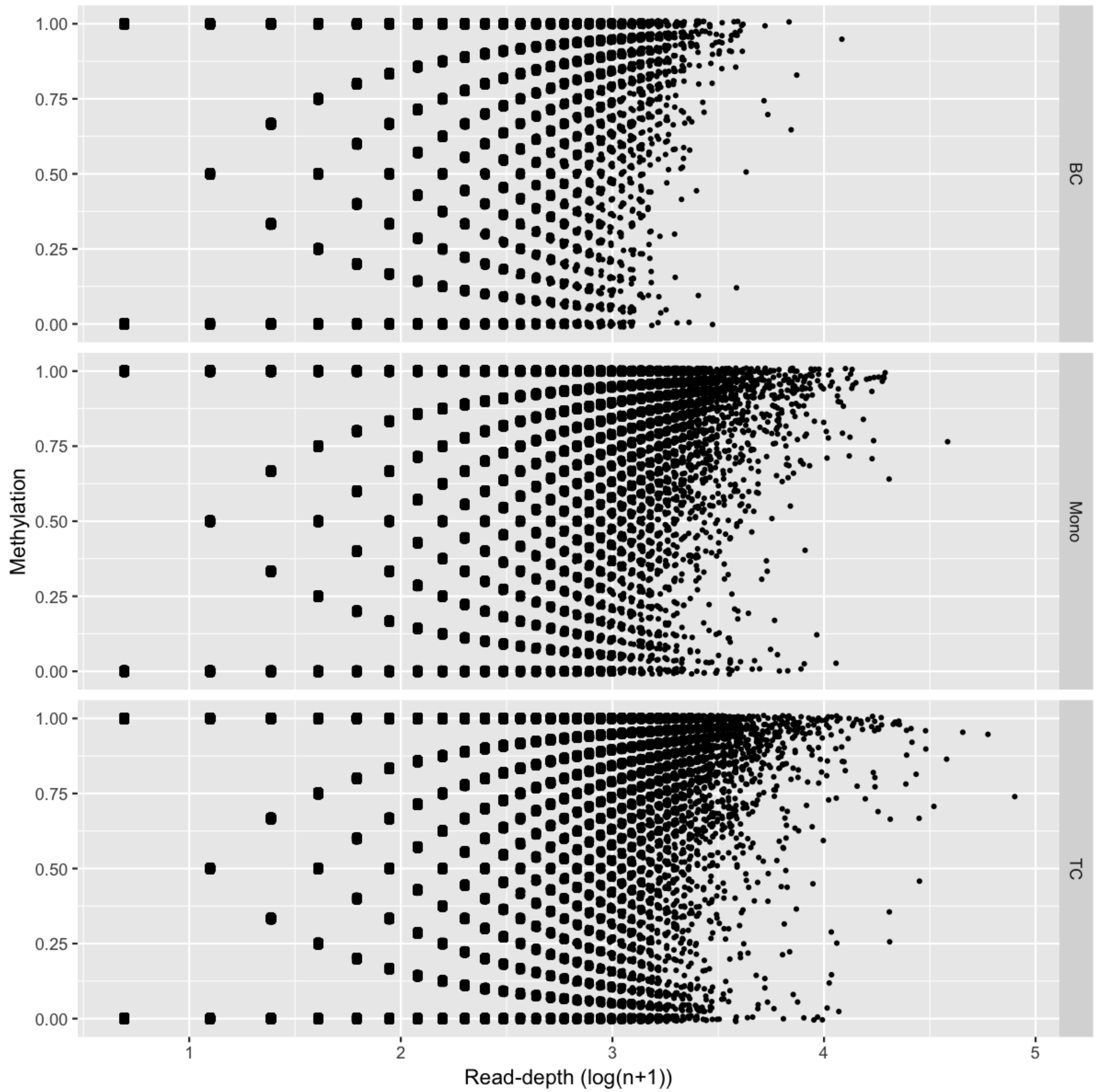


Figure S3: *Methylation level versus* log(*read-depth*+1) *in* BLK *data separated for each cell-type; A small amount of jitter is added to the points.*

#### S3. Details of implementation of the DMCFB method

In this section, we provide several algorithms for implementing our Bayesian functional regression method in the *R-package* DMCFB. The algorithms include the S4-methods, the functions to optimize the number of cores based on available system physical memory, and the functions to find DMCs as well as the functions to plot the results.

---

**Algorithm 1:** Reading methylation data and creating a BSData object in DMCFB

---

**Input:** Files containing samples of genomic positions (in .bed format).
**Input:** A DataFrame containing genomic positions as well as covariates information
Read data: '*object* ← readBismark(<file-paths>, <file-names>, <number of cores>)'.
Assign covariates: 'colData(*object*)← DataFrame(.)'.
**Output:** A BSData object

---

**Algorithm 2:** Implementation of Bayesian functional regression model in DMCFB

---

**Input:** A BSData object
Initialize: *object*
Initialize: $bwa, bwb, nBurn, nMC, nThin, alpha, svd, nCores, pSize, sfiles$ // optional
$object.new$ ← findDMCFB($object, bwa, bwb, nBurn, nMC, nThin, alpha, sdv, nCores, pSize, sfiles$)
**Output:** A BSDMC object that includes MCMC samples
**Output:** A BSDMC object that includes DMCs

---

**Algorithm 3:** Implementation of Bayesian functional regression model in DMCFB

---

**Input:** A BSDMC object that includes MCMC samples
Initialize: *object*
Initialize: $alpha, sfiles$ // optional
$object.new$ ← findDMCFB($object, alpha, sfiles$)
**Output:** A BSDMC object that includes DMCs

---

**Algorithm 4:** Implementation of plots for DMC identification in `DMCFB`

**Input:** A `BSDMC` object that includes DMCs

Initialize: *object*
Initialize: $region, nSplit, parList$ // optional
Initialize in $parList$: $mfrow, type, main, xlab, ylab, col, ticks, nticks, pch, mgp, oma, mar, cex,$ // optional
Initialize in $parList$: $cex.axis, cex.lab, cex.main, lty, lwd, legend, ablines, legend.adj$ // optional
`plotDMCFB`$(object, region, nSplit, parList)$

**Output:** plots that display DMCs as well as the average methylation for each group

**Algorithm 5:** Details of `findDMCFB` method in `DMCFB`

**Input:** A `BSData` object

**if** {`metadata(object)$`*MClist does not exist*} **then**
  $object \leftarrow$ `.MCMCFB`$(object, bwa, bwb, nBurn, nMC, nThin, sdv, nCores, pSize, sfiles)$
$object \leftarrow$ `.findFB`$(object, alpha)$

**Output:** A `BSDMC` object that includes MCMC samples

**Output:** A `BSDMC` object that includes DMCs

**Algorithm 6:** Optimizing the number of cores based on available physical memory in `.MCMCFB` in `DMCFB`

**Input:** A `BSData` object

$xp \leftarrow start(object)$
$nPos \leftarrow length(xp)$
$maxram \leftarrow (get\_ram())[[1]]/1e + 9$
**if** $nCores$ *is missing* **then**
  **if** $maxram/multicoreWorkers() > 5$ **then**
    $nCores \leftarrow multicoreWorkers()$
  **else**
    $nCores \leftarrow floor(maxram/(5 * nBurn * nThin * pSize * bwa * bwb/10000000))$
    $nCores \leftarrow max(1, min(nCores, multicoreWorkers()))$

**Output:** The optimized number of cores

**Algorithm 7:** Creating batches of genomic positions in `.MCMCFB` function in `DMCFB`

**Input:** A `BSData` object

```
xp ← start(object)
nPos ← length(xp)
maxram ← (get_ram())[[1]]/1e + 9
if ((maxram/nCores) > 10) then
    Pos.list1 ← list()
    if (nPos ≥ (floor(5 * nCores/4) * pSize)) then
        Pos.list1 ← list(xp = xp)
    else
        Pos.list1 ← split(xp, ceiling(seq_along(xp)/(nCores * 2 * pSize)))
        aa ← length(Pos.list1)
        if (length(Pos.list1[[aa]]) < (floor(nCores/4) * pSize)) then
            Pos.list1[[aa − 1]] ← c(Pos.list1[[aa − 1]], Pos.list1[[aa]])
            Pos.list1 ← Pos.list1[−aa]
else
    Pos.list1 ← list()
    if (nPos <= (floor(5 * nCores/4) * pSize)) then
        Pos.list1 ← list(xp = xp)
    else
        Pos.list1 ← split(xp, ceiling(seq_along(xp)/(nCores * pSize)))
        aa ← length(Pos.list1)
        if (length(Pos.list1[[aa]]) < (floor(nCores/4) * pSize)) then
            Pos.list1[[aa − 1]] ← c(Pos.list1[[aa − 1]], Pos.list1[[aa]])
            Pos.list1 ← Pos.list1[−aa]
```

**Output:** A list of batches that include genomic partitions

**Algorithm 8:** Creating formula in `.MCMCFB` function in `DMCFB`

**Input:** A `BSData` object

```
formula ← names(colData(object))
```

**Output:** A formula

**Algorithm 9:** Collecting MCMC samples in `.MCMCFB` function in `DMCFB`

```
Input: A list including BSData objects
for {i ← list of objects including BSData regions} in parallel do
    Y0 ← methylation reads; M0 ← read-depths
    nits ← nBurn + nMC ∗ nThin
    G ← # of levels in grouping variable
    L ← # of positions in data, i.e. length(xp)
    nz ← #of covariates + 2
    Xns1 ← ns(xp, df = bwa); Xns2 ← ns(xp, df = bwb)
    Subject ← rep(seq_len(nseq), each = L)
    Beta.old ← matrix(0, nrow = nseq, ncol = bwb)
    Xg ← rep(1, nseq)%x%Xns1; Xb ← rep(1, nseq)%x%Xns2
    Xd.mat ← matrix of all splines
    Xe ← matrix of 1s, logarithm of read-depths and other existing covariates
    data.all ← cbind(as.vector(data.matrix(Y0)), as.vector(data.matrix(M0)))
    Beta.mean ← as.vector(Xns2% ∗ %t(Beta.old))
    GammaDeltaEta.mean ← Xgde% ∗ %GammaDeltaEta.old
    BPrec ← diag(rep(1/sdv, bwb))
    GDEPrec ← diag(rep(1/sdv, bwa ∗ G + nz))
    for myiter ← 1 : nits do
        gdata ← data.frame(Y = data.all[, 1], m = data.all[, 2], Beta.mean, Xgde)
        fitg ← speedglm(cbind(Y, m − Y) ∼ offset(Beta.mean) − 1 + Xgde, data = gdata, family =
          binomial(logit))
        mle.Sig ← summary(fitg)$cov.unscaled
        mle.Prec ← ginv(mle.Sig)
        Post.Cov ← ginv(mle.Prec + GDEPrec)
        Post.mean ← Post.Cov% ∗ %(mle.Prec% ∗ %coef(fitg))
        GammaDeltaEta.old ← mvrnorm(1, mu = Post.mean, Sigma = Post.Cov)
        GammaDeltaEta.mean ← Xgde% ∗ %GammaDeltaEta.old
        for (i ← 1 : nseq) do
            Y ← Y0[, i]; m ← M0[, i]
            ivec ← Subject == i
            idata ← data.frame(Y = Y, m = m, GammaDeltaEta.mean = GammaDeltaEta.mean[ivec], Xns2)
            fiti ← speedglm(cbind(Y, m − Y) ∼ offset(GammaDeltaEta.mean) − 1 + Xns2, data =
              idata, family = binomial(logit))
            mle.Sig ← summary(fiti)$cov.unscaled
            mle.Prec ← ginv(mle.Sig)
            Post.Cov ← ginv(mle.Prec + BPrec)
            Post.mean ← Post.Cov% ∗ %(mle.Prec% ∗ %coef(fiti))
            Beta.old[i, ] ← mvrnorm(1, mu = Post.mean, Sigma = Post.Cov)
        Beta.mean ← as.vector(Xns2% ∗ %t(Beta.old))
        fit0 ← glm(cbind(data.all[, 1], data.all[, 2] − data.all[, 1]) ∼
          offset(GammaDeltaEta.mean + Beta.mean) − 1, family =' binomial')
        if (myiter > nBurn&(myiter%%nThin == 0)) then
            ico ← ico + 1
            Beta.store[ico, , ] ← Beta.old
            GammaDeltaEta.store[ico, ] ← GammaDeltaEta.old
Output: A MCMC list
```

#### S4. List of Selected Methods for Comparison

The chosen methods are presented in Table S1. The overall performance of `DSS-Single`, for instance in terms of accuracy, was close to that of `DSS`, although different for individual DMRs and non-DMRs, thus its results are omitted. For `bumphunter`, all positions, even those with missing values, were included in the analysis to improve the performance. In `bsseq`, we have removed positions where all the samples in one group were missing at that position. In `HMM-Fisher`, distances between CpGs were set to 1 to avoid crashing. See Table S1 for the remaining modifications.

We were not able to include some methods in simulation studies such as `MethPipe` (Song et al., 2013), `methylPipe` (Kishore et al., 2015), `methylKit` (Akalin et al., 2012), and `ComMet` (Saito and Mituyama, 2015)) since these tools require specific file formats, or we were unable to implement them. In any case, these methods have been reported to be inferior to some of the methods that were used for comparison in this study (Klein and Hebestreit, 2015; Wu et al., 2015; Yu and Sun, 2016).

Table S1

*List of DNA methylation analytical tools with their features and capabilities, and the changes made to increase their efficiency*

| Package | Model | Platform | Data | r.NA | filter | Clustering | Smoothing | Parallel | Notes |
|---|---|---|---|---|---|---|---|---|---|
| `BiSeq` | beta regression | BSS | bed | ✓ | ✓ | ✓ | ✓ | partly | The default “perc.samples” is changed from 4/5 to 1/2. |
| `bsseq` | binomial & linear | BSS | bed | ✓ |  | ✓ | ✓ | ✓ | “abs(meanDiff)” is changed to 0.1. |
| `bumphunter` | linear mixed model | BSS | reads | ✓ | ✓ | ✓ | ✓ | ✓ | “cutoff” is changed to 0.05. No position was removed. |
| `DMCHMM` | HMM | BSS | reads bed |  |  | ✓ | ✓ | ✓ | — |
| `DMRcaller` | Kernel smoothing | BSS | custom | ✓ | ✓ | ✓ | ✓ | ✓ | “minProportionDifference” is changed from 0.4 to 0.1. |
| `DSS` | Bayesian hierarchical | BSS RNA | reads | ✓ | ✓ |  |  |  | “p.threshold” is set to 0.05. |
| `DSS-Single` | Bayesian hierarchical | BSS RNA | reads | ✓ | ✓ |  | ✓ |  | “p.threshold” is set to 0.05. |
| `HMM-DM` | HMM | BSS | reads | ✓ | ✓ |  | ✓ |  | “meanDiff.cut”, “max.distance” an[illegible] “max.EM” are changed from 0.3, 1[illegible] and 1 to 0.1, 1000 and 300, respecti[illegible] |
| `HHM-Fisher` | HMM | BSS | reads |  |  |  | ✓ |  | Distances between CpGs are set to [illegible] to avoid crashing. |
| `WFMM` | Wavelet-based functional mixed model |  | custon | ✓ | ✓ |  | ✓ | ✓ | Only Windows; no `R` package; difficult to implement parallel comp[illegible] |

### S5. Simulation Study from the Main Paper

#### *S5.1. Simulation Set-ups*

In the following table we present the simulation set-ups for the first and second scenarios in Tables S2 and S3, respectively.

Table S2
*Simulation setup parameters for the first scenario*

| DMR | Chromosome | Start position | End position | Effective size | Length |
|---|---|---|---|---|---|
| DMR1 | 8 | 10431839 | 10511940 | 0.1 | 1620 |
| DMR2 | 8 | 10671903 | 10751815 | 0.2 | 1094 |
| DMR3 | 8 | 10911977 | 10991817 | 0.3 | 1205 |
| DMR4 | 8 | 11231950 | 11311987 | 0.4 | 1267 |
| DMR5 | 8 | 11631990 | 11641956 | 0.4 | 126 |
| DMR6 | 8 | 11671950 | 11681894 | 0.3 | 102 |
| DMR7 | 8 | 11714976 | 11725000 | 0.2 | 223 |
| DMR8 | 8 | 11783926 | 11793886 | 0.1 | 224 |
| DMR9 | 8 | 11870570 | 11924576 | 0.05 | 924 |
| DMR10 | 8 | 12084164 | 12191969 | 0.02 | 433 |

Table S3
*Simulation setup parameters for the second scenario*

| DMR | Chromosome | Start position | End position | Effective size | Length |
|---|---|---|---|---|---|
| DMR1 | 8 | 10431839 | 10441997 | 0.1 | 172 |
| DMR2 | 8 | 10701967 | 10729553 | 0.2 | 363 |
| DMR3 | 8 | 10951995 | 10971974 | 0.3 | 274 |
| DMR4 | 8 | 11281992 | 11301965 | 0.4 | 358 |
| DMR5 | 8 | 11584972 | 11629892 | 0.4 | 903 |
| DMR6 | 8 | 11679894 | 11714976 | 0.3 | 643 |
| DMR7 | 8 | 11764964 | 11794972 | 0.2 | 703 |
| DMR8 | 8 | 11874947 | 11924642 | 0.1 | 669 |
| DMR9 | 8 | 12044993 | 12061576 | 0.05 | 307 |
| DMR10 | 8 | 12204790 | 12232000 | 0.02 | 120 |

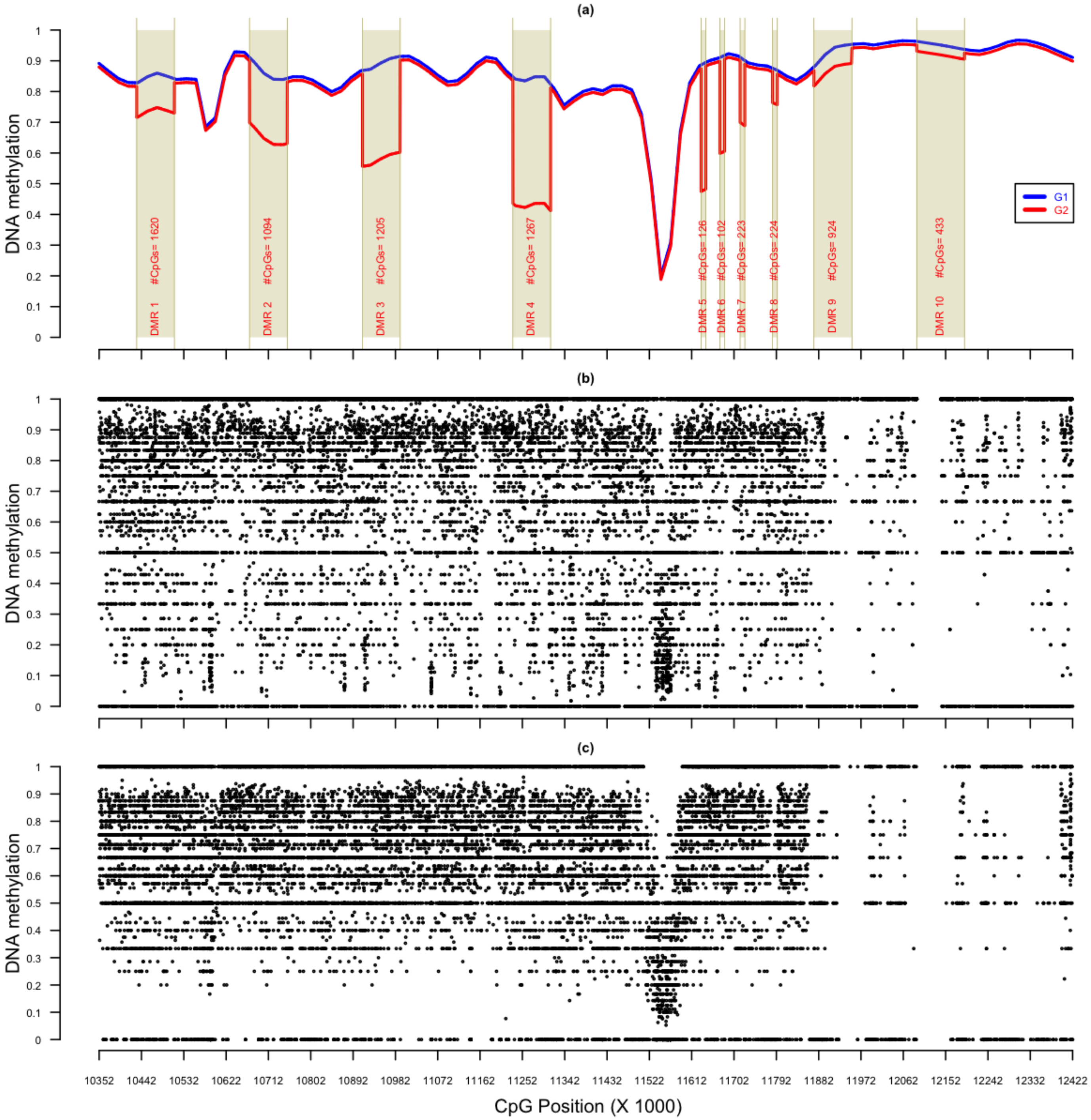


Figure S4: *Methylation profiles illustrating simulation design for the first scenario. (a) The smooth methylation profile (G1), obtained by smoothing aggregated B-cell samples, from which the true mean methylation levels in the simulated data sets are generated. (b) Actual methylation proportions for one B-cell sample. (c) A set of simulated methylation data for the same sample as in (b), with error generated from a normal distribution with standard deviation equal 0.18.*

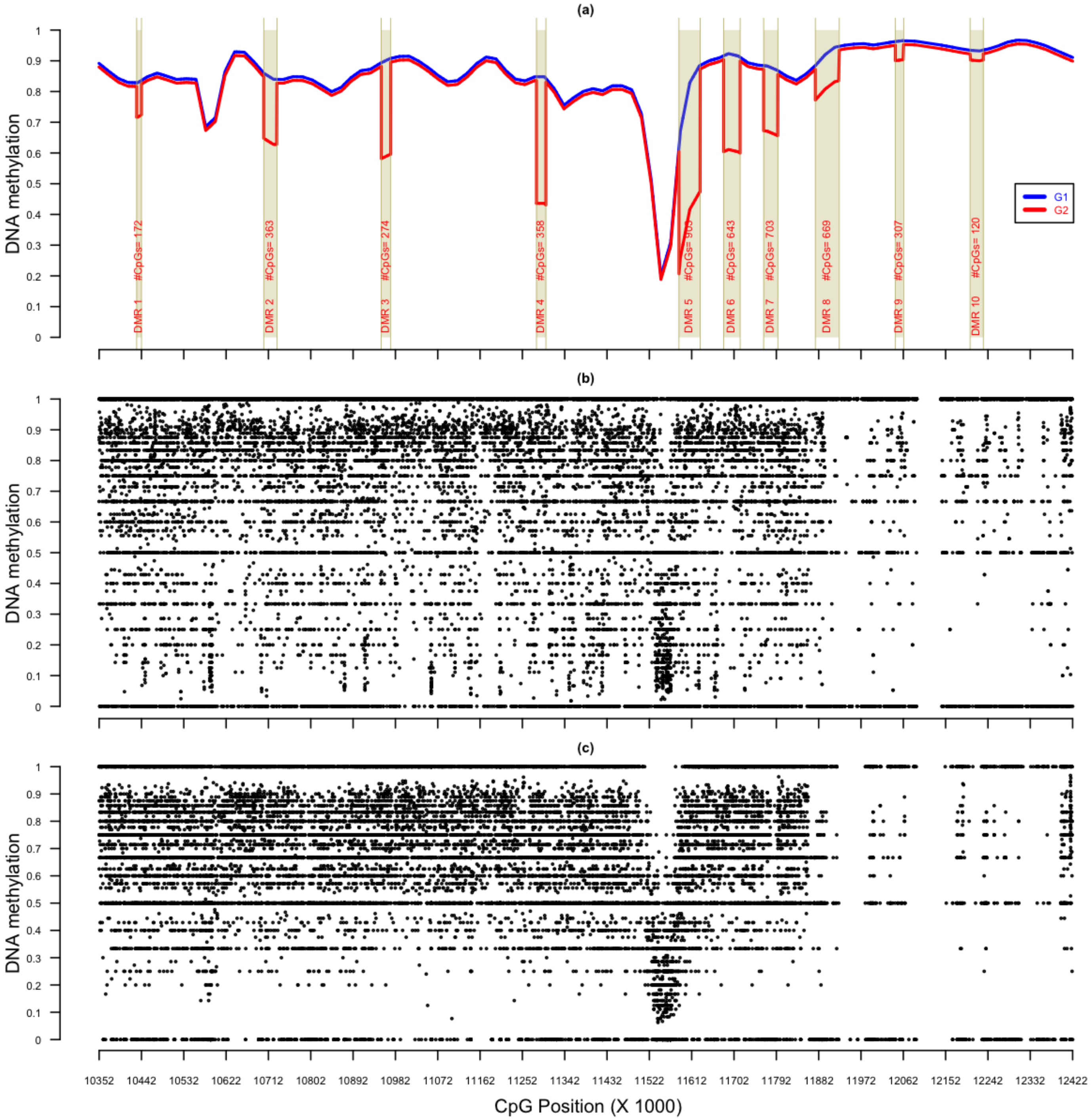


FIGURE S5: *Methylation profiles illustrating simulation design for the second scenario. (a) The smooth methylation profile (G1), obtained by smoothing aggregated B-cell samples, from which the true mean methylation levels in the simulated data sets are generated. (b) Actual methylation proportions for one B-cell sample. (c) A set of simulated methylation data for the same sample as in (b), with error generated from a normal distribution with standard deviation equal 0.18.*

#### *S5.2. Figures and Tables of Simulation Setup Scenario 1, $\sigma = 0.18$*

In this section we provide the results of simulation study for the first scenario when errors are generated from a normal distribution with standard deviation equal to 0.18.

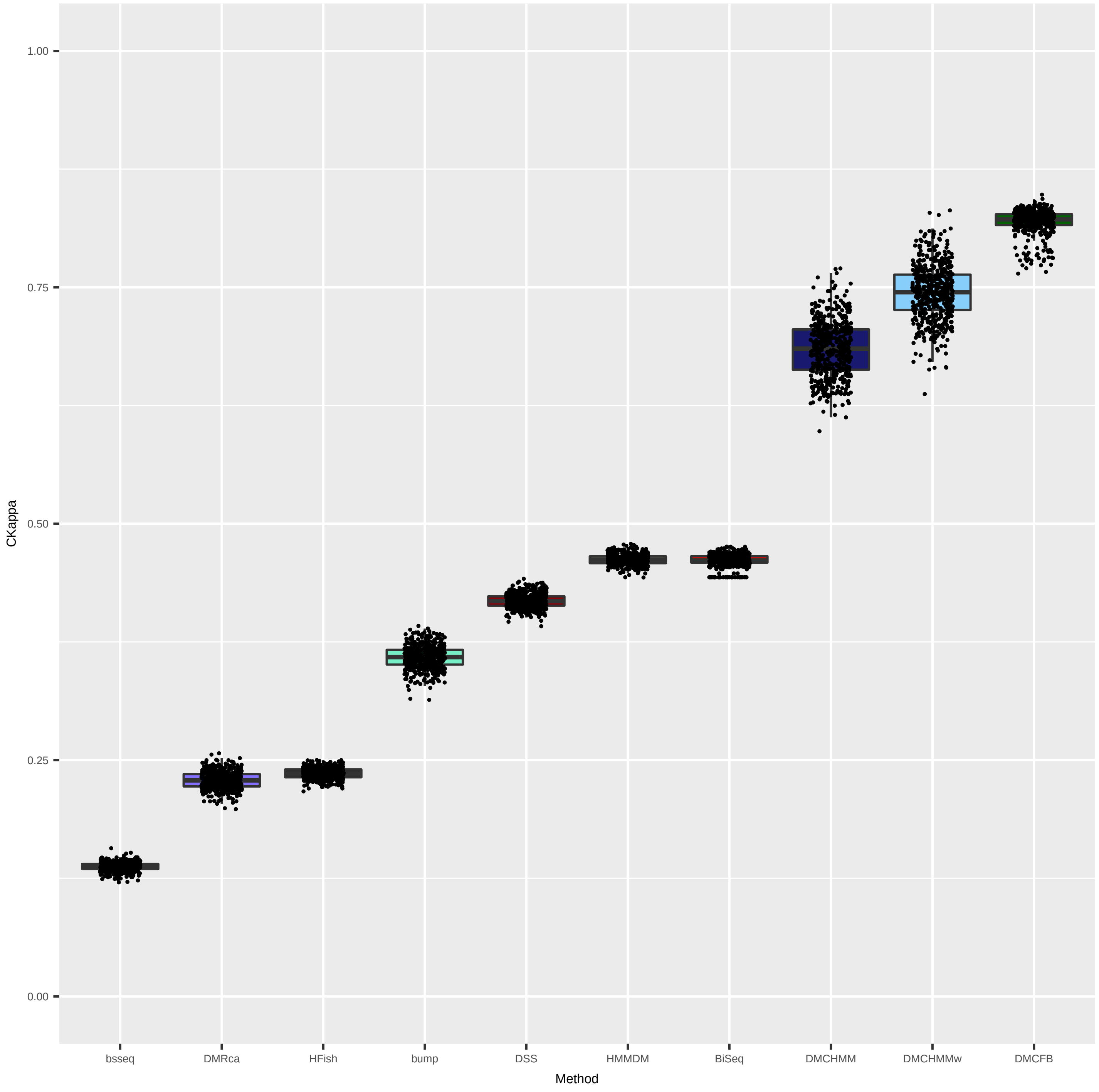


FIGURE S6: *The box-plots of Cohen's Kappa for each method in simulated data for the first scenario; Errors are generated from $N(\mu = 0, \sigma = 0.18)$.*

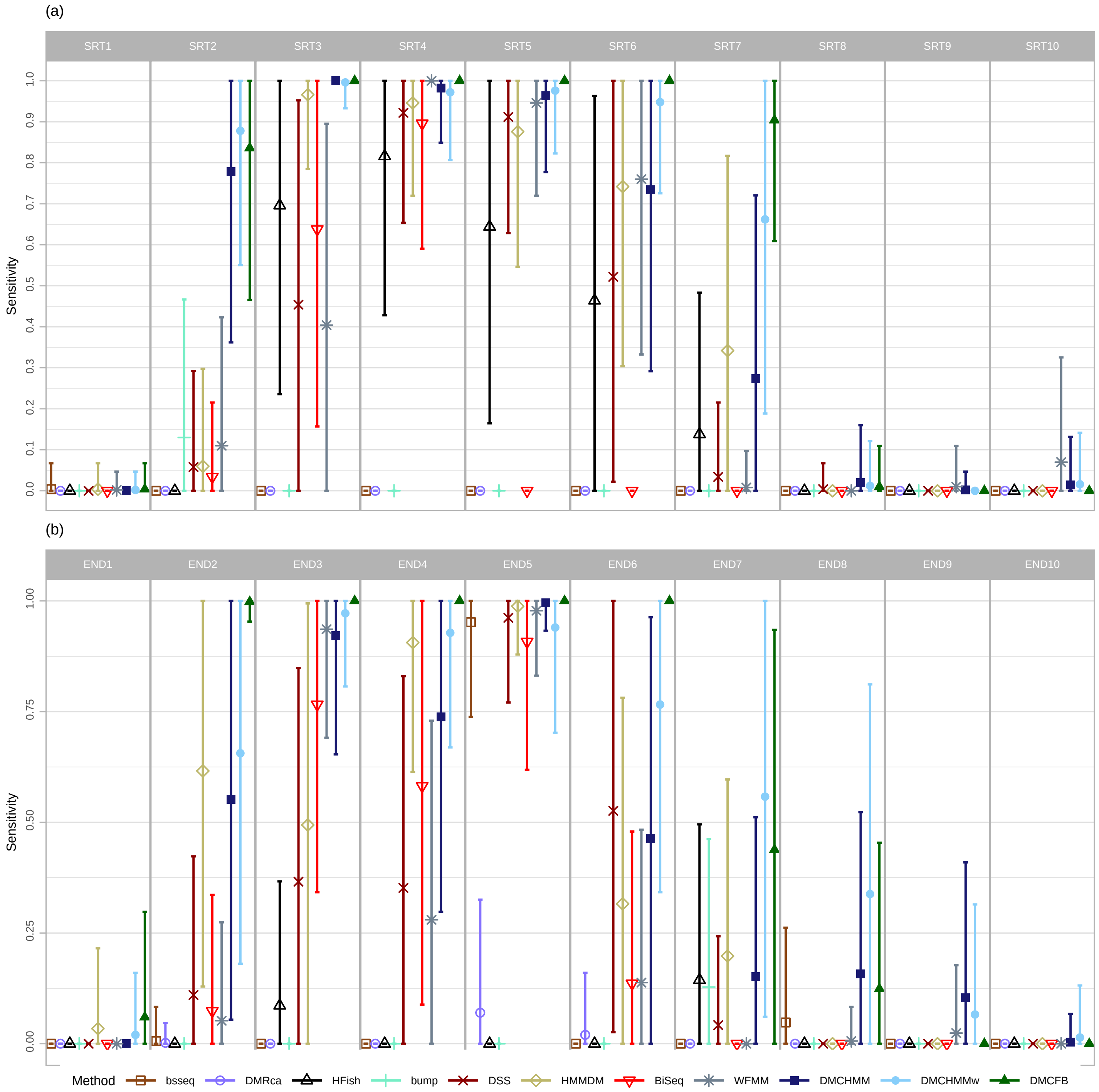


FIGURE S7: *Average number of times that the start (a) and end (b) positions of* DMRs *are identified as* DMCs *for different methods in simulated data for the first scenario; Errors are generated from* $N(\mu = 0, \sigma = 0.18)$.

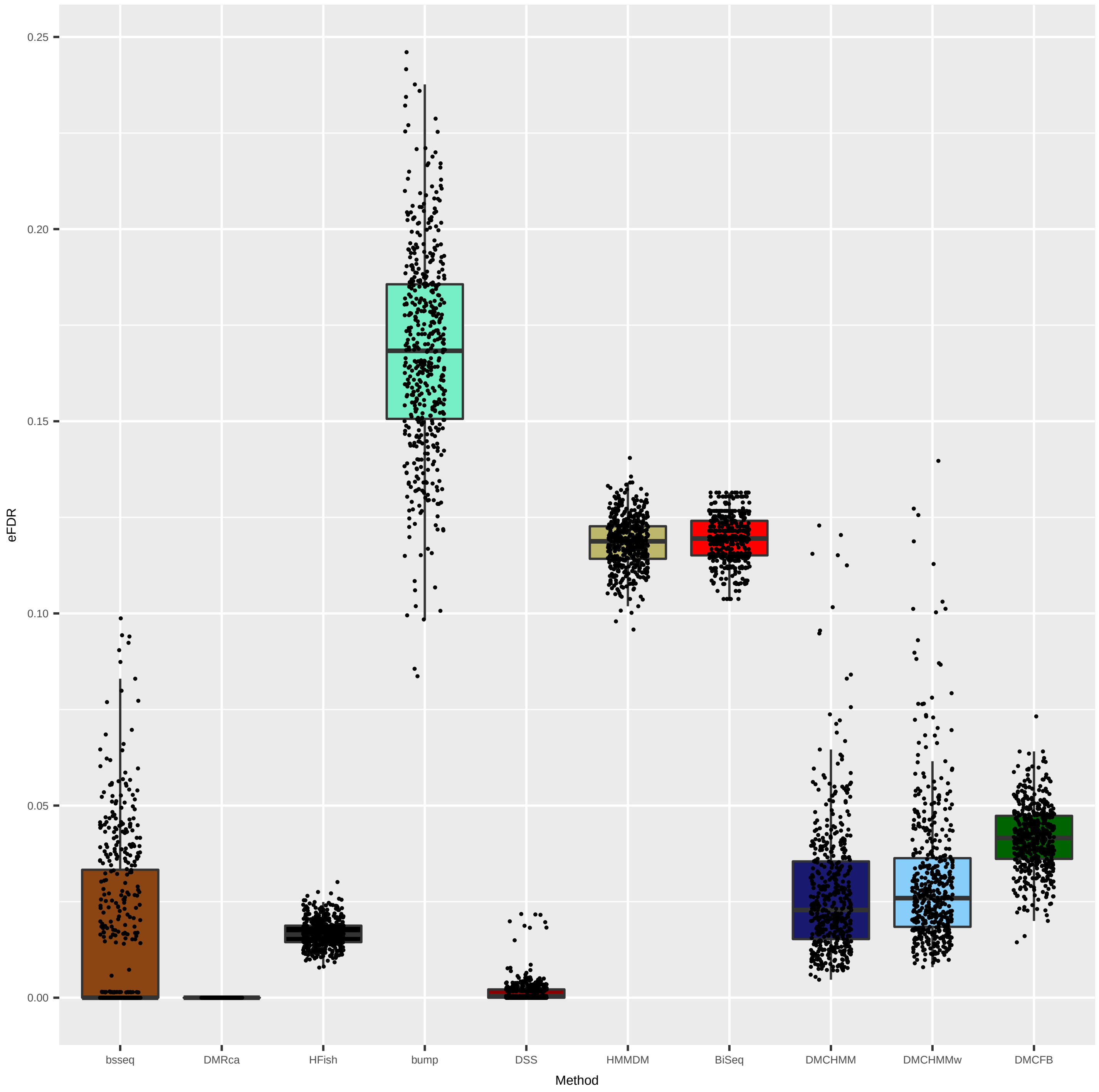


FIGURE S8: *Average empirical FDR for different methods in the first scenario; Errors are generated from* $N(\mu = 0, \sigma = 0.18)$. *The vertical axis is truncated.*

##### *S5.3. Figures and Tables of Simulation Setup Scenario 1, $\sigma = 0.24$*

We have used a second set of simulation scenarios to assess the performance of our proposed method. In this section we present the results of the first scenario in which errors were generated from $N(0, 0.24)$ to add to methylation levels of G1 and G2 curves.

The average overall accuracy (ACC) and average overall modified accuracy (MACC) is plotted in Figure S9 for different methods. Figure S10 shows Cohen's Kappa values. The average 'Sensitivity' and the average '1-Specificity' are illustrated in Figure S11. The proportions of the times the start and the end of DMRs are identified are depicted in Figure S12. Finally, Figure S13 shows the empirical FDR for different methods. In general, we see even better results with a larger noise in the data using `DMCFB`. This observation shows that our method is robust with respect to the variation in the data.

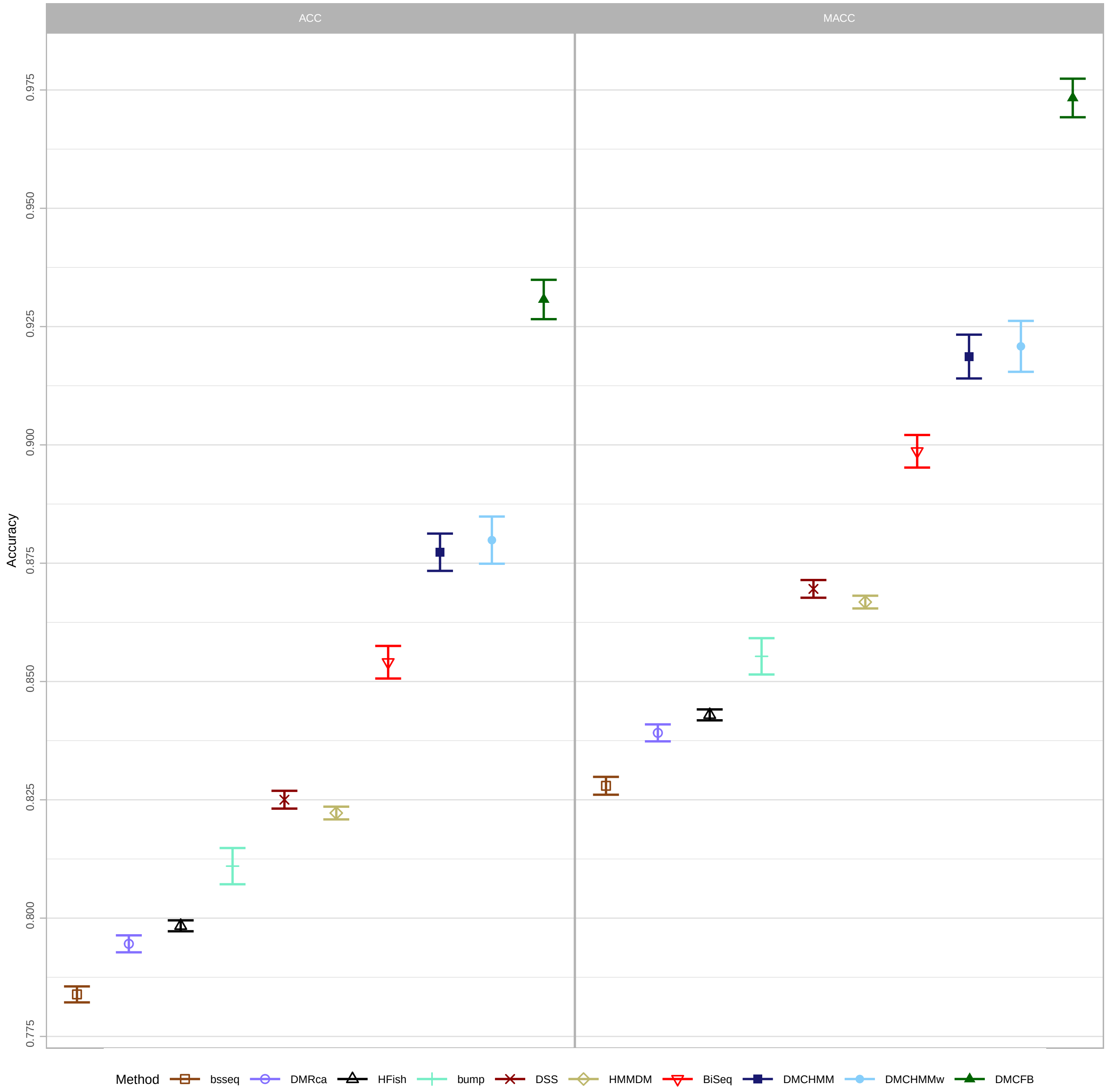


FIGURE S9: *The average overall accuracy (ACC) and average overall modified accuracy (MACC) for different methods in simulated data for the first scenario; Errors are generated from $N(\mu = 0, \sigma = 0.24)$. The vertical axis is truncated.*

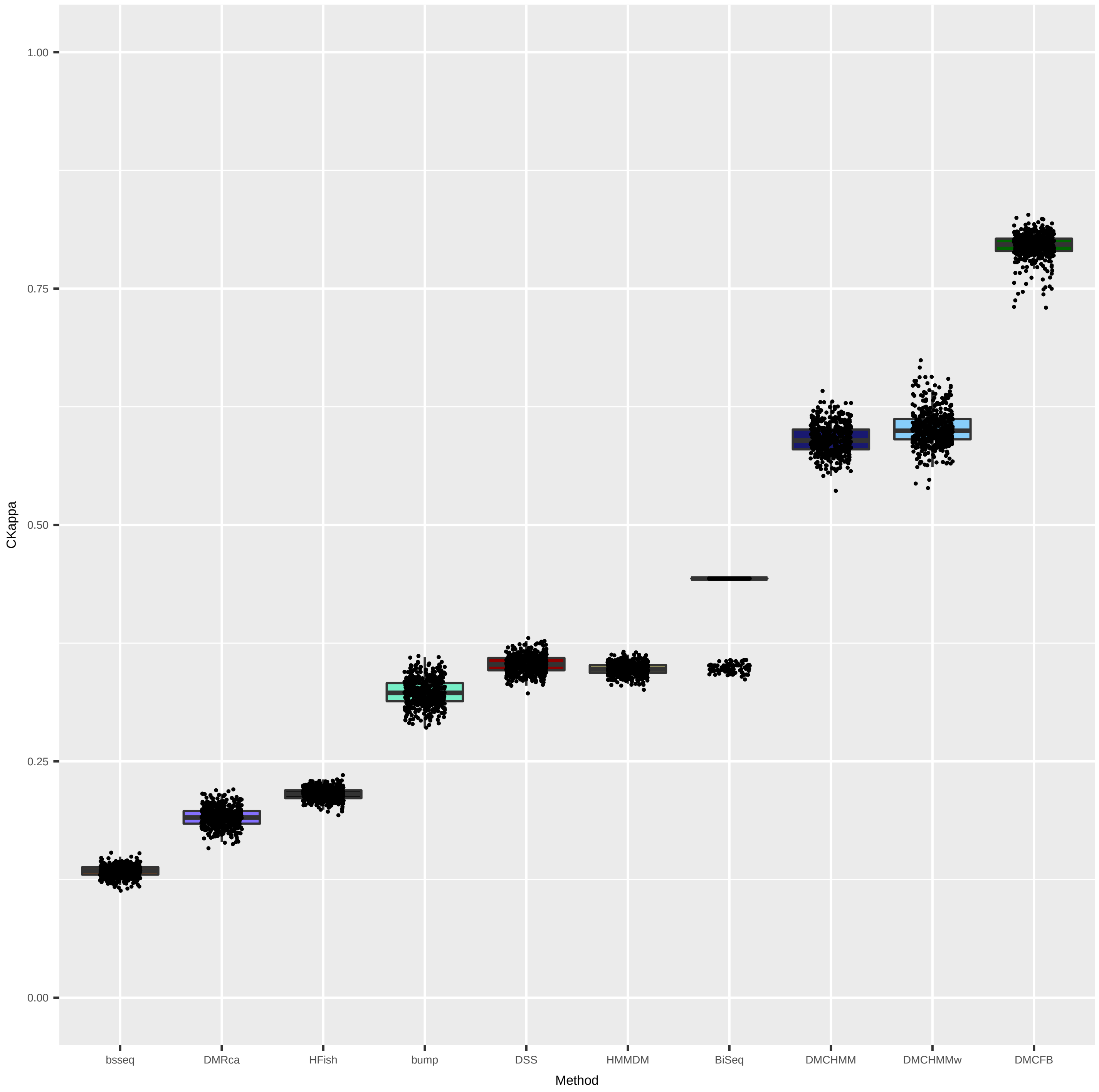


FIGURE S10: *The box-plots of Cohen's Kappa for each method in simulated data for the first scenario; Errors are generated from* $N(\mu = 0, \sigma = 0.24)$. *The vertical axis is truncated.*

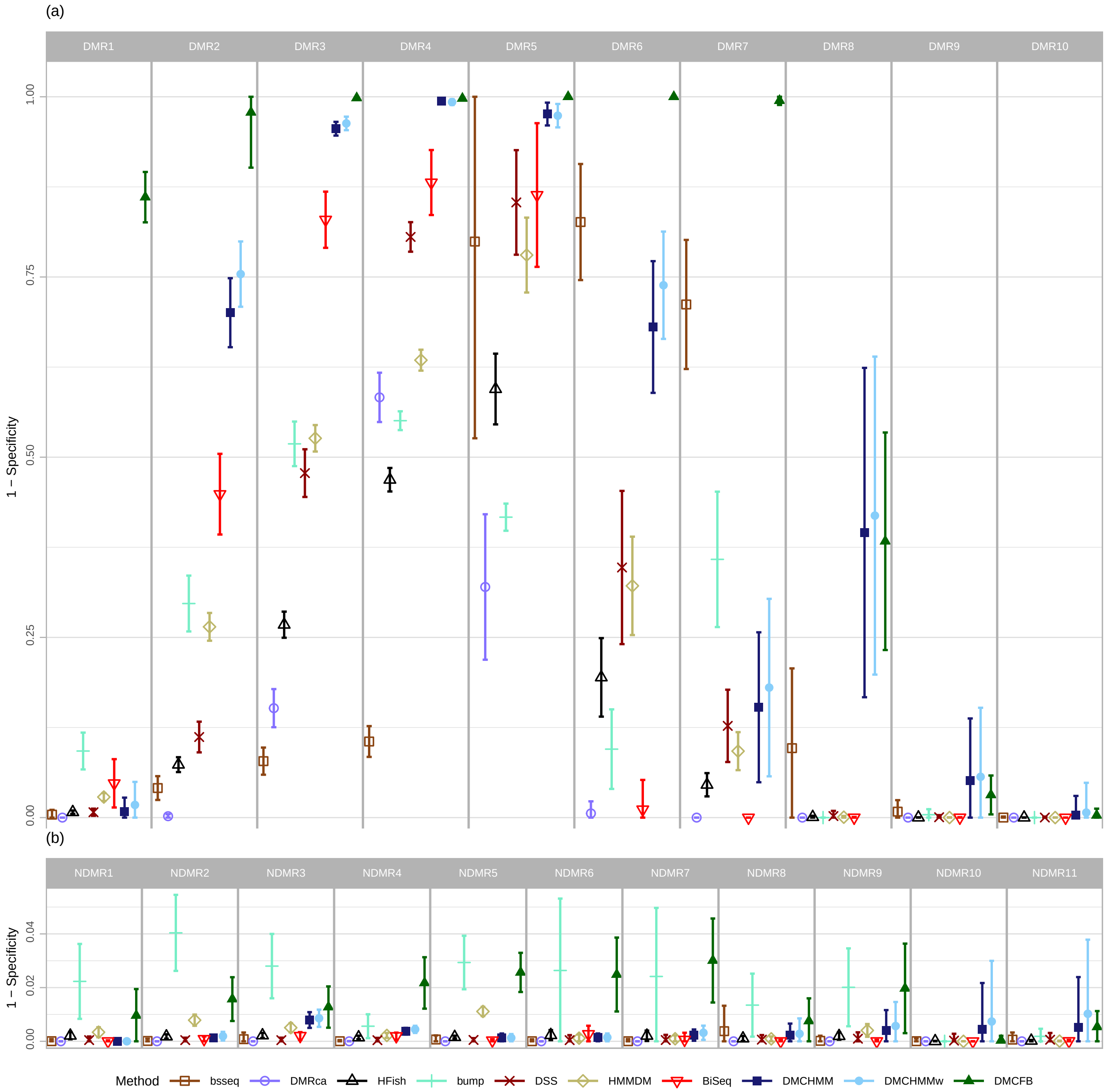


FIGURE S11: *(a) The average proportion of correctly identified* DMCs *(Sensitivity) for each method separated by* DMRs *(sd error bars are added); (b) The average proportion of incorrectly identified* DMCs *(1 - Specificity) for each method separated by* NDMRs *(The axis is truncated) in simulated data for the first scenario; Errors are generated from* $N(\mu = 0, \sigma = 0.24)$.

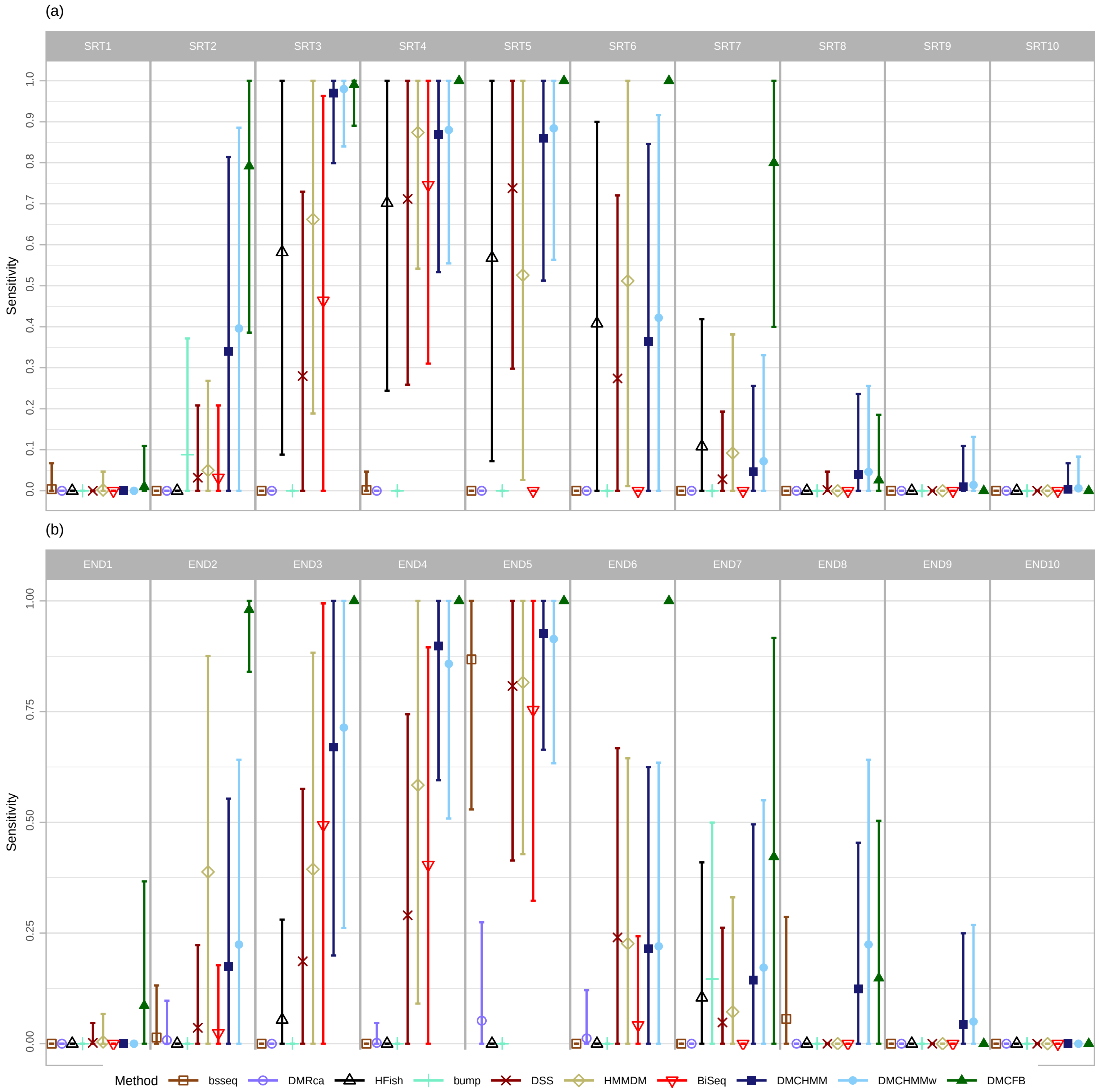


FIGURE S12: *Average number of times that the start (a) and end (b) positions of* DMRs *are identified as* DMCs *for different methods in simulated data for the first scenario; Errors are generated from* $N(\mu = 0, \sigma = 0.24)$.

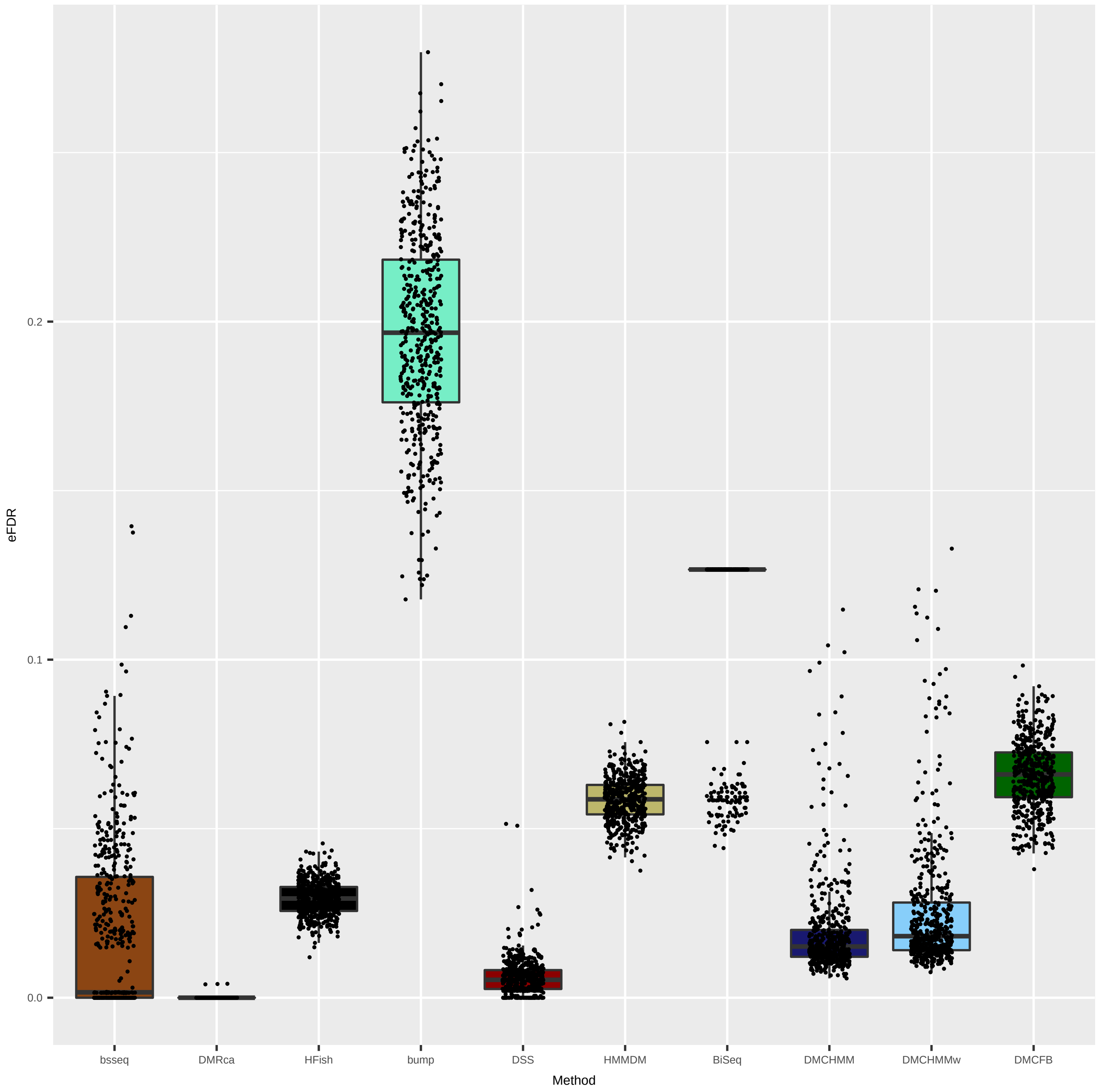


FIGURE S13: *Average empirical FDR for different methods in simulated data for the first scenario; errors are generated from* $N(\mu = 0, \sigma = 0.24)$*. The vertical axis is truncated.*

##### *S5.4. Figures and Tables of Simulation Set-up Scenario 2, $\sigma = 0.18$*

In this section we consider the second scenario given in Table S3 in Section S5.1 with DMRs of different lengths and locations to further assess the performance of our method.

The overall ACC and MACC are depicted in Figure S14. Comparison of methods in terms of the Cohen's Kappa criterion is provided in Figure S15. Figure S16 depicts 'Sensitivity' and '1 - Specificity' (and their standard deviation error-bar) for each method separated by DMRs and non-DMRs, respectively. The proportion of the start and the end of each DMR using different methods are shown in Figure S17. Finally, the empirical FDR for different methods is illustrated in Figure S18. All in all, the results are aligned with what we observed for the first simulation set-up.

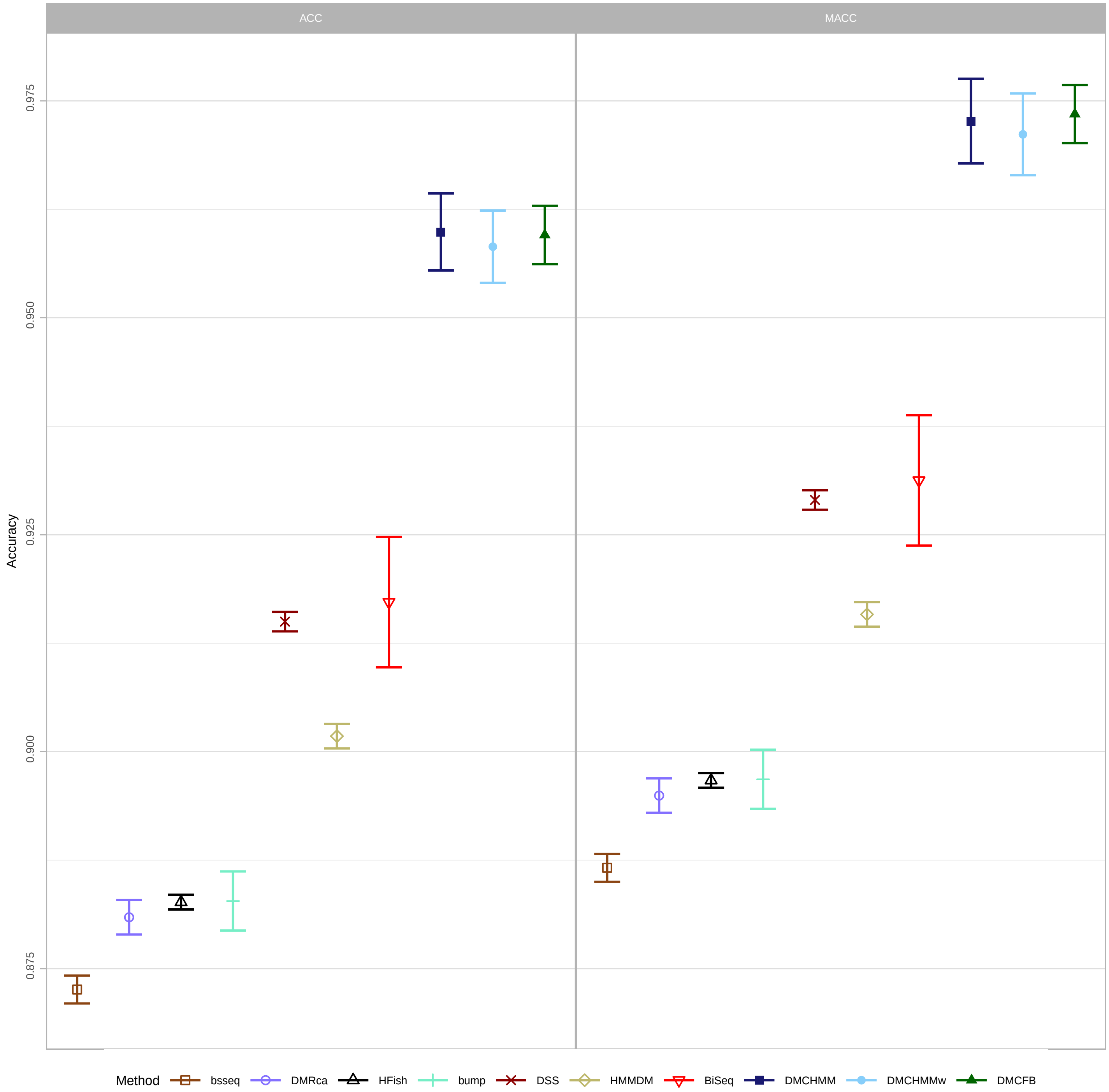


FIGURE S14: *The average overall accuracy (ACC) and average overall modified accuracy (MACC) for different methods in simulated data for the second scenario; Errors are generated from* $N(\mu = 0, \sigma = 0.18)$.

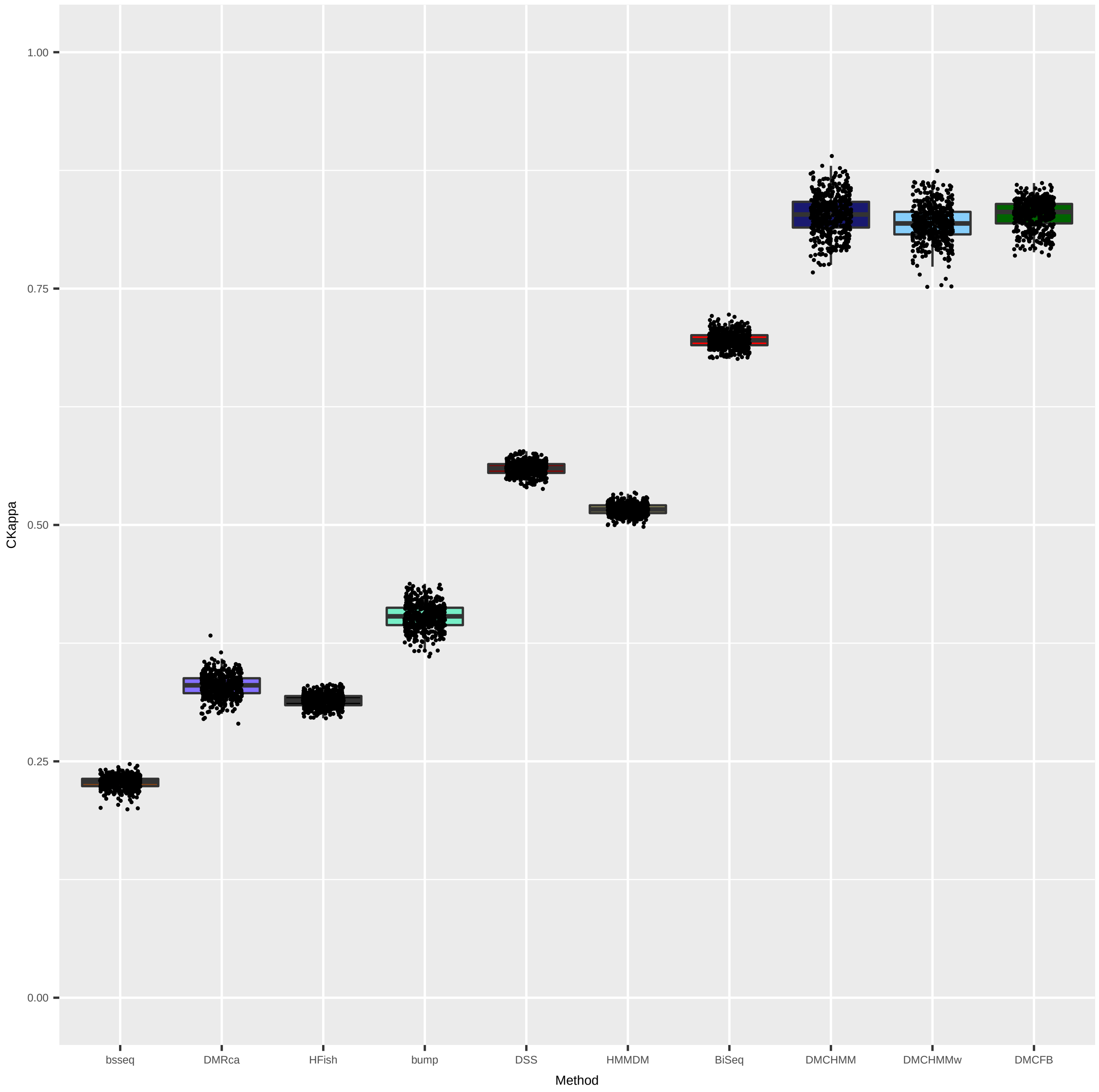


FIGURE S15: *The box-plots of Cohen's Kappa for each method in simulated data for the second scenario; Errors are generated from $N(\mu = 0, \sigma = 0.18)$. The vertical axis is truncated.*

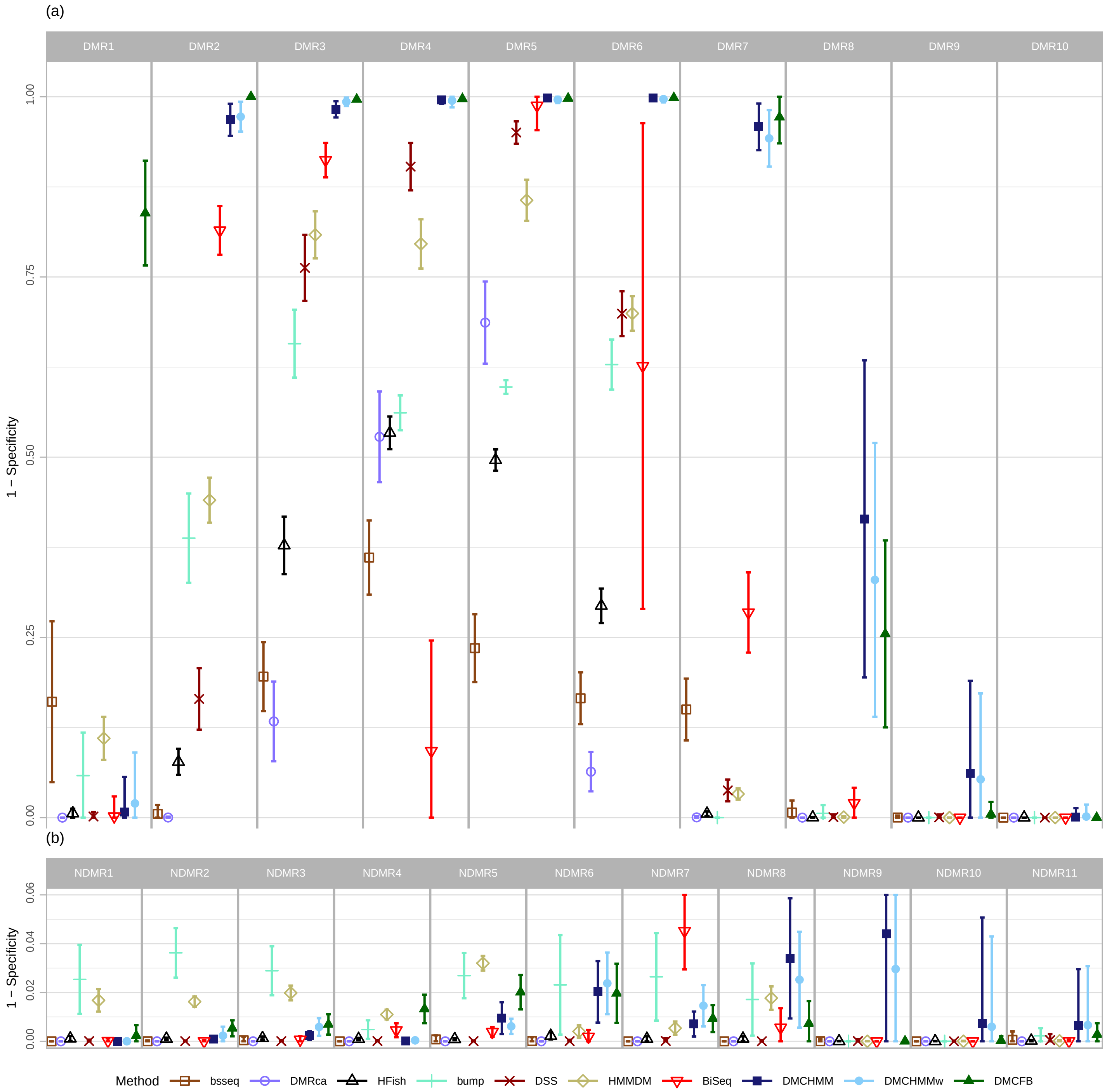


FIGURE S16: *(a) The average proportion of correctly identified* DMCs *(Sensitivity) for each method separated by* DMRs *(sd error bars are added); (b) The average proportion of incorrectly identified* DMCs *(1 - Specificity) for each method separated by* NDMRs *(The axis is truncated) in simulated data for the second scenario; Errors are generated from* $N(\mu = 0, \sigma = 0.18)$.

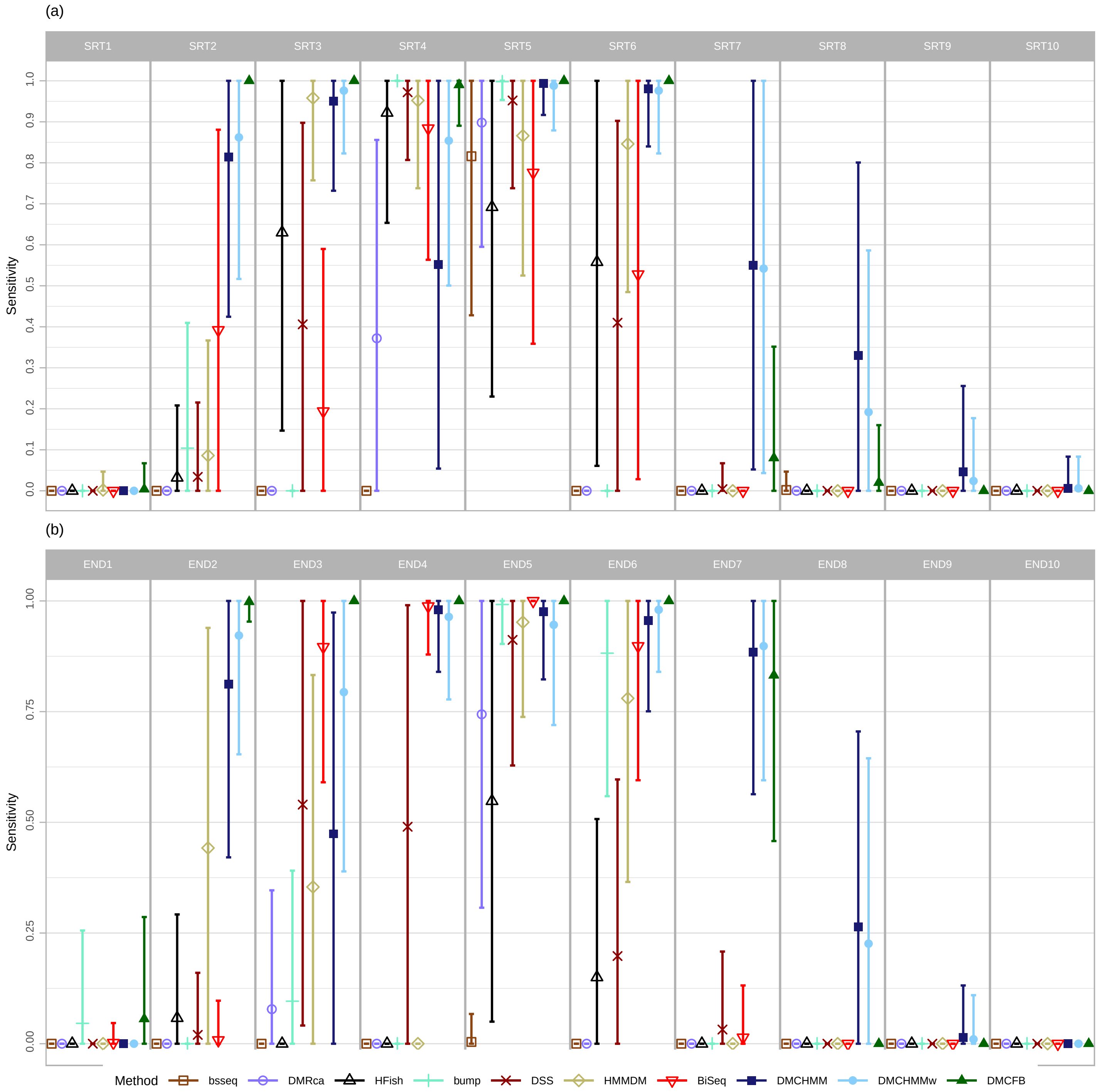


FIGURE S17: *Average number of times that the start (a) and end (b) positions of* DMRs *are identified as* DMCs *for different methods in simulated data for the second scenario; Errors are generated from* $N(\mu = 0, \sigma = 0.18)$.

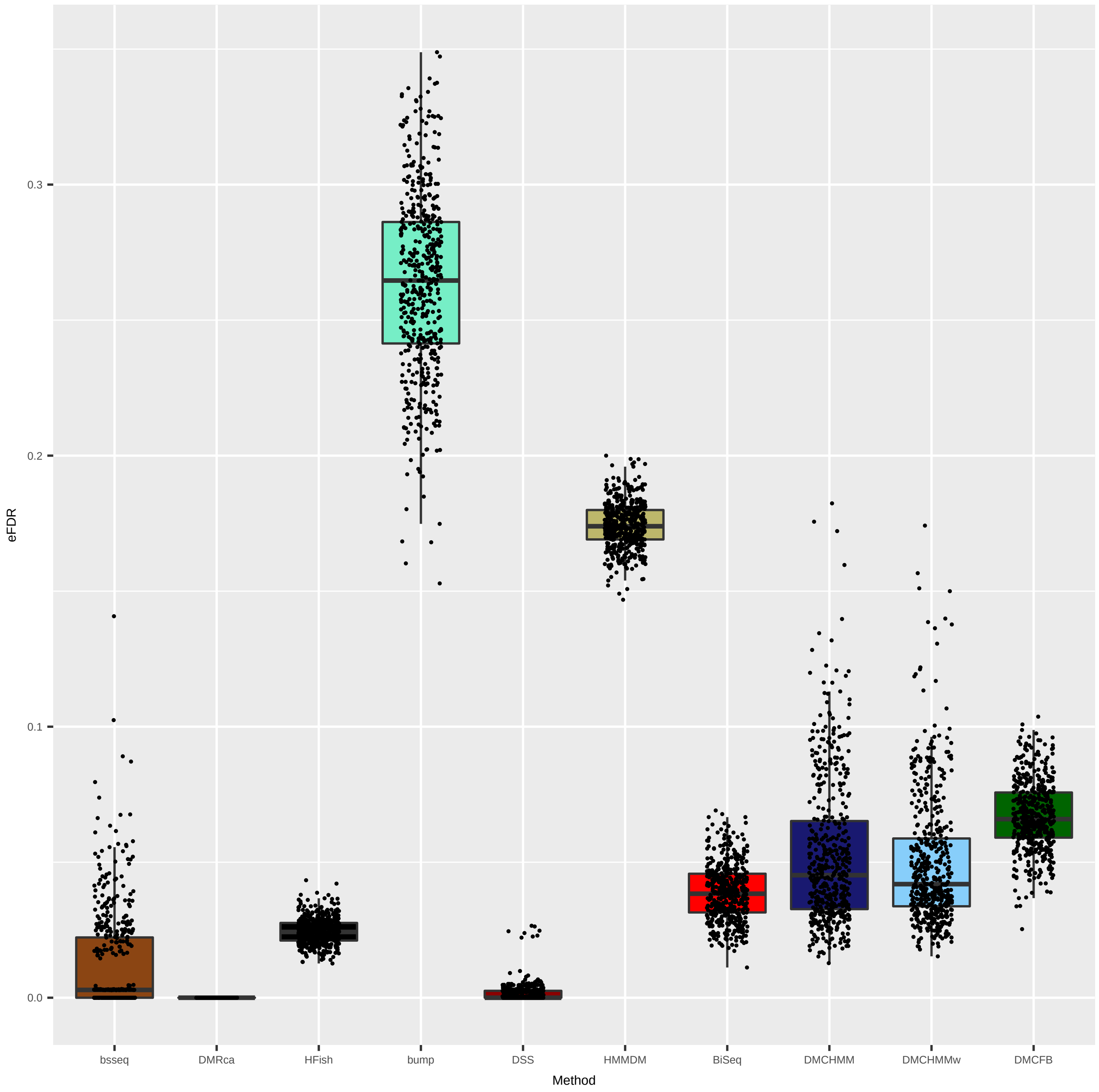


FIGURE S18: *Average empirical FDR for different methods in simulated data for the second scenario; Errors are generated from $N(\mu = 0, \sigma = 0.18)$. The vertical axis is truncated.*

#### S6. Sensitivity Analysis

Sensitivity analyses with respect to the choices of band-width and precision of prior are provided here.

##### S6.1. Sensitivity analysis with respect to band-width; Scenario 1, $\sigma = 0.18$

A set of band-width values in $\{20, 25, \ldots, 60\}$ is chosen for sensitivity analysis. In the following figures we provide the results.

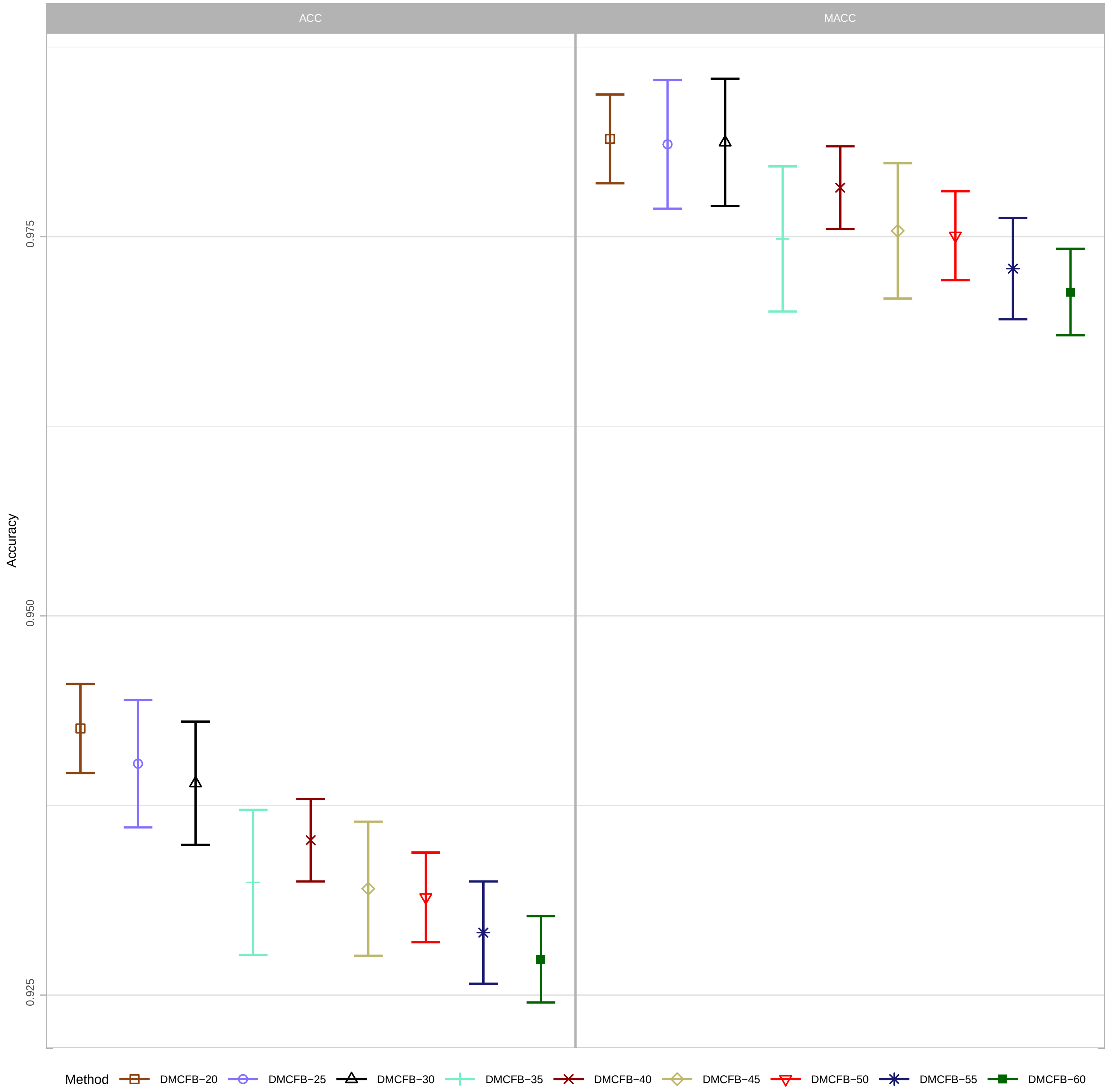


FIGURE S19: *The average overall accuracy (ACC) and average overall modified accuracy (MACC) for each choice of band-width in* `DMCFB` *in simulated data for the first scenario; Errors are generated from* $N(\mu = 0, \sigma = 0.18)$*. The vertical axis is truncated.*

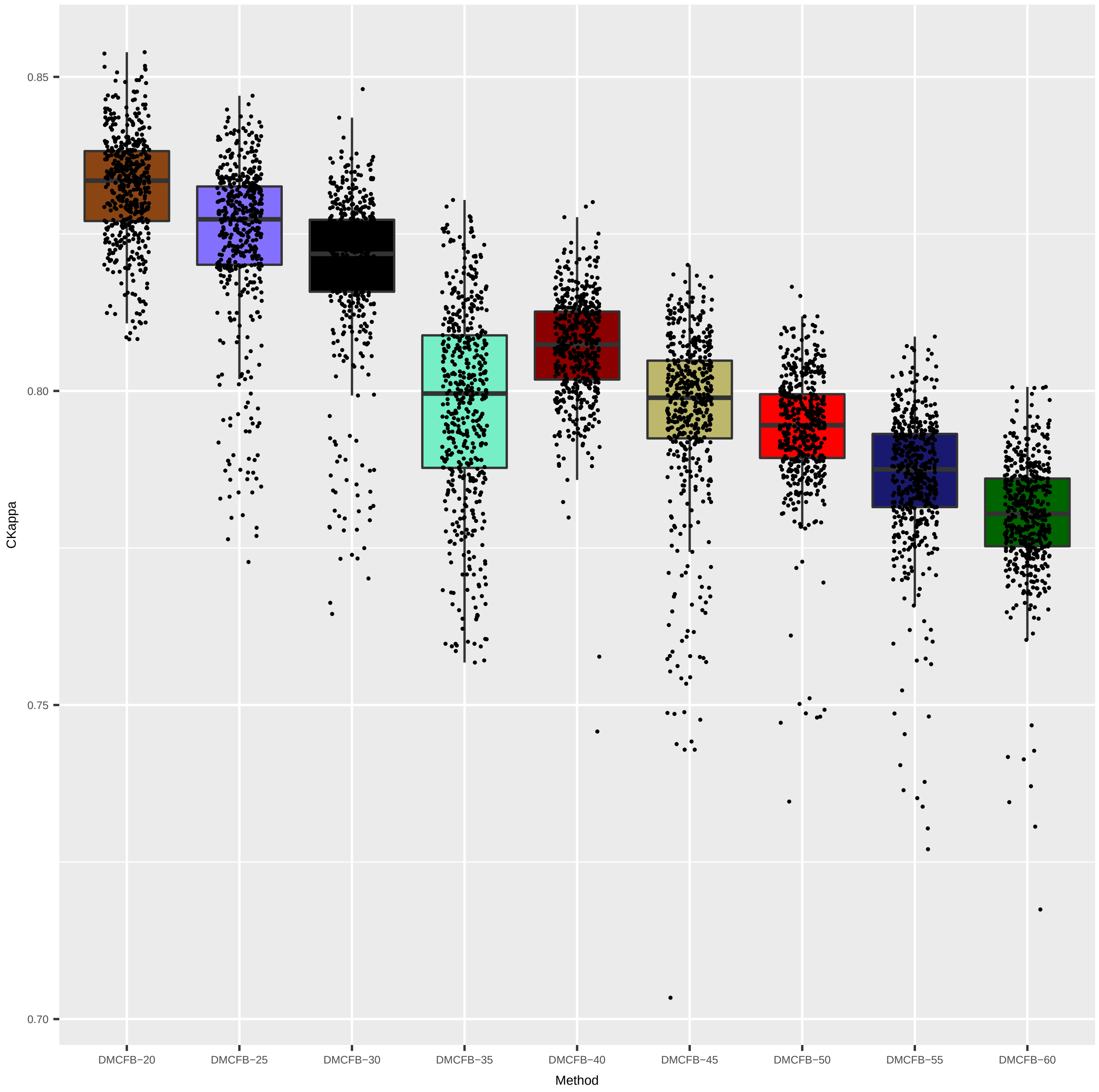


FIGURE S20: *The box-plots of Cohen's Kappa for each choice of band-width in* `DMCFB` *in simulated data for the first scenario; Errors are generated from* $N(\mu = 0, \sigma = 0.18)$. *The vertical axis is truncated.*

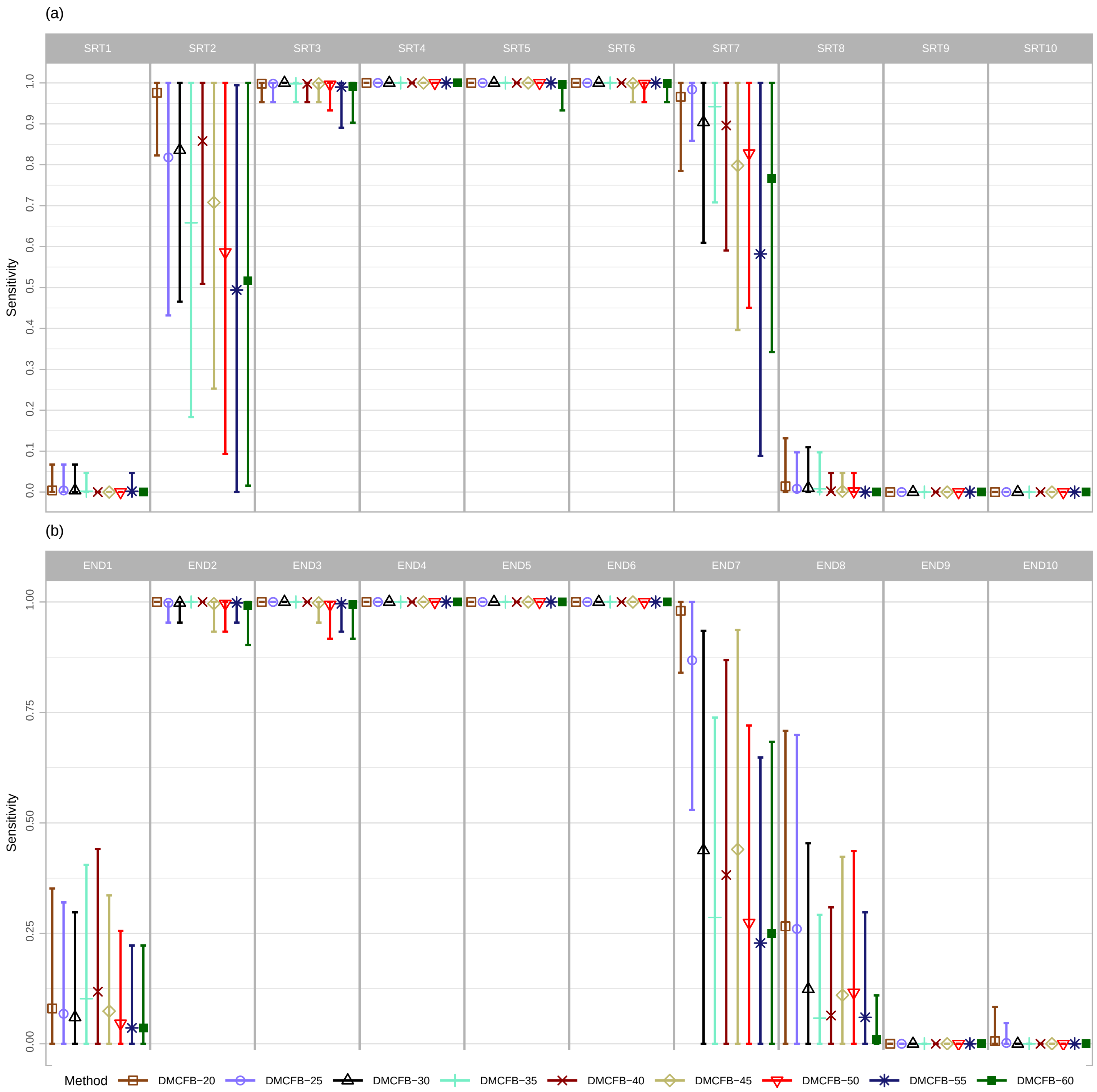


FIGURE S21: *Average number of times that the start (a) and end (b) positions of* DMRs *are identified as* DMCs *for each choice of band-width in* DMCFB *in simulated data for the first scenario; Errors are generated from* $N(\mu = 0, \sigma = 0.18)$.

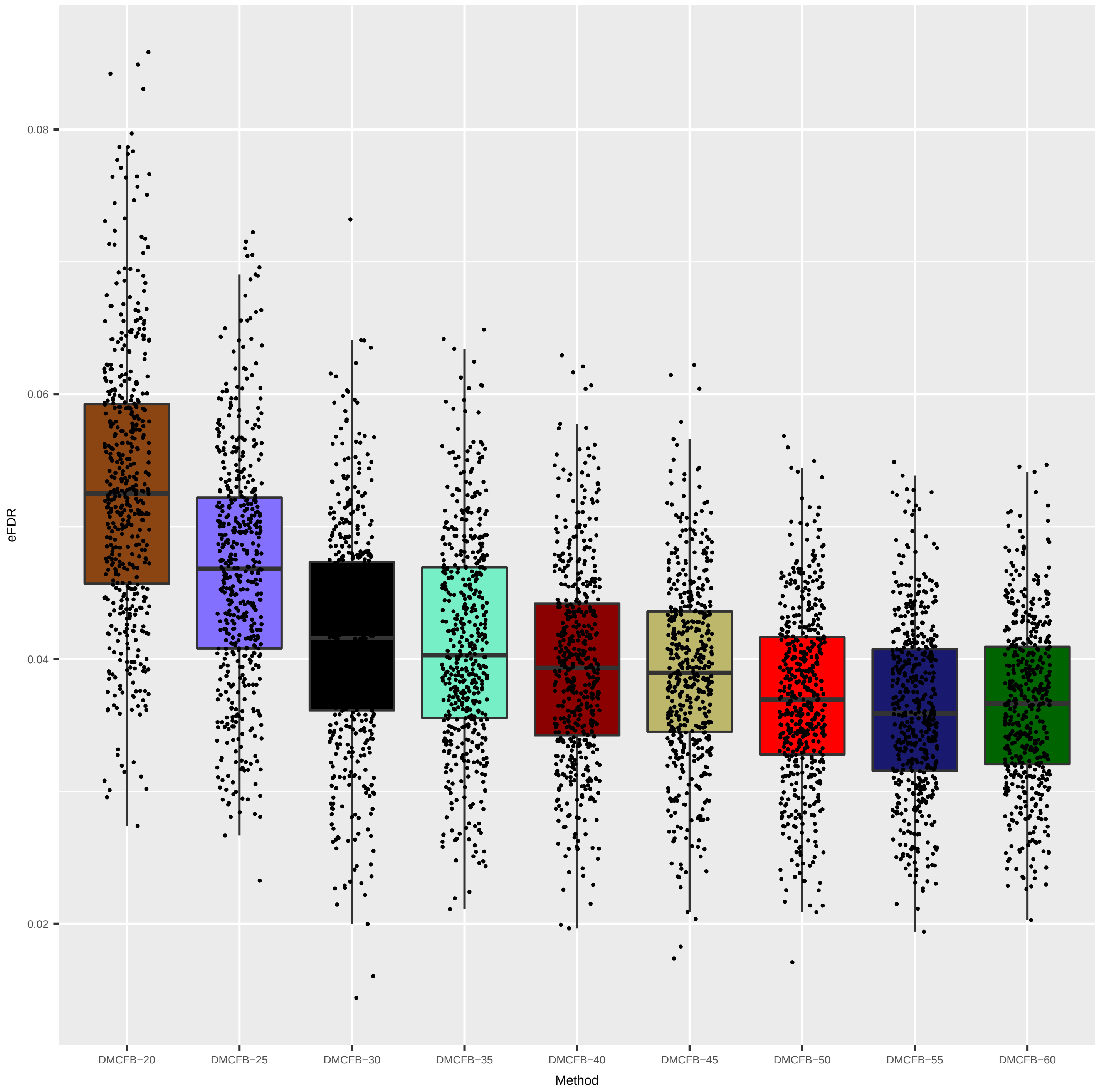


FIGURE S22: *Average empirical FDR for each choice of band-width in* DMCFB *in simulated data for the first scenario; Errors are generated from* $N(\mu = 0, \sigma = 0.18)$. *The vertical axis is truncated.*

##### *S6.2. Sensitivity analysis with respect to band-width; Scenario 2, $\sigma = 0.18$*

A set of band-width values in $\{20, 25, \ldots, 60\}$ is chosen for sensitivity analysis. In the following figures we provide the results.

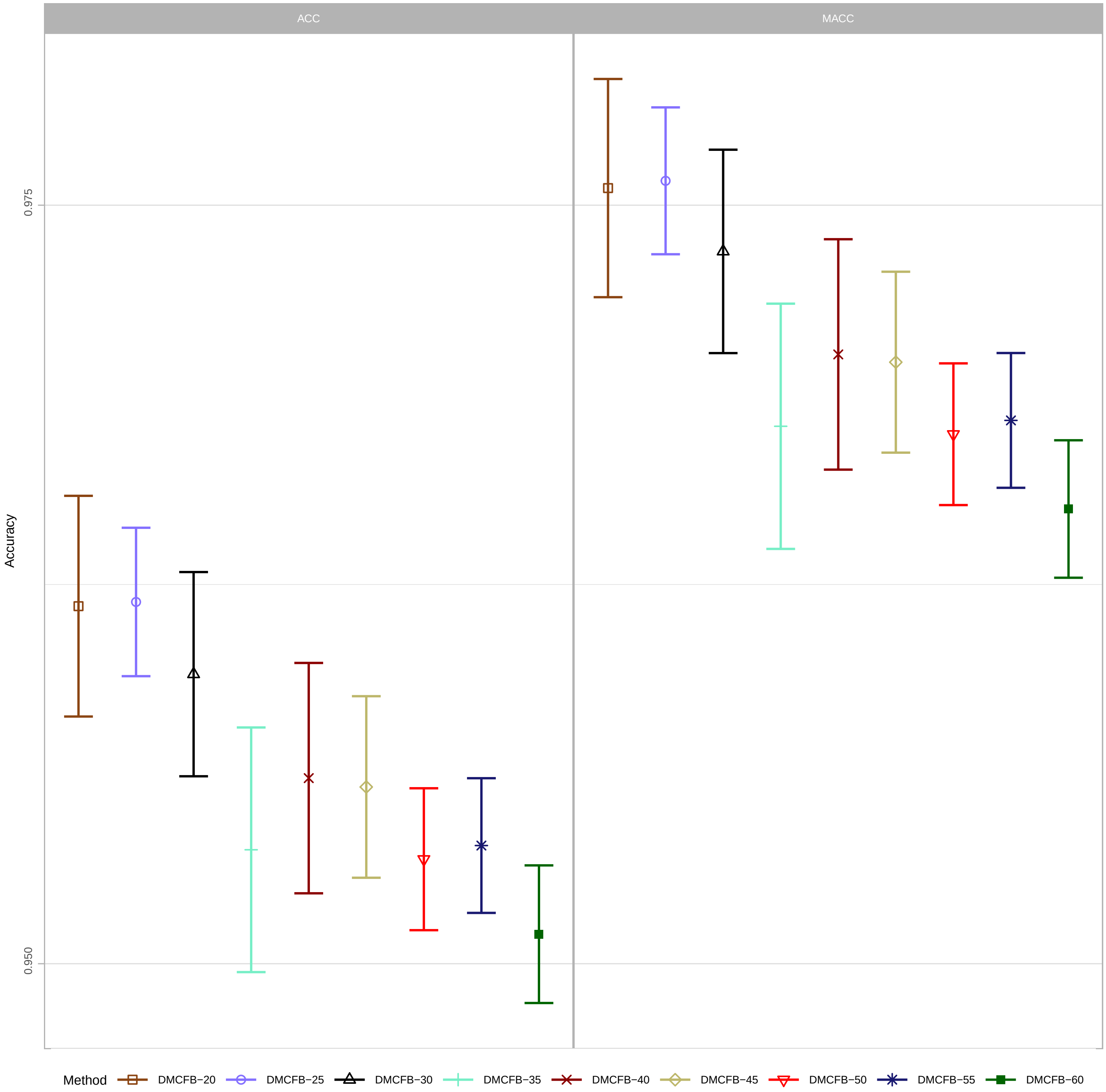


FIGURE S23: *The average overall accuracy (ACC) and average overall modified accuracy (MACC) for each choice of band-width in* `DMCFB` *in simulated data for the second scenario; Errors are generated from* $N(\mu = 0, \sigma = 0.18)$*. The vertical axis is truncated.*

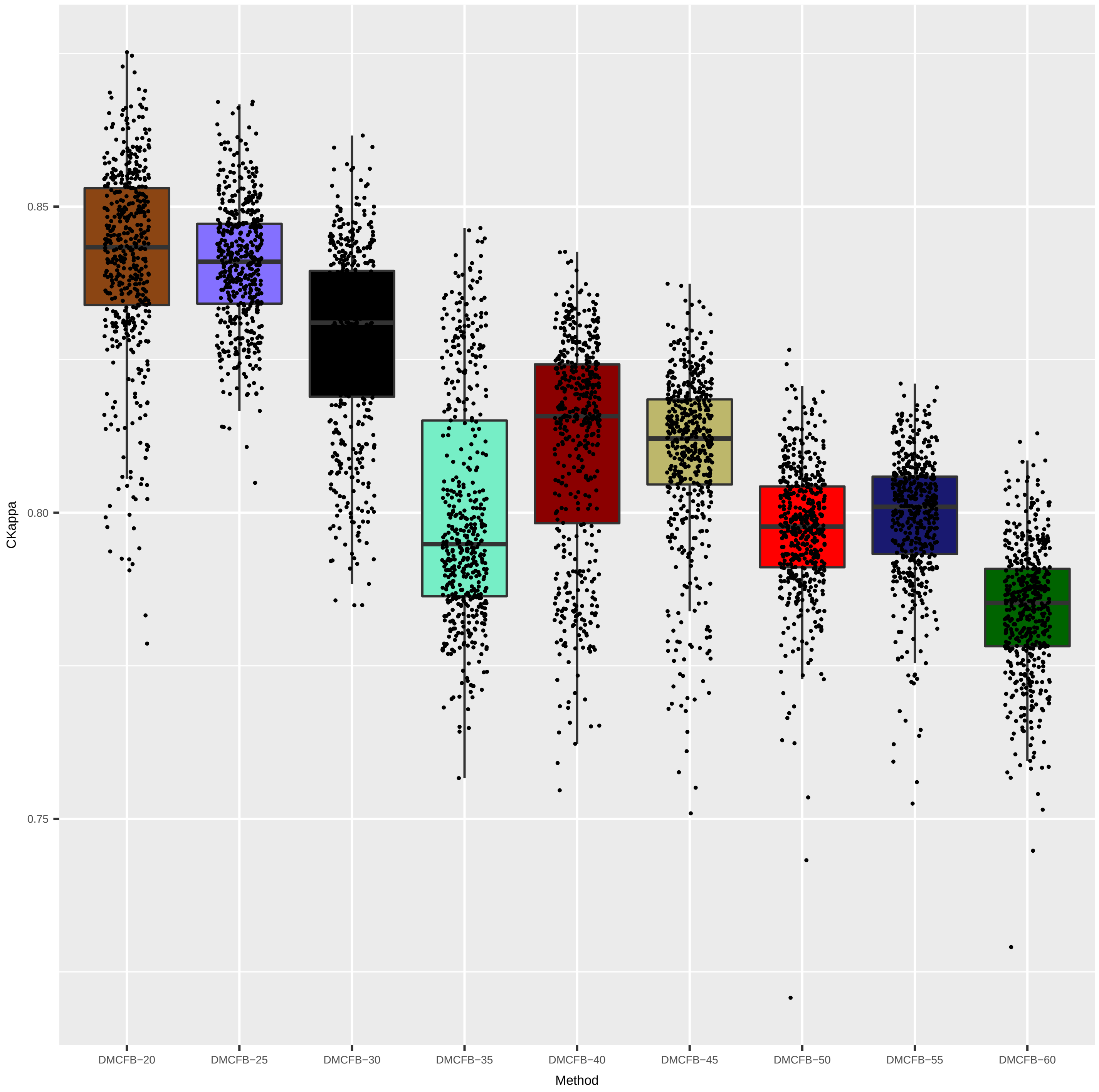


FIGURE S24: *The box-plots of Cohen's Kappa for each choice of band-width in* `DMCFB` *in simulated data for the second scenario; Errors are generated from* $N(\mu = 0, \sigma = 0.18)$*. The vertical axis is truncated.*

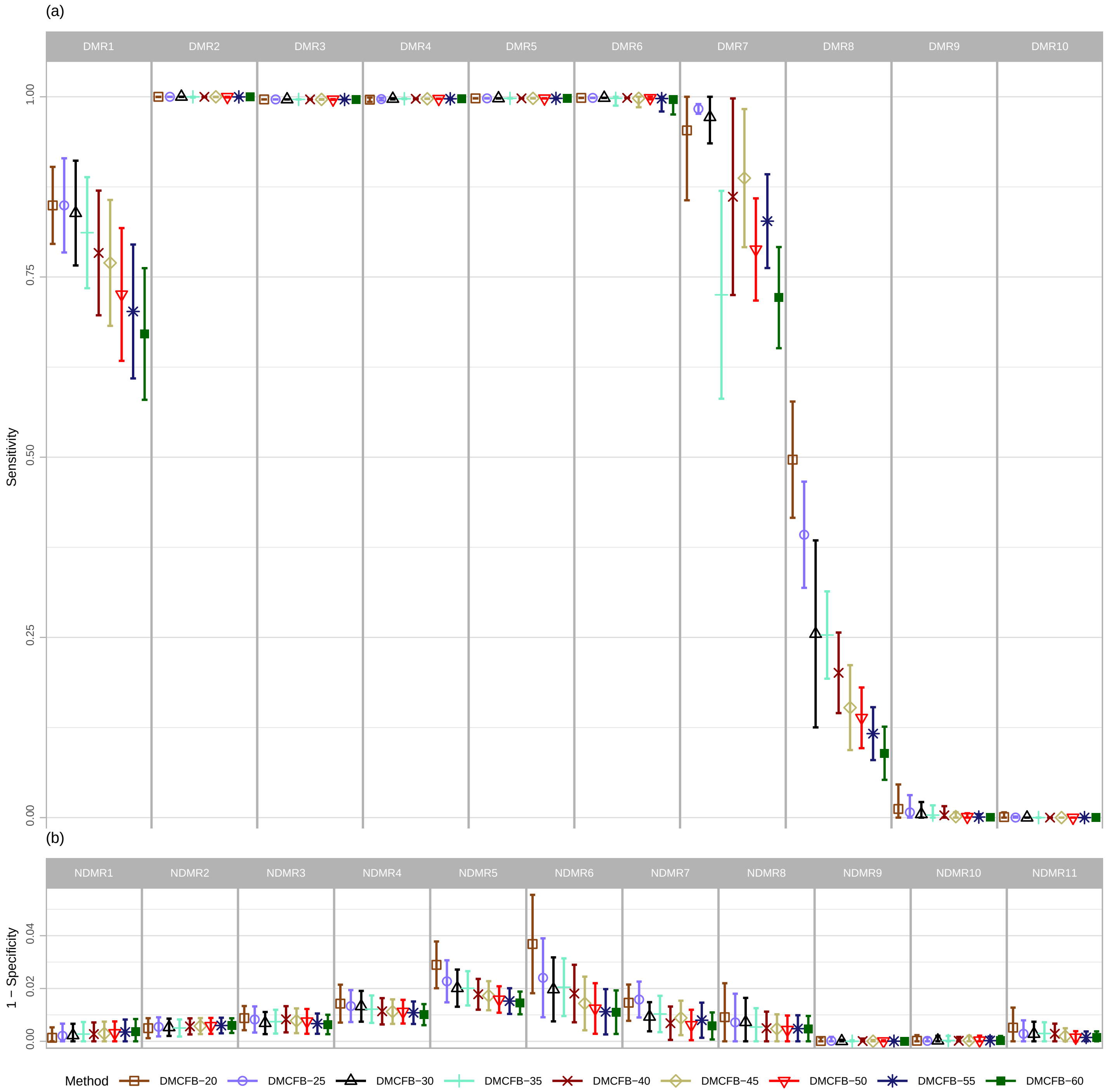


FIGURE S25: *(a) The average proportion of correctly identified* DMCs *(Sensitivity) for each choice of band-width separated by* DMRs *(sd error bars are added); (b) The average proportion of incorrectly identified* DMCs *(1 - Specificity) for each choice of band-width separated by* NDMRs *(The axis is truncated) in* **DMCFB** *in simulated data for the second scenario; Errors are generated from* $N(\mu = 0, \sigma = 0.18)$.

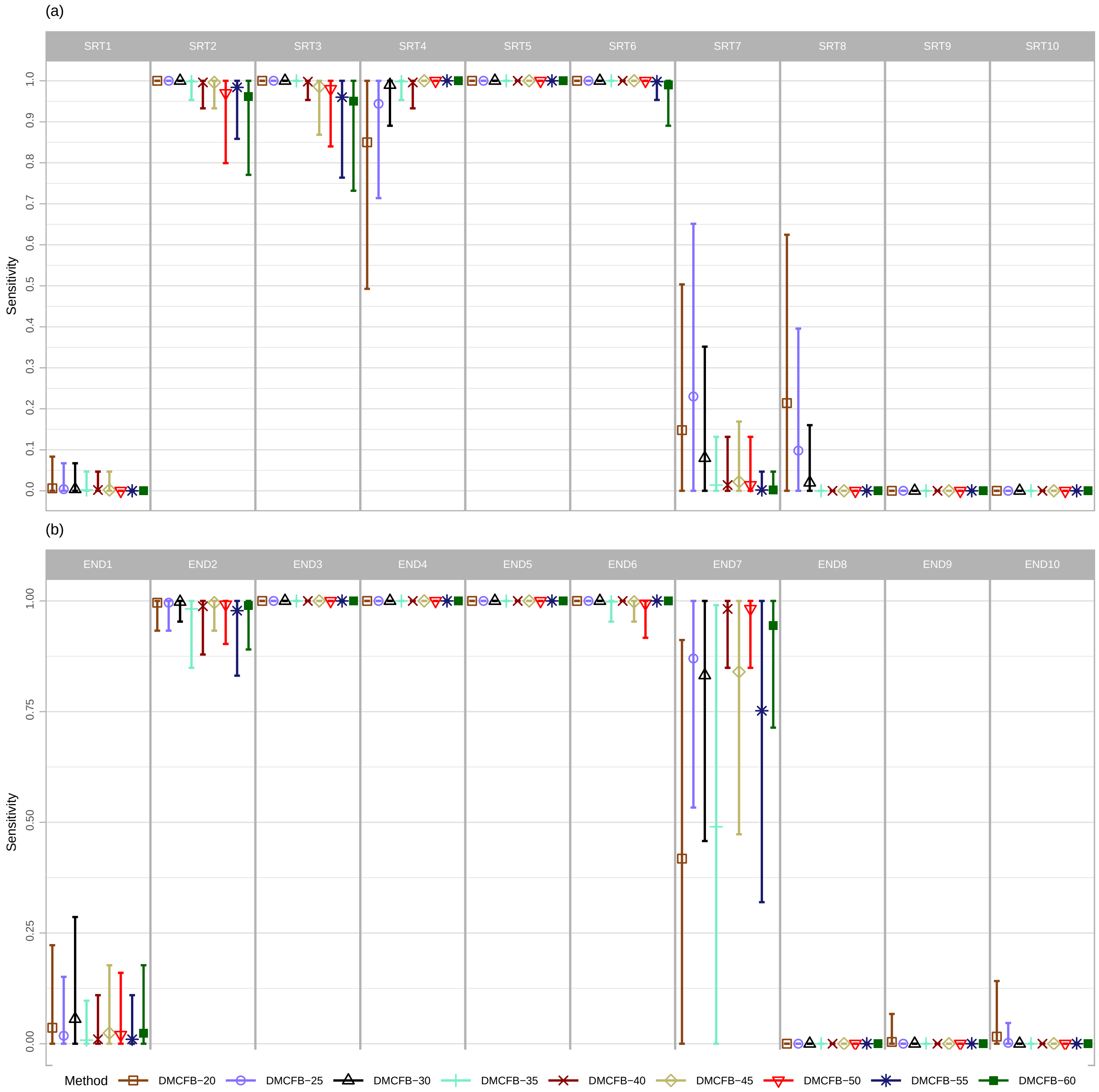


FIGURE S26: *Average number of times that the start (a) and end (b) positions of* DMRs *are identified as* DMCs *for each choice of band-width in* DMCFB *in simulated data for the second scenario; Errors are generated from* $N(\mu = 0, \sigma = 0.18)$.

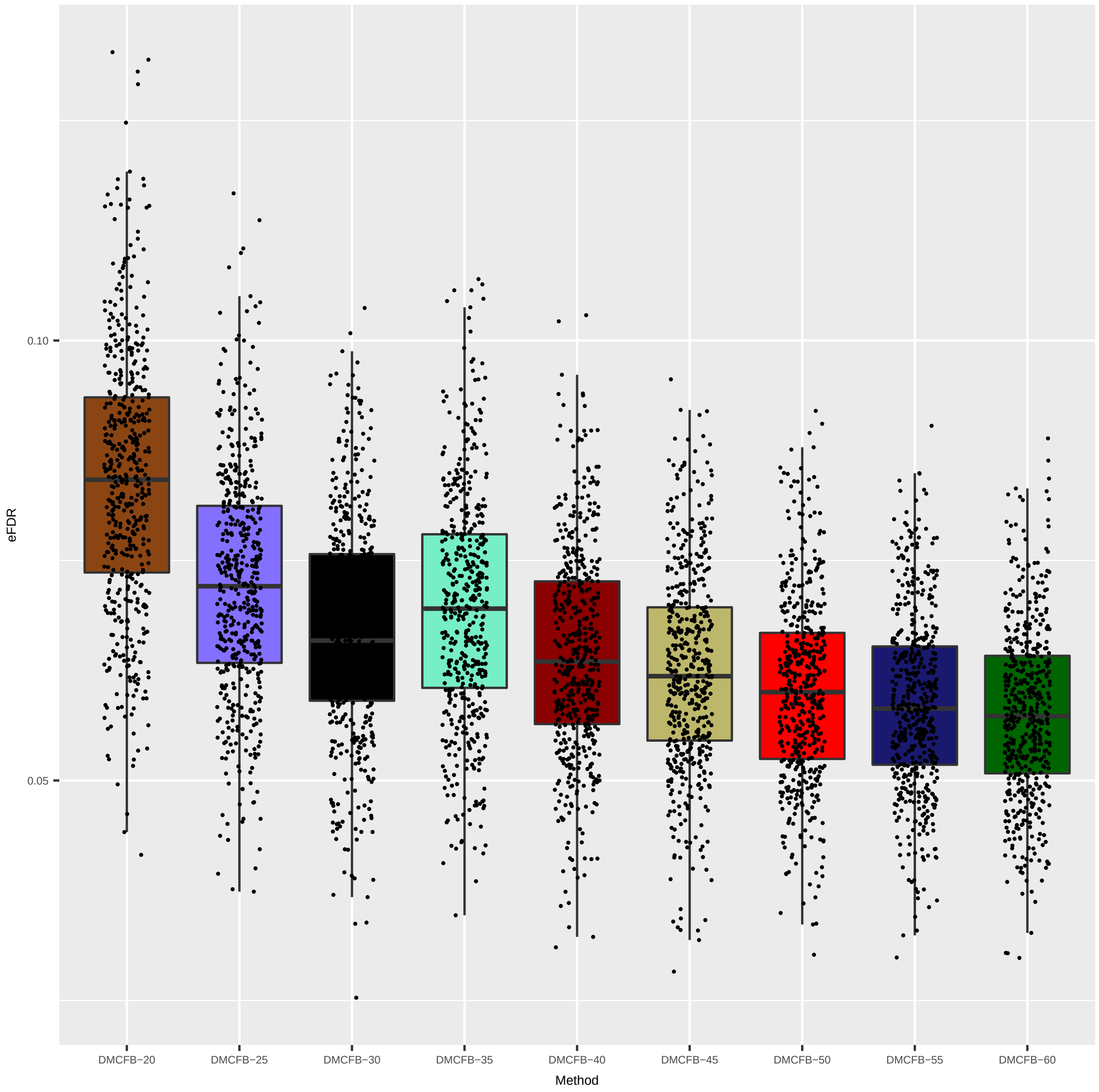


FIGURE S27: *Average empirical FDR for each choice of band-width in* `DMCFB` *in simulated data for the second scenario; Errors are generated from* $N(\mu = 0, \sigma = 0.18)$. *The vertical axis is truncated.*

#### S6.3. Sensitivity analysis with respect to prior precision; Scenario 1, $\sigma = 0.18$

A set of precision values in $\{30, 20, 10, 1/0.18, 1, 0.3, 0.2, 0.1\}$ is chosen for sensitivity analysis. In the following figures we provide the results.

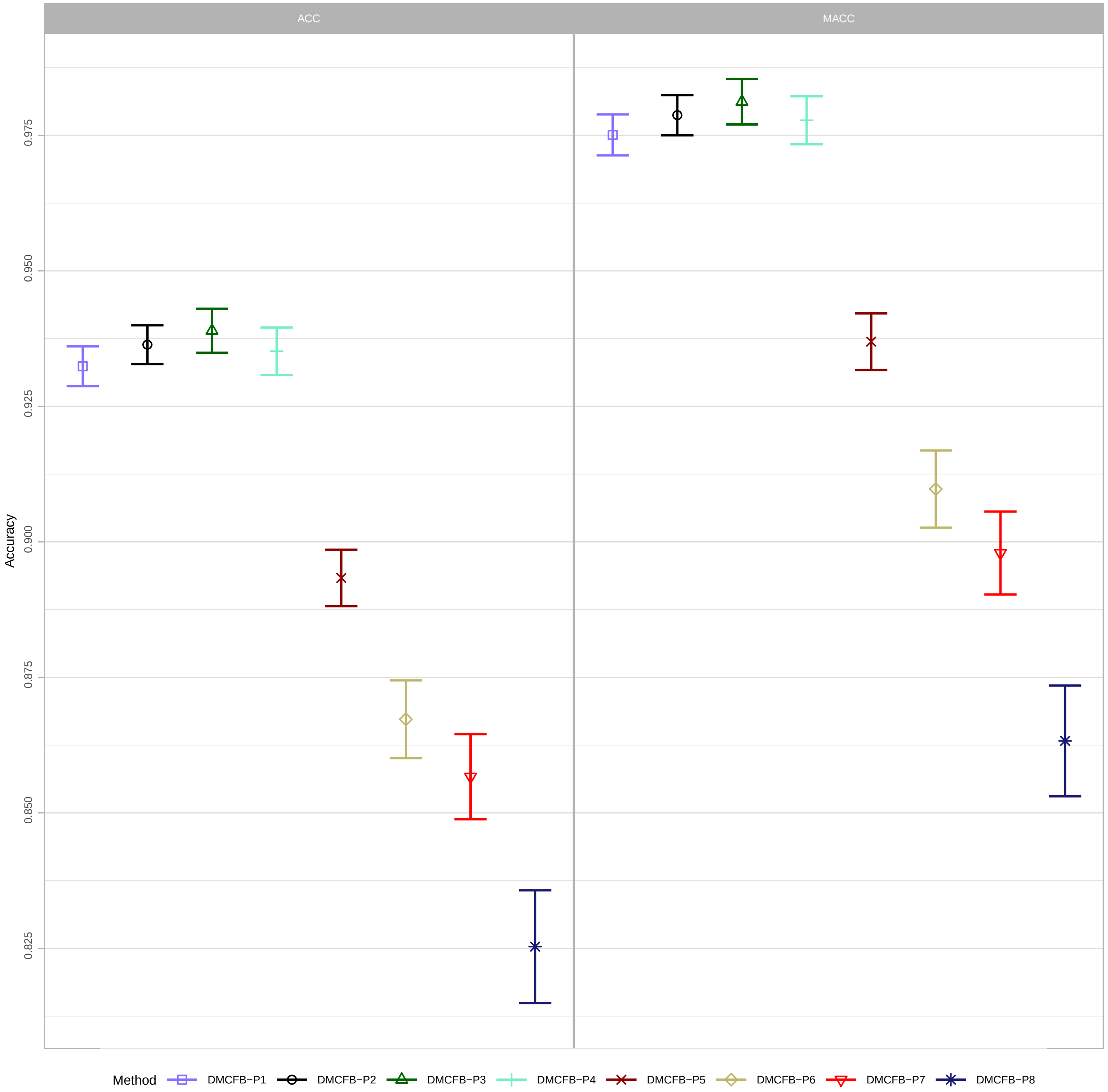


FIGURE S28: *The average overall accuracy (ACC) and average overall modified accuracy (MACC) for each choice of precision in `DMCFB` in simulated data for the first scenario; Errors are generated from $N(\mu = 0, \sigma = 0.18)$. The vertical axis is truncated.*

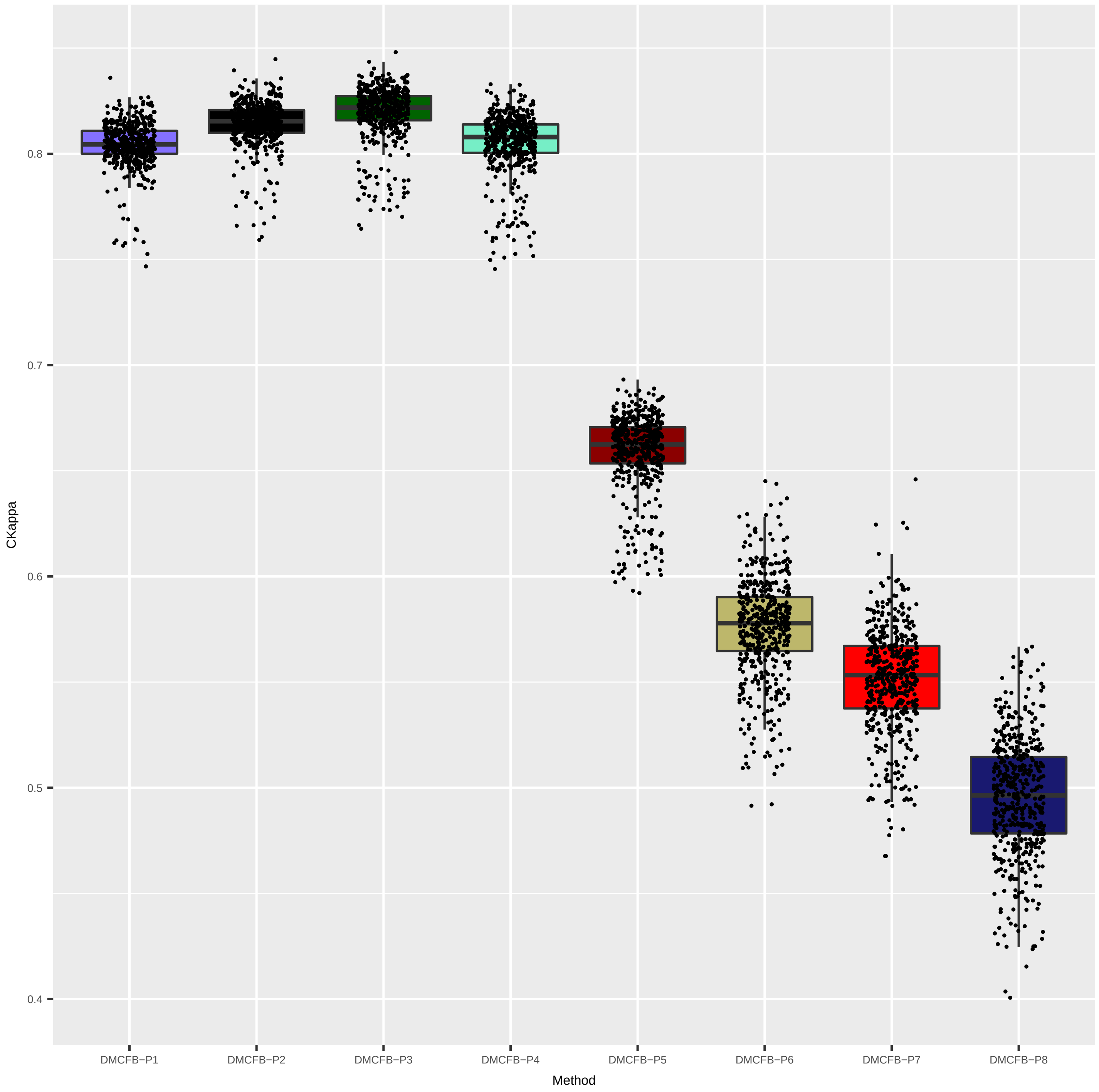


FIGURE S29: *The box-plots of Cohen's Kappa for each choice of precision in* `DMCFB` *in simulated data for the first scenario; Errors are generated from* $N(\mu = 0, \sigma = 0.18)$. *The vertical axis is truncated.*

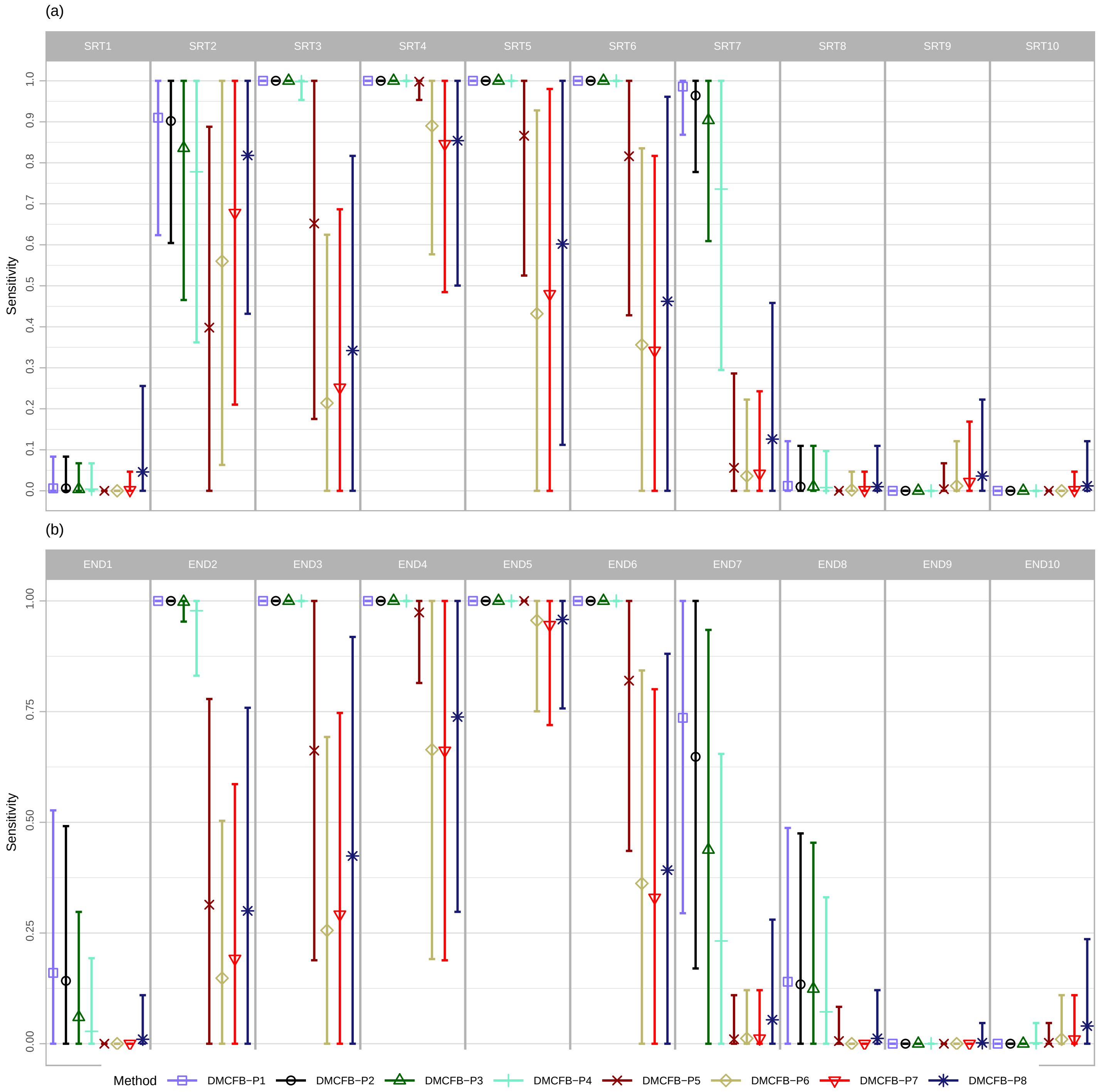


FIGURE S30: *Average number of times that the start (a) and end (b) positions of DMRs are identified as DMCs for each choice of precision in DMCFB in simulated data for the first scenario; Errors are generated from* $N(\mu = 0, \sigma = 0.18)$.

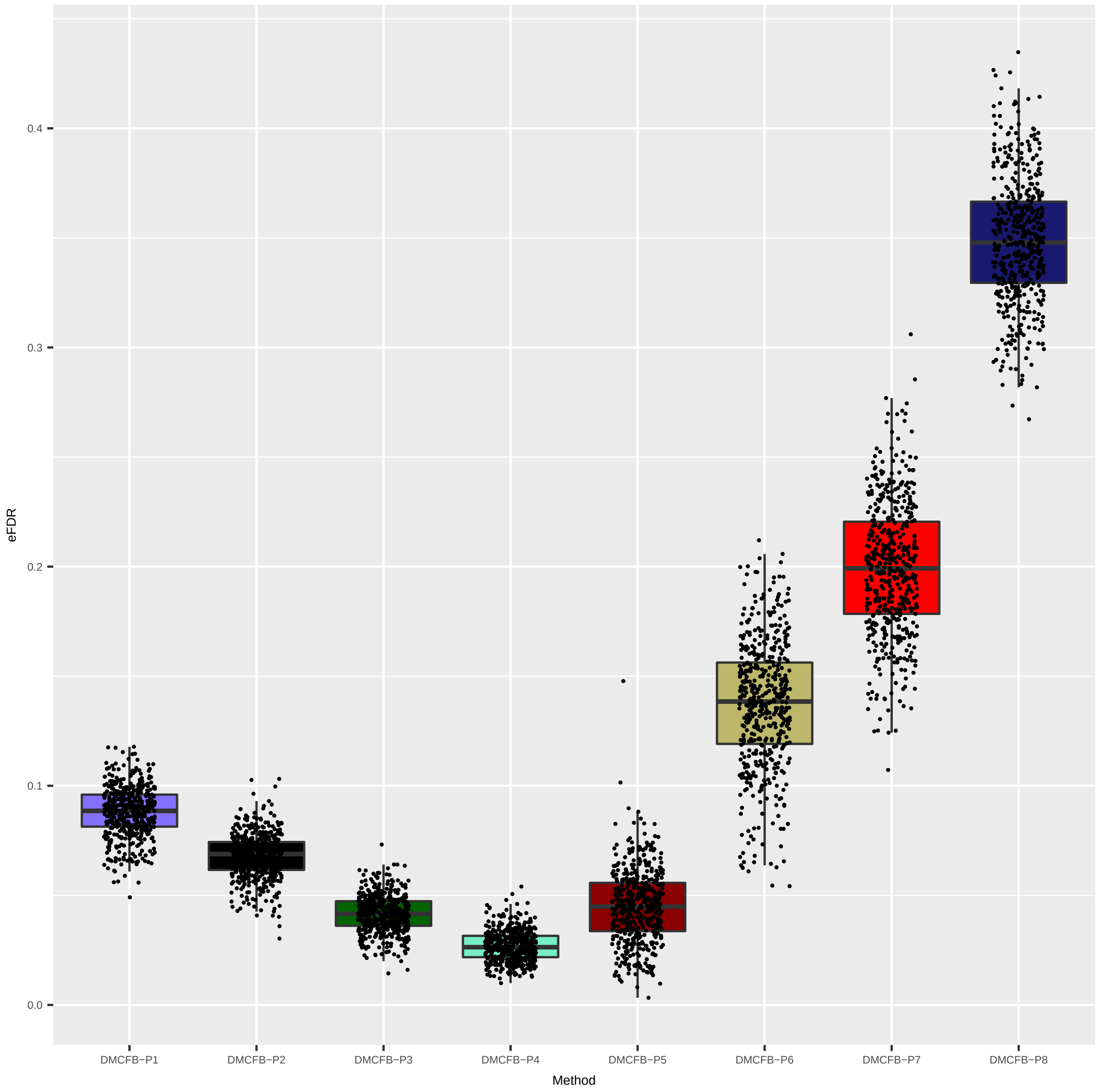


FIGURE S31: *Average empirical FDR for each choice of precision in* DMCFB *in simulated data for the first scenario; Errors are generated from* $N(\mu = 0, \sigma = 0.18)$. *The vertical axis is truncated.*

#### *S6.4. Sensitivity analysis with respect to pair band-width and partition size; Scenario 1,* $\sigma = 0.18$

Suppose a key region (with interesting associations with a grouping factor) is divided arbitrarily before the analysis. By re-assembling it afterwards, one may see or identify more clearly how a regulatory landscape is altered by the grouping factor. To study this we have chosen a set of pairs of band-width and partition size values as follows to assess its sensitivity in `DMCFB`. In the following table we provide the values.

TABLE S4
*Simulation set-up parameters for the first scenario*

| Set | Band-width size | Partition size |
|---|---|---|
| 1 | 30 | 500 |
| 2 | 45 | 750 |
| 3 | 60 | 1000 |
| 4 | 75 | 1250 |
| 5 | 90 | 1500 |
| 6 | 105 | 1750 |
| 7 | 120 | 2000 |
| 8 | 135 | 2250 |

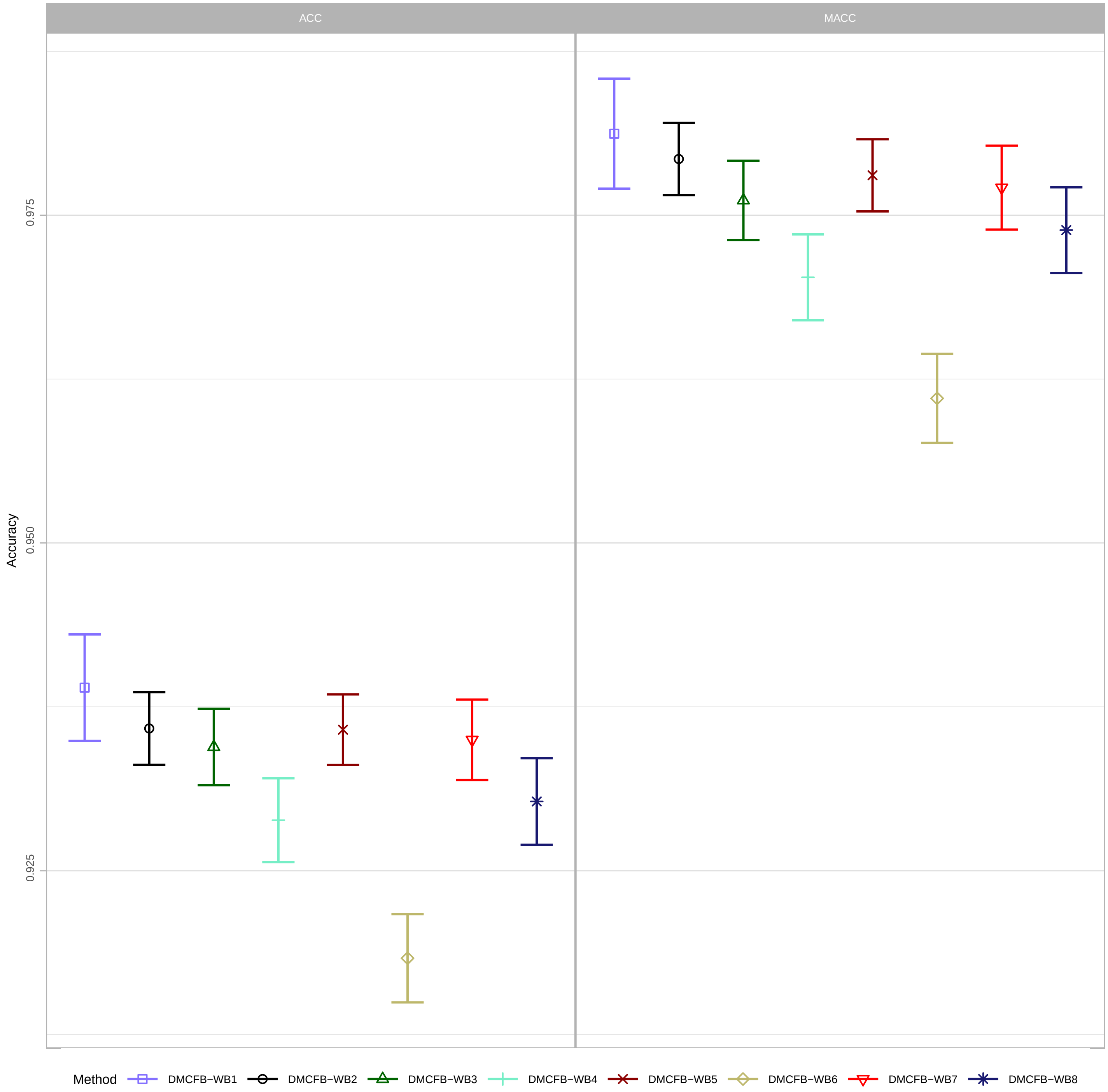


FIGURE S32: *The average overall accuracy (ACC) and average overall modified accuracy (MACC) for each choice of joint band-width and partition size in* `DMCFB` *in simulated data for the first scenario; Errors are generated from* $N(\mu = 0, \sigma = 0.18)$. *The vertical axis is truncated.*

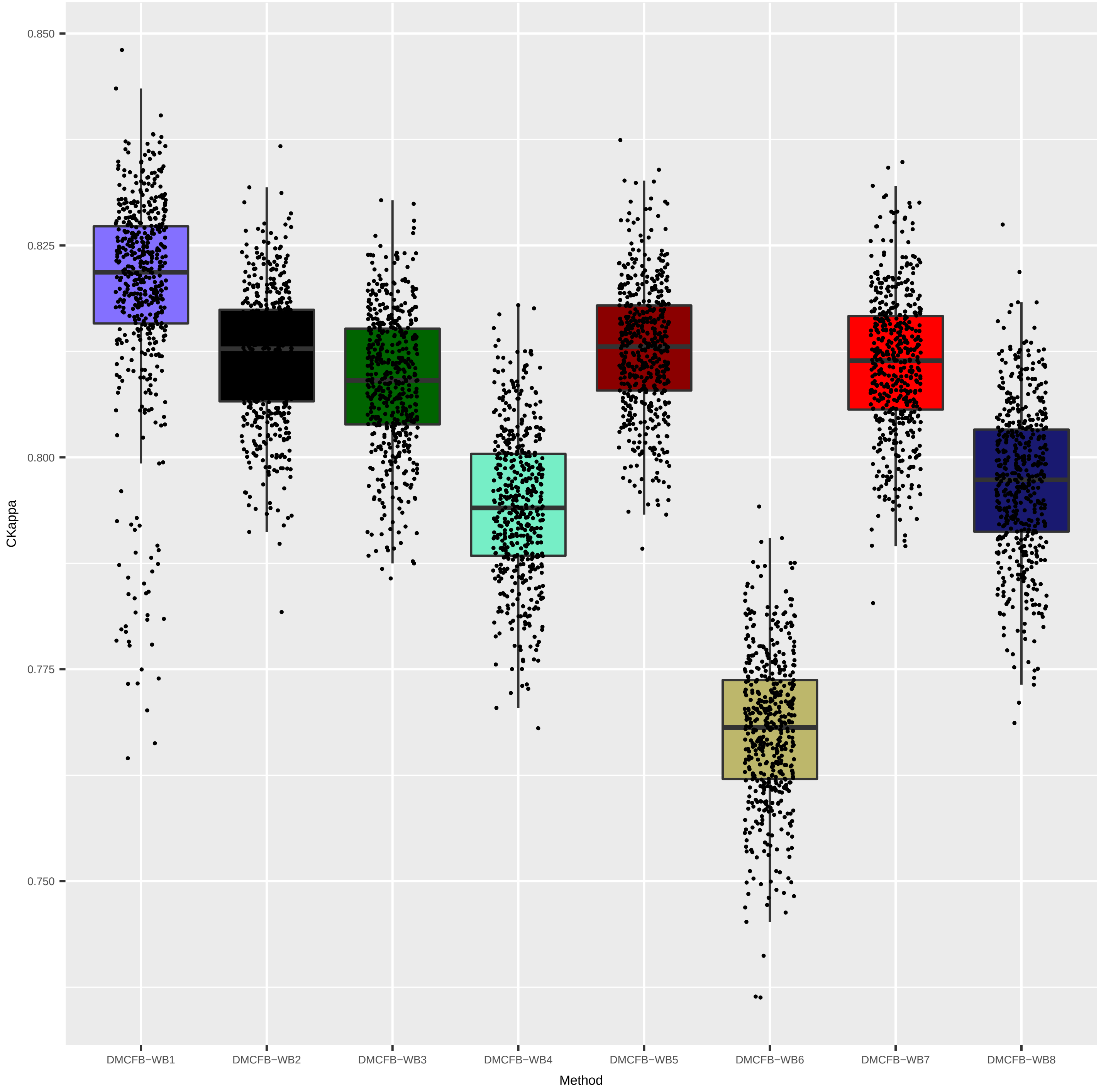


FIGURE S33: *Box-plots of Cohen's Kappa for each choice of joint band-width and partition size in* `DMCFB` *in simulated data for the first scenario; Errors are generated from* $N(\mu = 0, \sigma = 0.18)$. *The vertical axis is truncated.*

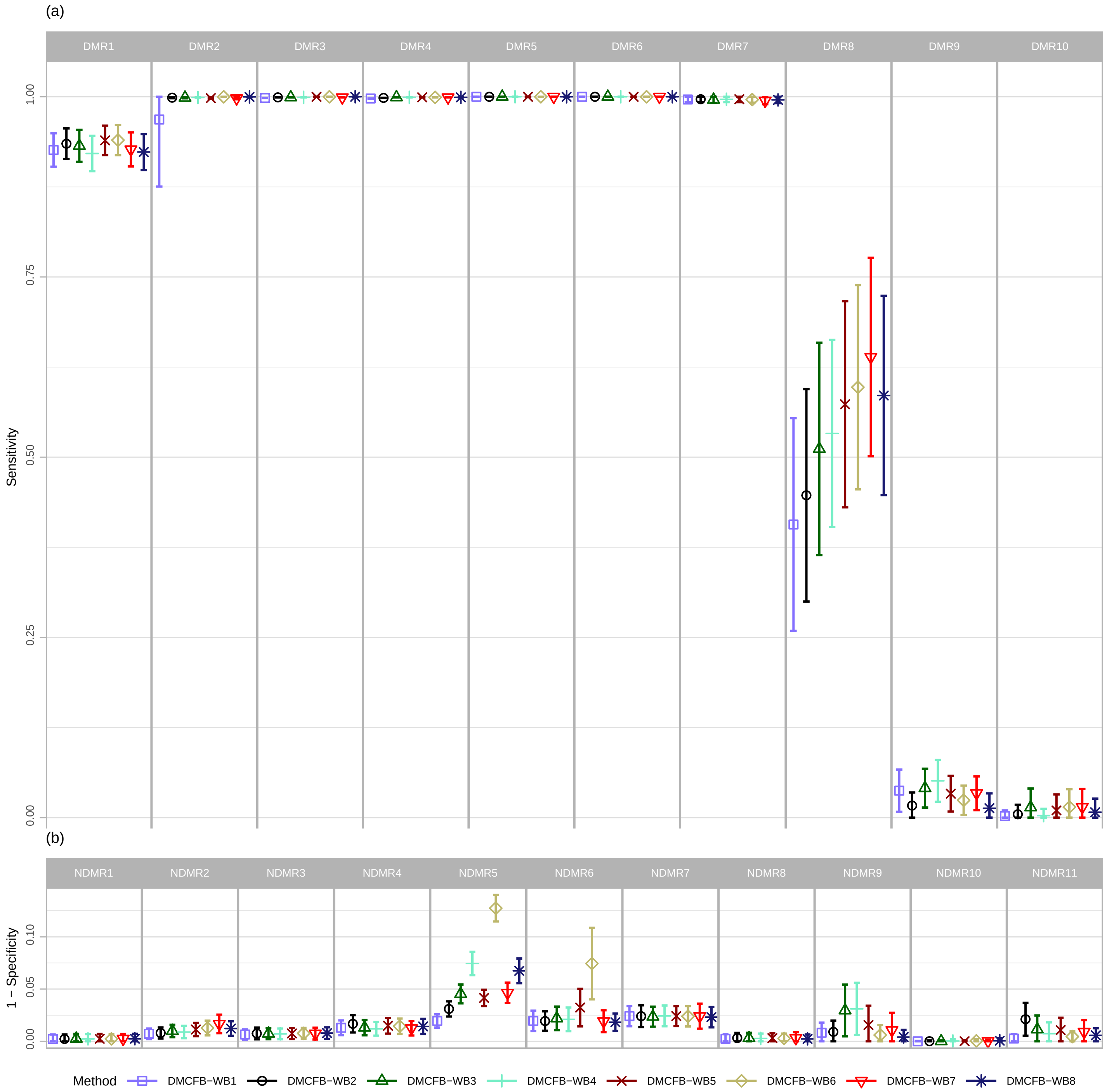


FIGURE S34: *(a) The average proportion of correctly identified* DMCs *(Sensitivity) for each choice of joint band-width and partition size separated by* DMRs *(sd error bars are added); (b) The average proportion of incorrectly identified as* DMCs *(1 - Specificity) for each choice of joint band-width and partition size separated by* NDMRs *(The axis is truncated) in* `DMCFB` *in simulated data for the first scenario; Errors are generated from* $N(\mu = 0, \sigma = 0.18)$.

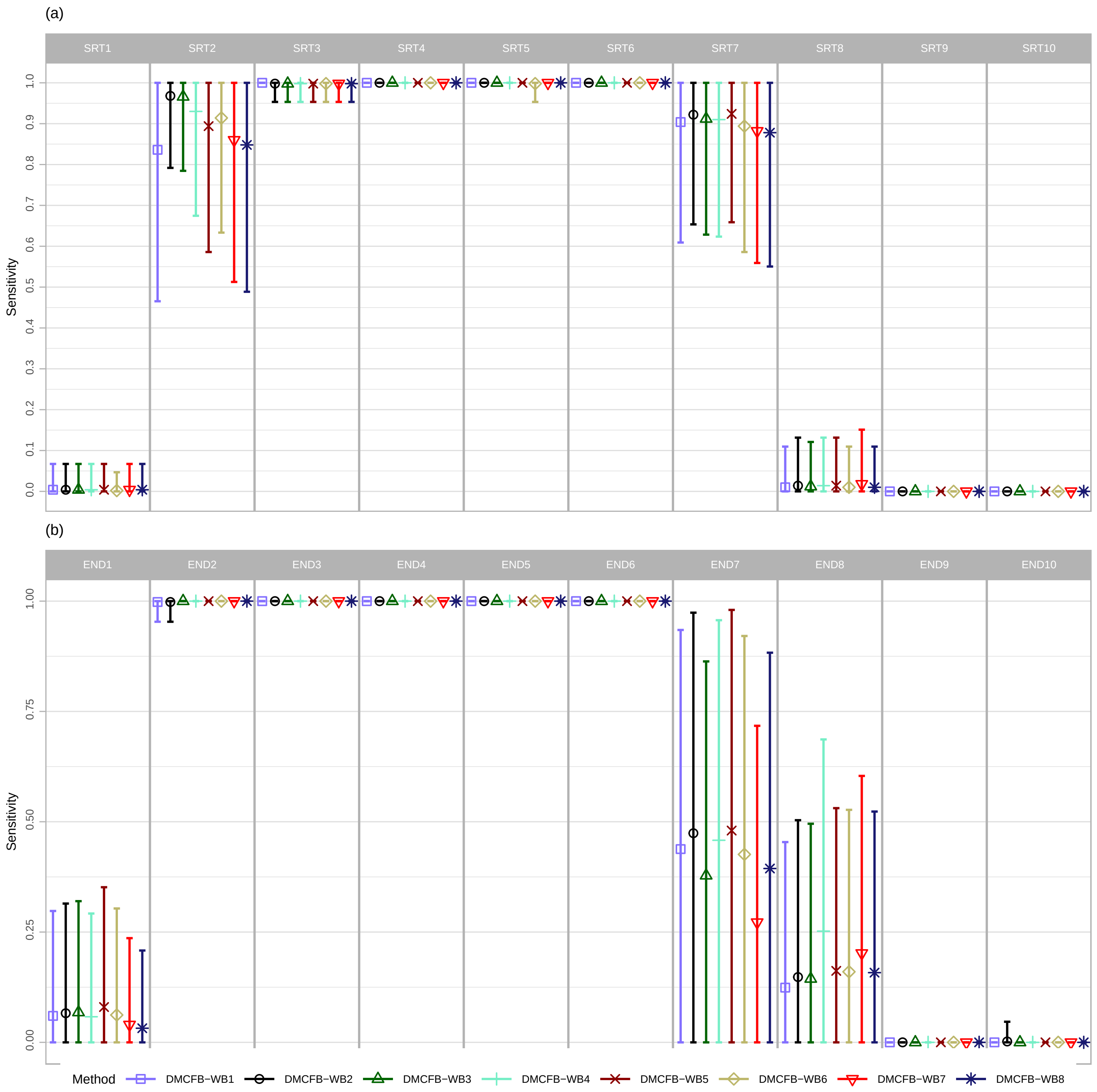


FIGURE S35: *Average number of times that the start (a) and end (b) positions of* DMRs *are identified as* DMCs *for each choice of joint band-width and partition size in* **DMCFB** *in simulated data for the first scenario; Errors are generated from* $N(\mu = 0, \sigma = 0.18)$.

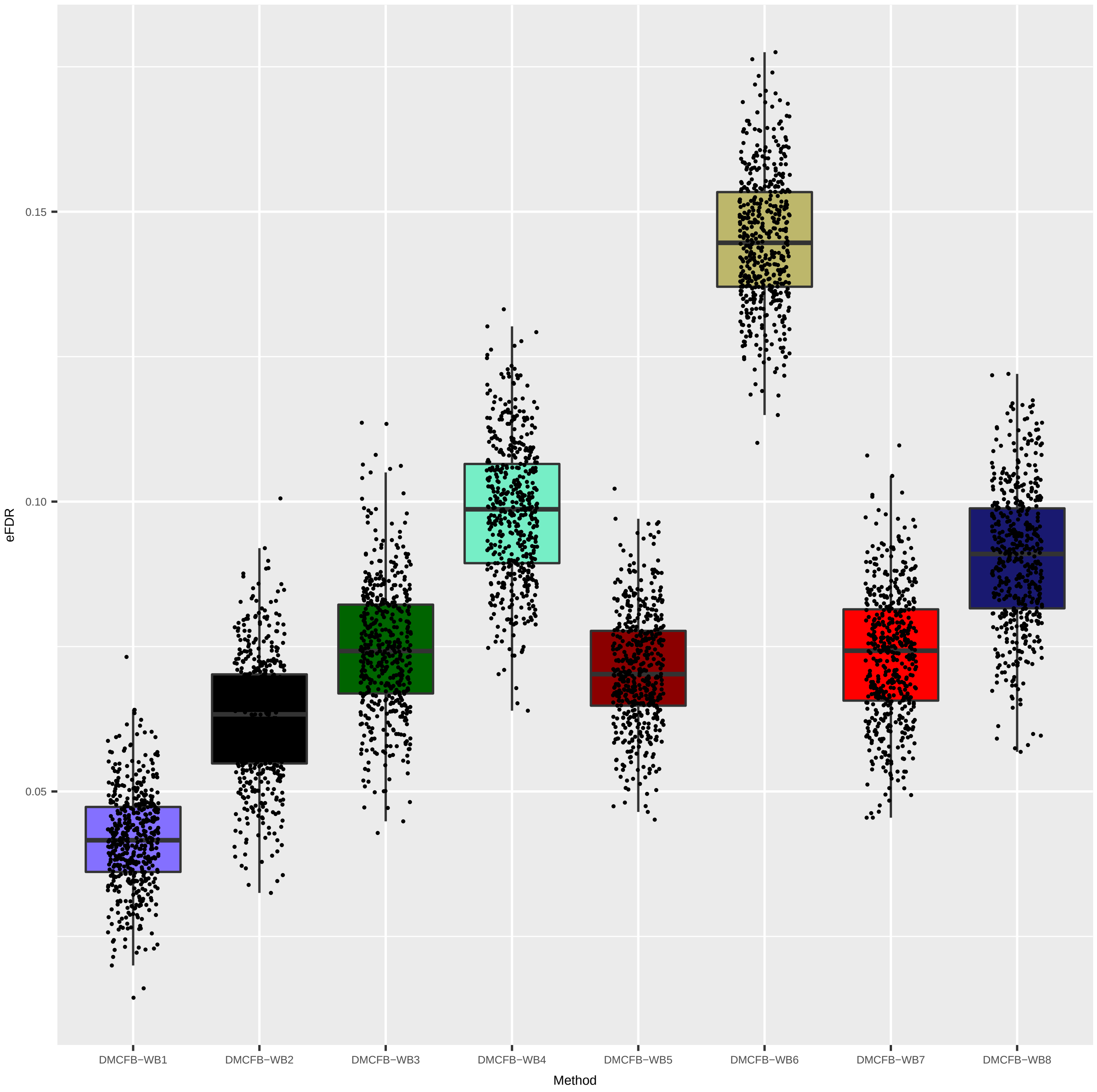


FIGURE S36: *Average empirical FDR for each choice of joint band-width and partition size in* DMCFB *in simulated data for the first scenario; Errors are generated from* $N(\mu = 0, \sigma = 0.18)$. *The vertical axis is truncated.*

#### S7. Analysis of the BLK data

We have compared methylation profiles between cell types near the BLK gene described in Section 5 of the main paper using DMCFB. Figure S37 shows pairwise DMCs for cell-type comparisons for the whole BLK region. We presented a graph of a small segment near the BLK gene in the main paper. We can observe that Monocyte cells and T-cells are hypermethylated compared to B-cells. Also, T-cells are hypomethylated compared to Monocyte cells. This fact stands out for the promoter of the BLK gene.

We also analyzed the BLK data using different FDR levels (Table S5). The FDR level $5 \times 10^{-2}$, which is quite liberal, identifies 63% of CpGs as DMCs, while the two FDR levels $1 \times 10^{-5}$ and $1 \times 10^{-8}$ show quite identical results; i.e., 36% of positions are DMCs.

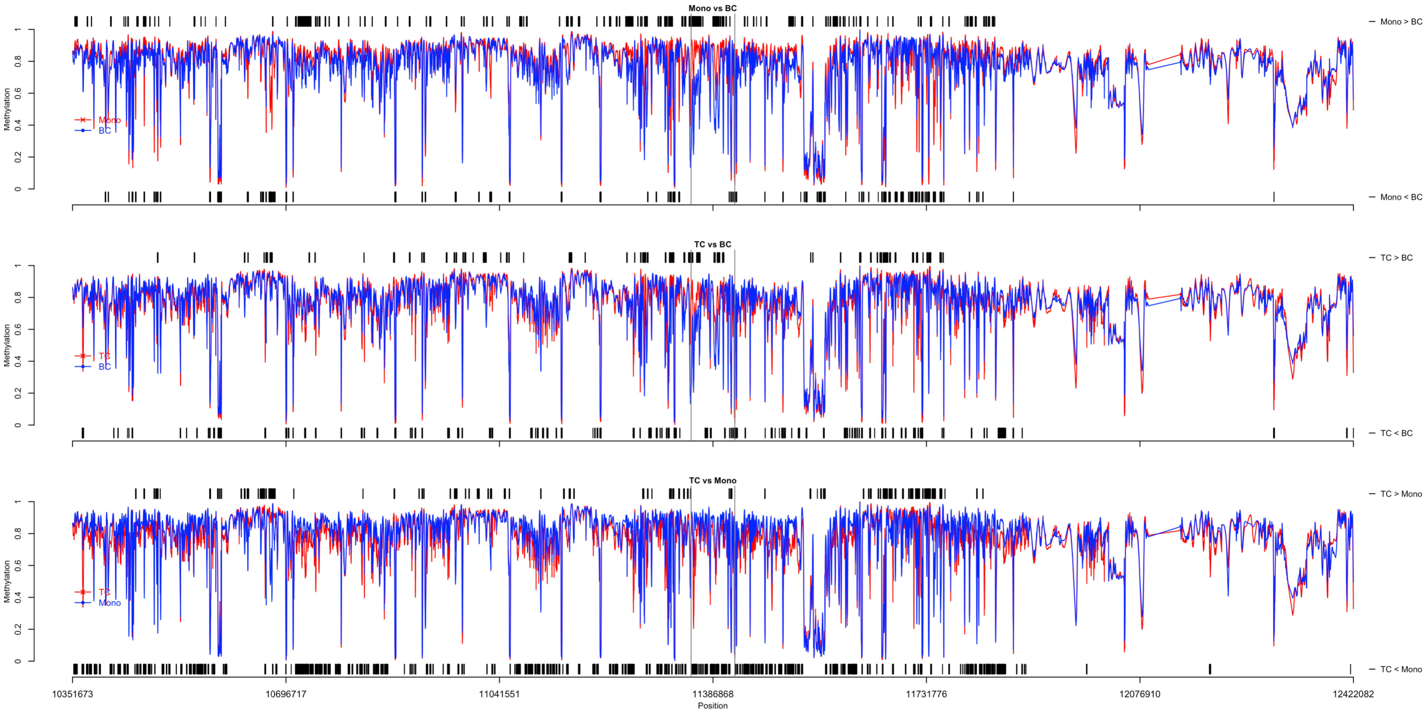


FIGURE S37: *(a)-(c) Identified DMCs for three pairwise comparisons of cell-type methylation data near the BLK gene. Short black vertical lines indicate CpGs where one cell type was significantly different from the other, at credible interval level $\alpha = 1 \times 10^{-8}$. The average of methylation level for each cell-type are drawn using black and grey lines. B-cells are less methylated than other cell-types in the promoter of BLK.*

Table S5
BLK *data analysis on Chromosome 8 using* DMCFB.

| $\alpha$ | Pairwise contrasts | | NDMC[1] % | DMC % | Hypermethylated % | Hypomethylated % |
|---|---|---|---|---|---|---|
| $5 \times 10^{-2}$ | Mono | BC | 59.79 | 40.21 | 23.66 | 16.55 |
| | TC | BC | 67.73 | 32.18 | 10.43 | 21.85 |
| | TC | Mono | 55.61 | 44.38 | 11.10 | 33.28 |
| | All positions | | 37.16 | 62.84 | – | – |
| $1 \times 10^{-5}$ | Mono | BC | 81.53 | 18.46 | 8.62 | 9.84 |
| | TC | BC | 85.62 | 14.38 | 4.93 | 9.45 |
| | TC | Mono | 75.82 | 27.17 | 6.35 | 17.82 |
| | All positions | | 63.84 | 36.16 | – | – |
| $1 \times 10^{-8}$ | Mono | BC | 81.54 | 18.46 | 8.62 | 9.84 |
| | TC | BC | 85.63 | 14.37 | 4.92 | 9.44 |
| | TC | Mono | 75.83 | 24.17 | 6.35 | 17.82 |
| | All positions | | 63.86 | 36.14 | – | – |

[1]Not a DMC

#### S8. Analysis of the APL data

##### S8.1. Additional results from the reanalysis of APL data in the main paper

We reanalyzed the data set of patients with Acute Promyelocytic Leukemia (APL), published in Schoofs et al. (2013), publicly available on the Gene Expression Omnibus (accession no. GSE42119) in the main paper. These RRBS data contain 18 APL patients and 16 Control samples (which include 8 bone marrow samples from patients in remission, four healthy $CD34^+$ cells and four promyelocytes). APL is characterized by a chromosomal translocation involving the retinoic acid receptor-alpha gene on Chromosome 15 and 17 (Coombs, Tavakkoli and Tallman, 2015).

In what follows, we present some of the findings discussed in the main paper.

Table S6
*The percentage of CpGs identified as DMC by DMCFB and BiSeq displayed; Chromosomes 15 and 17.*

| Chromosome | DMCFB | BiSeq |
|---|---|---|
| 15 | 36.31 | 6.48 |
| 17 | 37.14 | 3.67 |

Table S7
*The number of CpGs identified as DMC by only DMCFB, only BiSeq and both methods at the same time, separated for CpG Islands and Shores; Chromosomes 15 and 17.*

| Chromosome | Location | DMCFB | BiSeq | Both |
|---|---|---|---|---|
| 15 | Islands | 594 | 218 | 207 |
| | Shores | 823 | 196 | 181 |
| 17 | Islands | 1171 | 279 | 265 |
| | Shores | 1931 | 160 | 146 |

Table S8

*Known binding sites identified as methylated by* `DMCFB` *and* `BiSeq`*; Chromosomes 15 and 17.*

| Chromosome | Start Position | End Position | Region ID | Gene Symbol | # of DMCs found by `DMCFB` | # of DMCs found by |
|---|---|---|---|---|---|---|
| 15 | 88578829 | 88579456 | PR253 | CIB1 LOC390637 | 3 | 0 |
| | 78056685 | 78059161 | PR275 | BCL2A1 | 3 | 0 |
| | 70448192 | 70448955 | PR847 | C15orf34 | 4 | 0 |
| | 75500584 | 75501136 | PR1239 | HMG20A | 2 | 0 |
| | 75073824 | 75074470 | PR1257 | PSTPIP1 | 20 | 0 |
| | 78856391 | 78857171 | PR1411 | KIAA1199 | 7 | 0 |
| | 43279416 | 43279955 | PR1574 | SHF | 7 | 0 |
| | 75499827 | 75500583 | PR1719 | HMG20A | 12 | 3 |
| | 70449306 | 70449834 | PR1886 | C15orf34 | 5 | 0 |
| | 90196321 | 90197028 | PR1920 | SLCO3A1 | 2 | 0 |
| | 90198507 | 90199221 | PR2167 | SLCO3A1 | 60 | 8 |
| | 73034713 | 73035433 | PR2463 | RPP25 | 1 | 0 |
| | 41207073 | 41207569 | PR2477 | TMEM62 | 1 | 0 |
| | 39571039 | 39571637 | PR2851 | ITPKA | 12 | 0 |
| 17 | 35508454 | 35509090 | PR202 | NR1D1 | 9 | 0 |
| | 25901212 | 25902428 | PR326 | DKFZP434O047 | 2 | 0 |
| | 73390073 | 73390889 | PR493 | FLJ45079 | 13 | 0 |
| | 78389194 | 78390110 | PR501 | ZNF750 | 10 | 0 |
| | 29720010 | 29720505 | PR717 | CCL1 | 7 | 0 |
| | 70106163 | 70107547 | PR718 | | 7 | 0 |
| | 200408 | 201496 | PR723 | RPH3AL | 1 | 0 |
| | 29715488 | 29716536 | PR873 | CCL1 | 1 | 0 |
| | 29717010 | 29717915 | PR980 | CCL1 | 30 | 5 |
| | 45859216 | 45859972 | PR1046 | FLJ20920 | 2 | 0 |
| | 7403202 | 7403881 | PR1086 | SENP3 TNFSF13 | 3 | 0 |
| | 37862039 | 37863088 | PR1095 | ATP6V0A1 | 9 | 0 |
| | 36717510 | 36718797 | PR1125 | | 177 | 125 |
| | 21131431 | 21132190 | PR1196 | MAP2K3 | 16 | 0 |
| | 76883854 | 76884753 | PR1437 | MGC15523 | 4 | 0 |
| | 3570012 | 3570716 | PR1467 | GSG2 | 6 | 0 |
| | 15790787 | 15791354 | PR1551 | ADORA2B | 5 | 0 |
| | 29720506 | 29721319 | PR1566 | CCL1 | 1 | 0 |
| | 2248484 | 2249021 | PR1598 | | 5 | 0 |
| | 1533274 | 1533967 | PR1606 | PRPF8 | 3 | 0 |
| | 37589221 | 37589717 | PR1611 | KCNH4 HCRT | 16 | 0 |
| | 34262533 | 34263888 | PR1616 | RPL23 | 9 | 0 |
| | 37792477 | 37793256 | PR1638 | STAT3 | 12 | 0 |
| | 70055200 | 70055848 | PR1639 | CD300C | 5 | 0 |
| | 8031673 | 8032581 | PR1663 | C17orf59 | 16 | 0 |
| | 17679278 | 17680086 | PR1746 | SREBF1 | 2 | 0 |
| | 8138753 | 8139459 | PR1821 | SLC25A35 | 8 | 0 |
| | 1312519 | 1313086 | PR1858 | CRK | 22 | 0 |
| | 70053711 | 70054472 | PR1995 | CD300C | 7 | 0 |
| | 8031211 | 8031672 | PR2034 | | 16 | 0 |
| | 7402038 | 7403201 | PR2253 | SENP3 TNFSF13 | 5 | 0 |
| | 27692272 | 27693059 | PR2412 | RHOT1 C17orf75 | 7 | 0 |
| | 21128960 | 21129724 | PR2466 | MAP2K3 | 7 | 0 |
| | 75384560 | 75385252 | PR2479 | CBX8 | 27 | 0 |
| | 7322452 | 7323068 | PR2508 | POLR2A | 4 | 0 |
| | 9878588 | 9879133 | PR2522 | GAS7 | 5 | 0 |
| | 3814601 | 3815391 | PR2611 | ATP2A3 | 10 | 0 |
| | 9876834 | 9877589 | PR2612 | | 7 | 0 |
| | 4789994 | 4790485 | PR2775 | ENO3 SLC25A11 | 6 | 0 |
| | 74441013 | 74441653 | PR2777 | TIMP2 | 7 | 0 |
| | 8017956 | 8018495 | PR2853 | TMEM107 | 2 | 0 |
| | 43125653 | 43126114 | PR2944 | TBKBP1 | 7 | 0 |
| | 39781024 | 39781679 | PR2958 | | 5 | 0 |

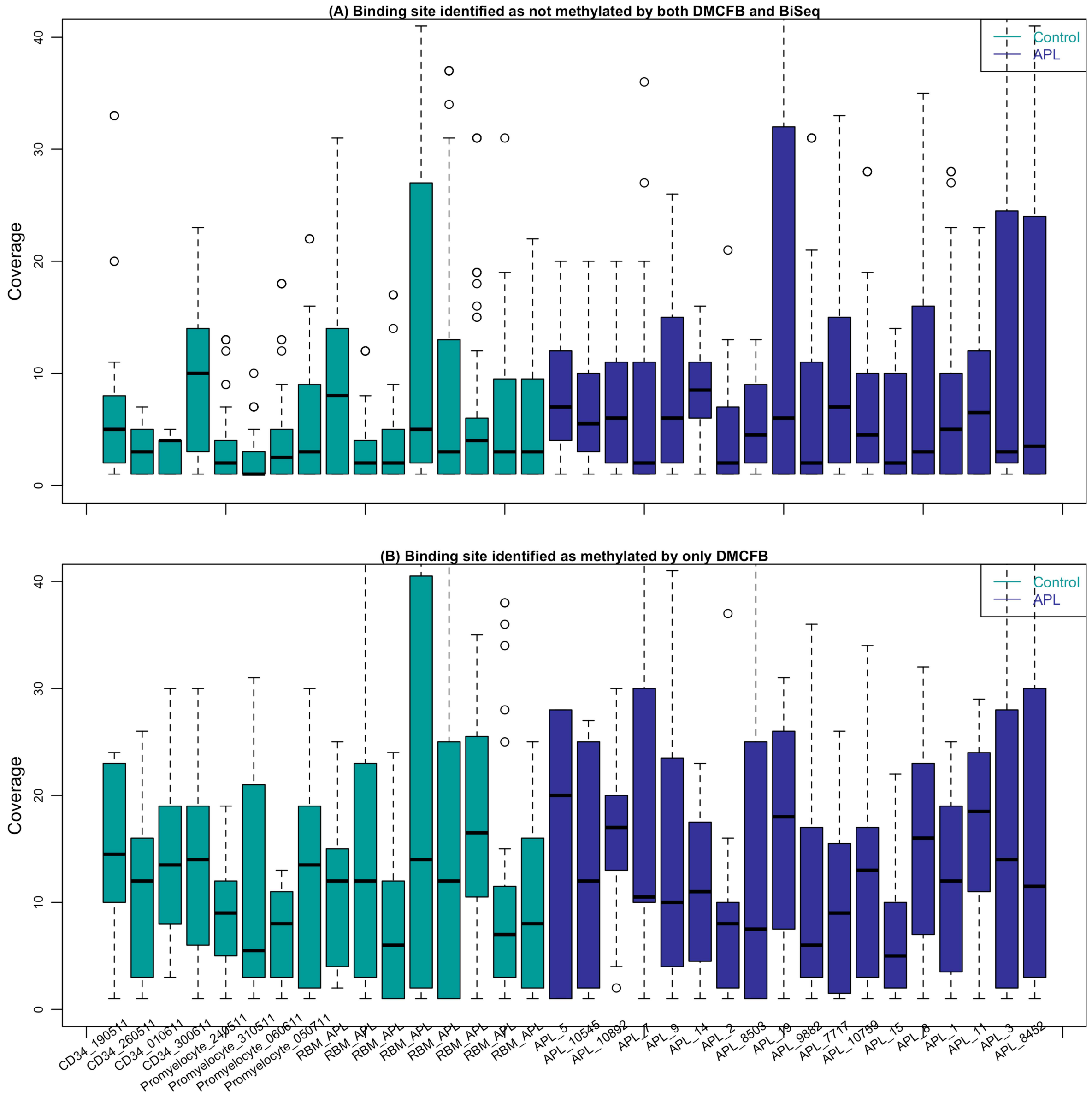


FIGURE S38: *(A) Box-plots of read-depth of CpG positions in known binding sites identified as not* DMCs *by* `DMCFB` *and* `BiSeq`*; (B) Box-plots of read-depth of CpG positions in known binding sites identified as* DMCs *by only* `DMCFB` *and not* `BiSeq`*. Chromosome 15.*

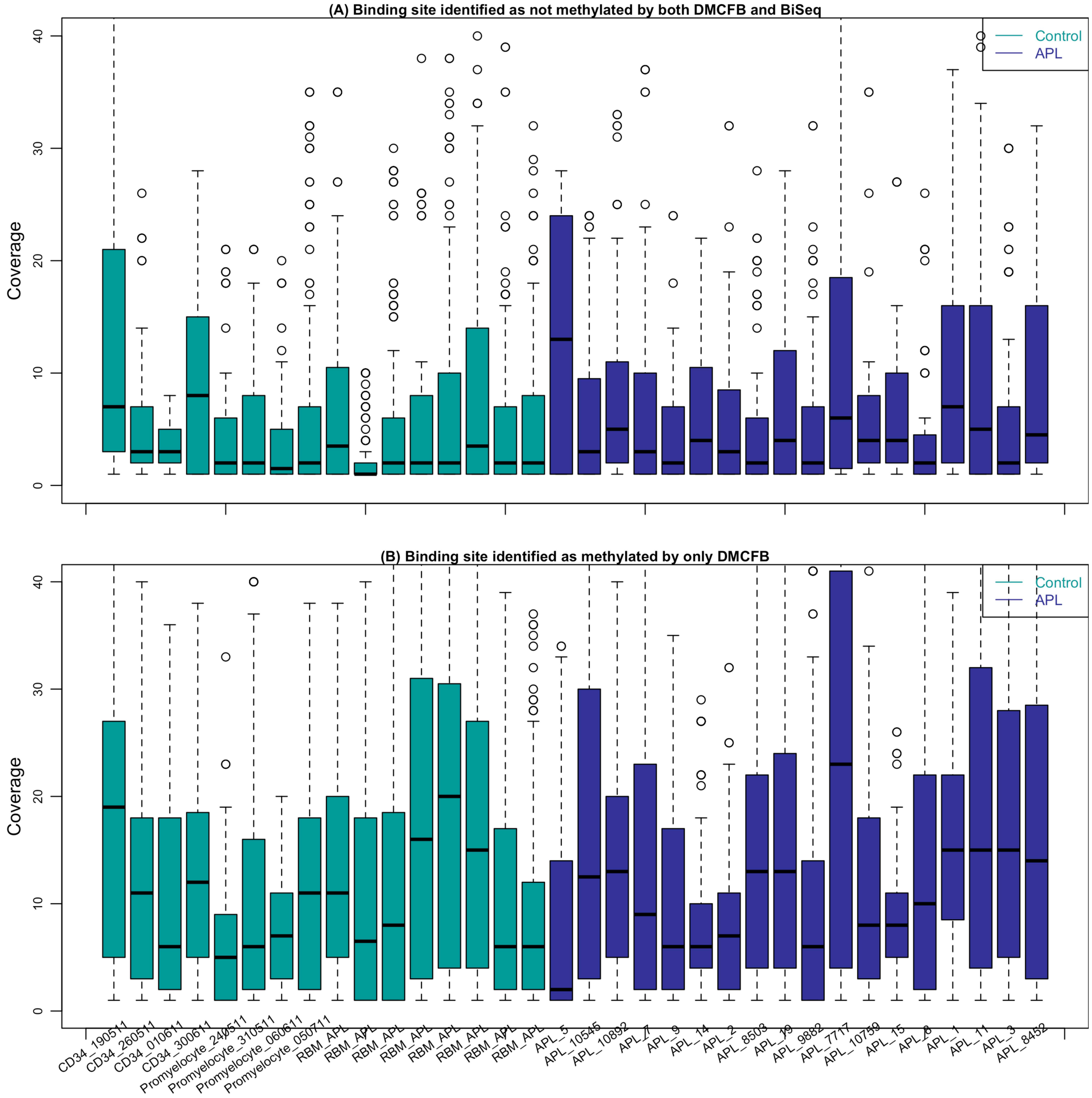


Figure S39: *(A) Box-plots of read-depth of CpG positions in known binding sites identified as not* DMCs *by* `DMCFB` *and* `BiSeq`*; (B) Box-plots of read-depth of CpG positions in known binding sites identified as* DMCs *by only* `DMCFB` *and not* `BiSeq`*. Chromosome 17.*

Table S9

*Counts of Shore regions are shown, categorized by whether the adjacent Island contains at least one* DMC *and whether the Shore (in the same direction) contains at least one* DMC. *Counts are shown* `DMCFB` *and* `BiSeq`*; Chromosomes 15 and 17.*

| | # of Shores | | | With DMC | |
|---|---|---|---|---|---|
| Chromosome | `DMCFB` | `BiSeq` | Stream | Island | Shore |
| 15 | 372 | 64 | Up | ✓ | ✓ |
| | 363 | 78 | Down | ✓ | ✓ |
| | 222 | 154 | Up | ✓ | ✗ |
| | 231 | 140 | Down | ✓ | ✗ |
| | 38 | 24 | Up | ✗ | ✓ |
| | 50 | 30 | Down | ✗ | ✓ |
| | 160 | 550 | Up | ✗ | ✗ |
| | 148 | 544 | Down | ✗ | ✗ |
| 17 | 770 | 40 | Up | ✓ | ✓ |
| | 789 | 53 | Down | ✓ | ✓ |
| | 401 | 239 | Up | ✓ | ✗ |
| | 382 | 226 | Down | ✓ | ✗ |
| | 195 | 34 | Up | ✗ | ✓ |
| | 177 | 33 | Down | ✗ | ✓ |
| | 268 | 1321 | Up | ✗ | ✗ |
| | 286 | 1322 | Down | ✗ | ✗ |

Table S10
*The number of CpGs identified as* DMC *and* NDMC *by* `DMCFB` *versus* `BiSeq`*, classified by CpG Islands, Shores and Deserts, and by the direction of methylation; Chromosomes 15 and 17.*

| | | | Chromosome | 15 | | | 17 | | |
|---|---|---|---|---|---|---|---|---|---|
| | | | Direction `BiSeq` | Equal | Hyper | Hypo | Equal | Hyper | Hypo |
| `DMCFB` | `BiSeq` | Type | `DMCFB` | | | | | | |
| DMC | DMC | Island | Equal | 0 | 0 | 0 | 0 | 0 | 0 |
| | | | Hyper | 0 | 10659 | 0 | 0 | 11231 | 0 |
| | | | Hypo | 0 | 277 | 213 | 0 | 68 | 288 |
| | | Shore | Equal | 0 | 0 | 0 | 0 | 0 | 0 |
| | | | Hyper | 0 | 2102 | 21 | 0 | 2696 | 0 |
| | | | Hypo | 0 | 22 | 148 | 0 | 0 | 461 |
| | | Desert | Equal | 0 | 0 | 0 | 0 | 0 | 0 |
| | | | Hyper | 0 | 1123 | 10 | 0 | 644 | 0 |
| | | | Hypo | 0 | 10 | 342 | 0 | 8 | 196 |
| | NDMC | Island | Equal | 0 | 0 | 0 | 0 | 0 | 0 |
| | | | Hyper | 6754 | 0 | 0 | 14457 | 0 | 0 |
| | | | Hypo | 24984 | 0 | 0 | 41664 | 0 | 0 |
| | | Shore | Equal | 0 | 0 | 0 | 0 | 0 | 0 |
| | | | Hyper | 6615 | 0 | 0 | 21450 | 0 | 0 |
| | | | Hypo | 4194 | 0 | 0 | 11540 | 0 | 0 |
| | | Desert | Equal | 0 | 0 | 0 | 0 | 0 | 0 |
| | | | Hyper | 39352 | 0 | 0 | 70287 | 0 | 0 |
| | | | Hypo | 6827 | 0 | 0 | 14523 | 0 | 0 |
| NDMC | DMC | Island | Equal | 0 | 2636 | 11 | 0 | 2531 | 29 |
| | | | Hyper | 0 | 0 | 0 | 0 | 0 | 0 |
| | | | Hypo | 0 | 0 | 0 | 0 | 0 | 0 |
| | | Shore | Equal | 0 | 449 | 30 | 0 | 395 | 5 |
| | | | Hyper | 0 | 0 | 0 | 0 | 0 | 0 |
| | | | Hypo | 0 | 0 | 0 | 0 | 0 | 0 |
| | | Desert | Equal | 0 | 206 | 254 | 0 | 85 | 53 |
| | | | Hyper | 0 | 0 | 0 | 0 | 0 | 0 |
| | | | Hypo | 0 | 0 | 0 | 0 | 0 | 0 |
| | NDMC | Island | Equal | 29453 | 0 | 0 | 68600 | 0 | 0 |
| | | | Hyper | 0 | 0 | 0 | 0 | 0 | 0 |
| | | | Hypo | 0 | 0 | 0 | 0 | 0 | 0 |
| | | Shore | Equal | 14560 | 0 | 0 | 49128 | 0 | 0 |
| | | | Hyper | 0 | 0 | 0 | 0 | 0 | 0 |
| | | | Hypo | 0 | 0 | 0 | 0 | 0 | 0 |
| | | Desert | Equal | 134185 | 0 | 0 | 200027 | 0 | 0 |
| | | | Hyper | 0 | 0 | 0 | 0 | 0 | 0 |
| | | | Hypo | 0 | 0 | 0 | 0 | 0 | 0 |

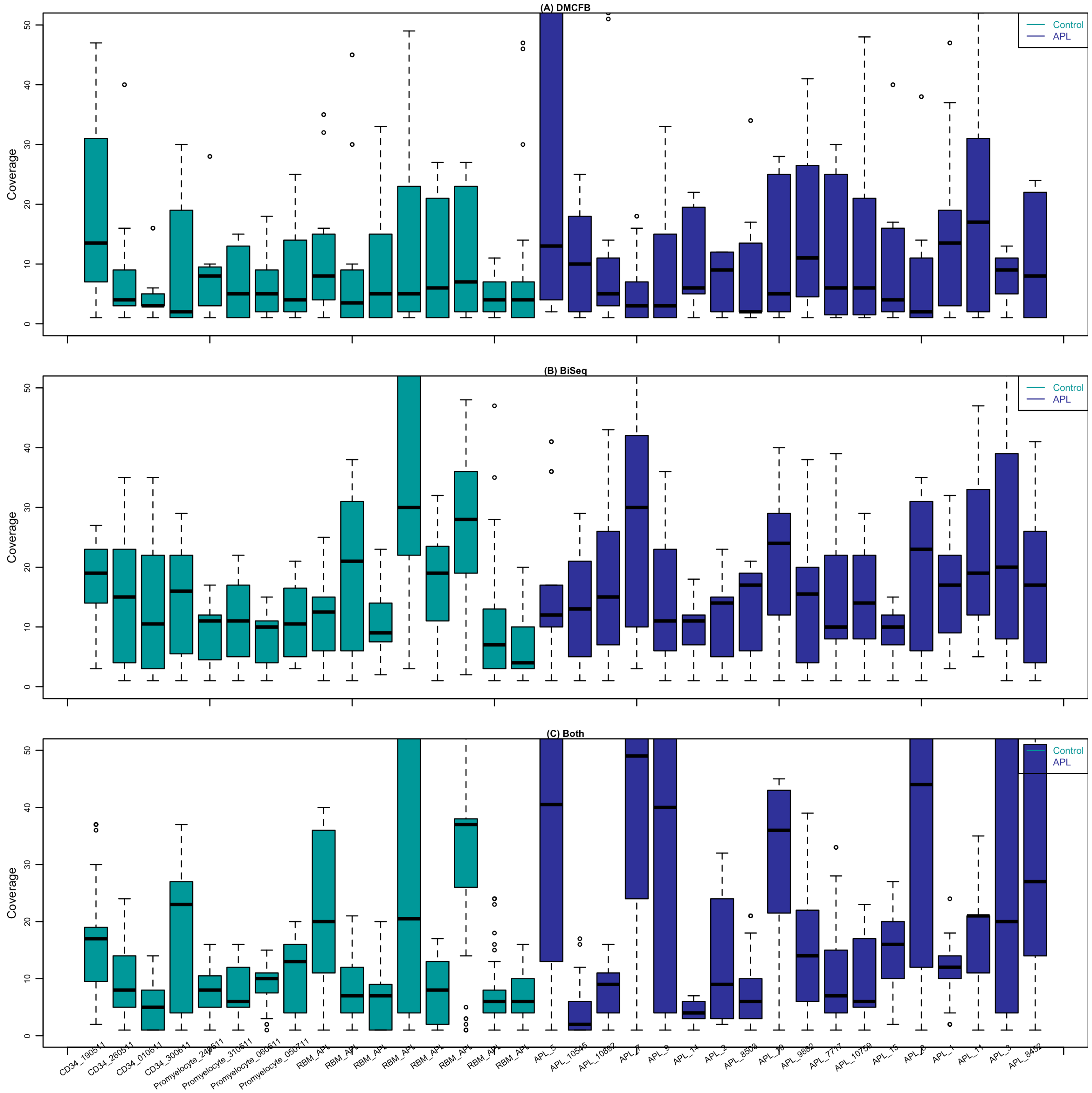


Figure S40: *Box-plots of read-depth of identified* DMCs *by* `DMCFB` *(top graph),* `BiSeq` *(middle graph) and both (bottom graph); Chromosome 15.*

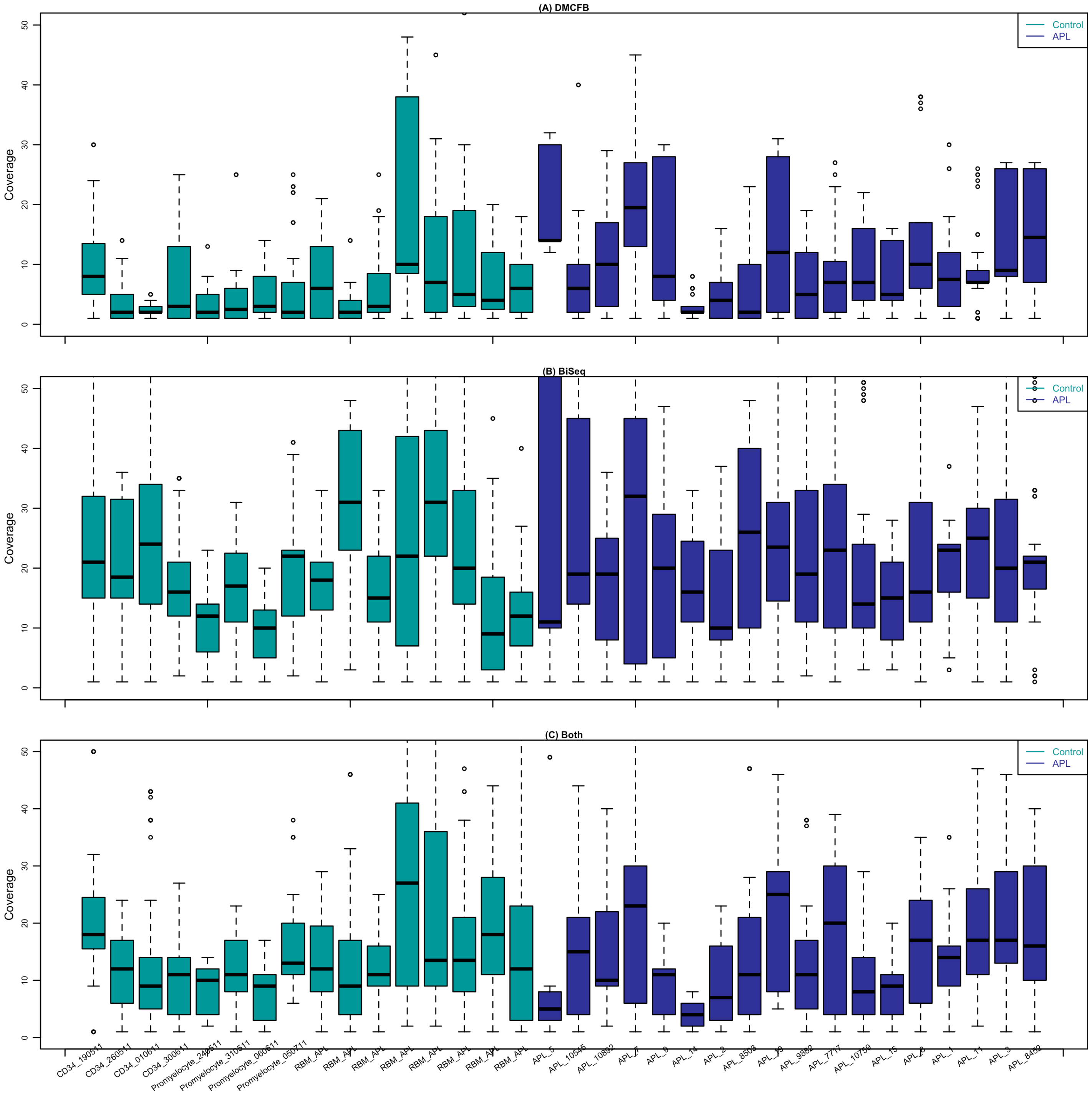


FIGURE S41: *Box-plots of read-depth of identified* DMCs *by* `DMCFB` *(top graph),* `BiSeq` *(middle graph) and both (bottom graph); Chromosome 17.*

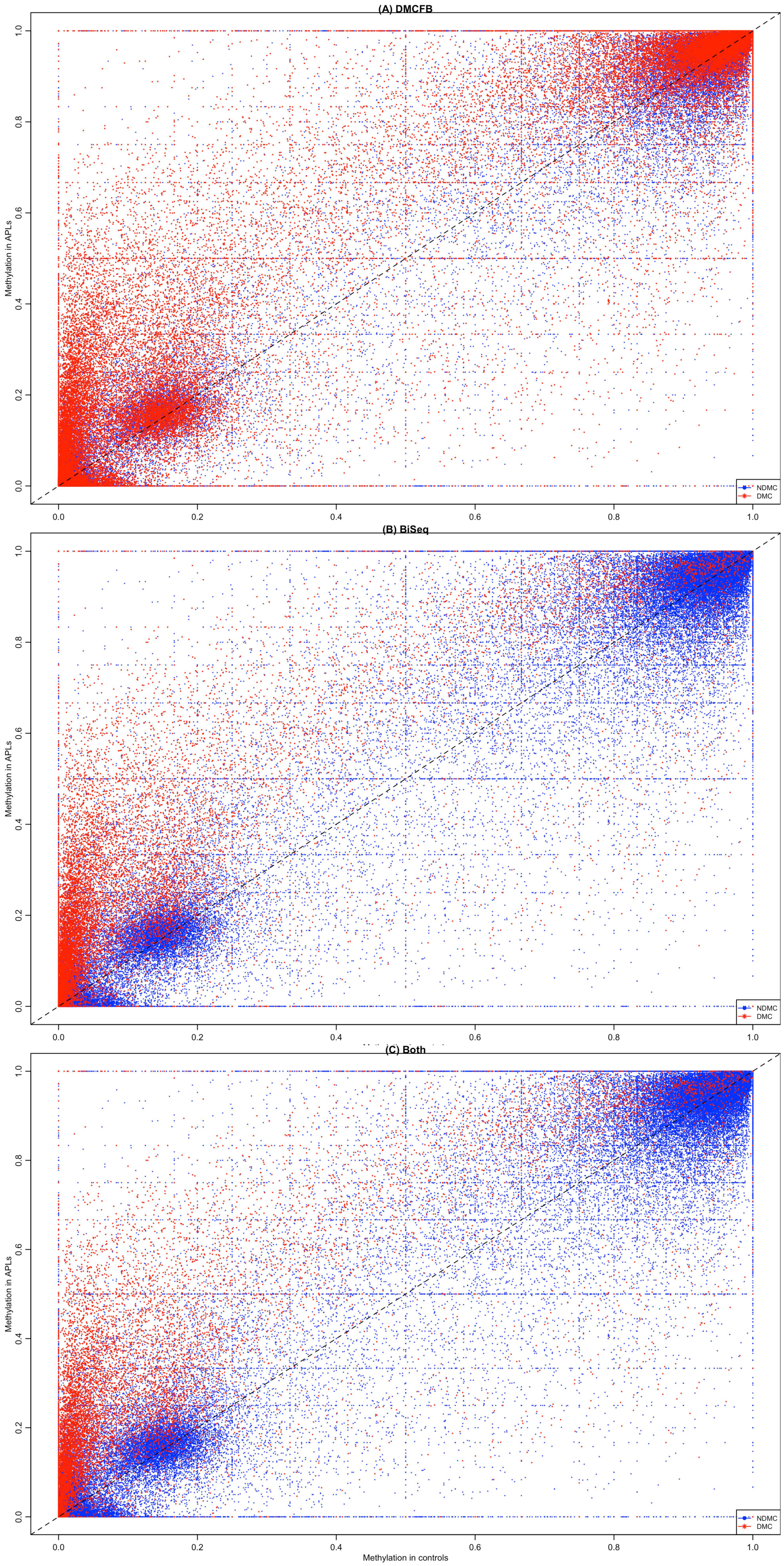


Figure S42: *Raw methylation levels of CpGs for the Control group versus the APL group differentiated for identified* DMCs *(red) and* NDMCs *(blue) by* **DMCFB** *(top graph),* **BiSeq** *(middle graph) and both (bottom graph); Chromosome 15.*

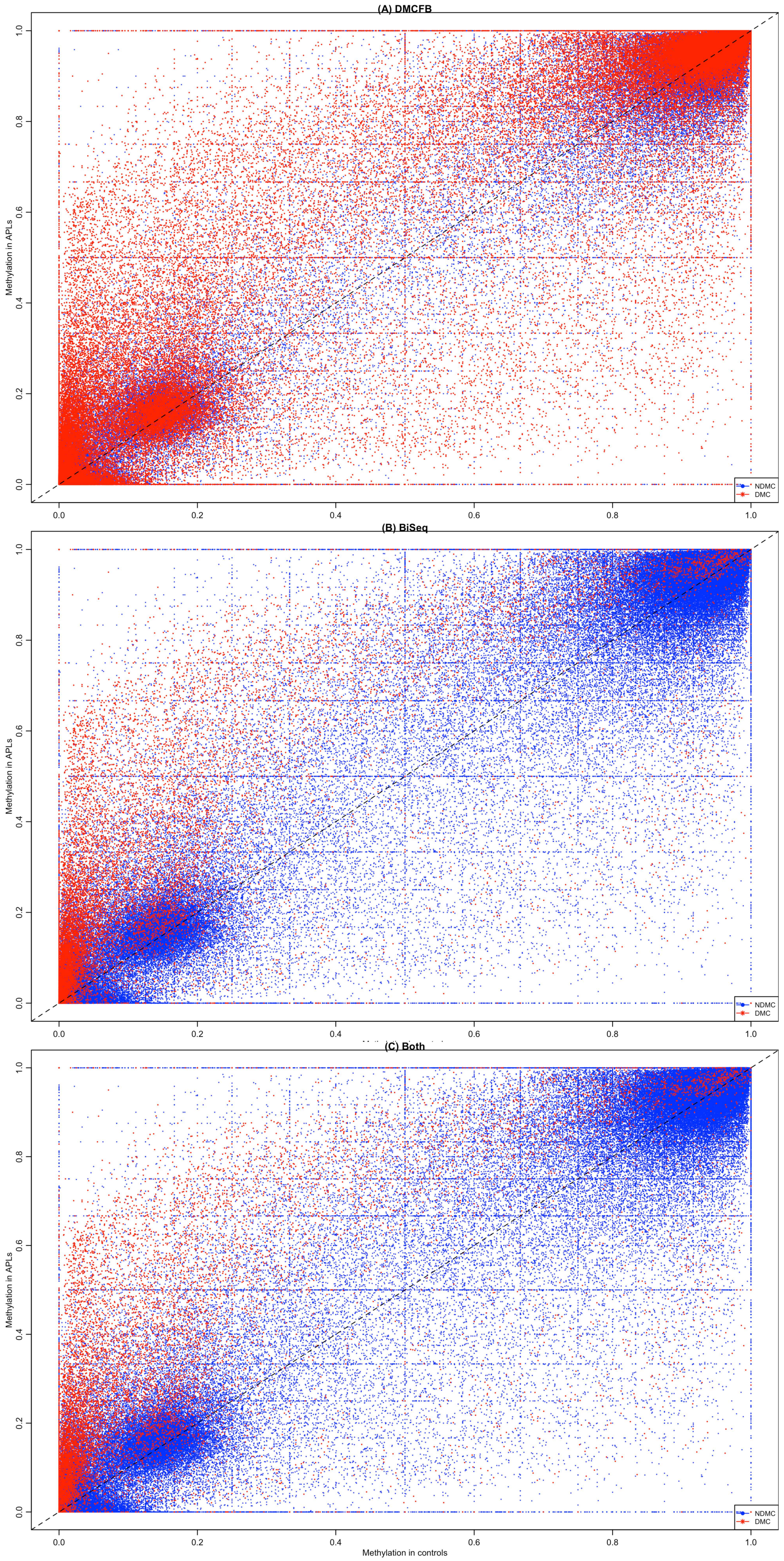


Figure S43: *Raw methylation levels of CpGs for the Control group versus the APL group differentiated for identified* DMCs *(red) and* NDMCs *(blue) by* ***DMCFB*** *(top graph),* ***BiSeq*** *(middle graph) and both (bottom graph); Chromosome 17.*

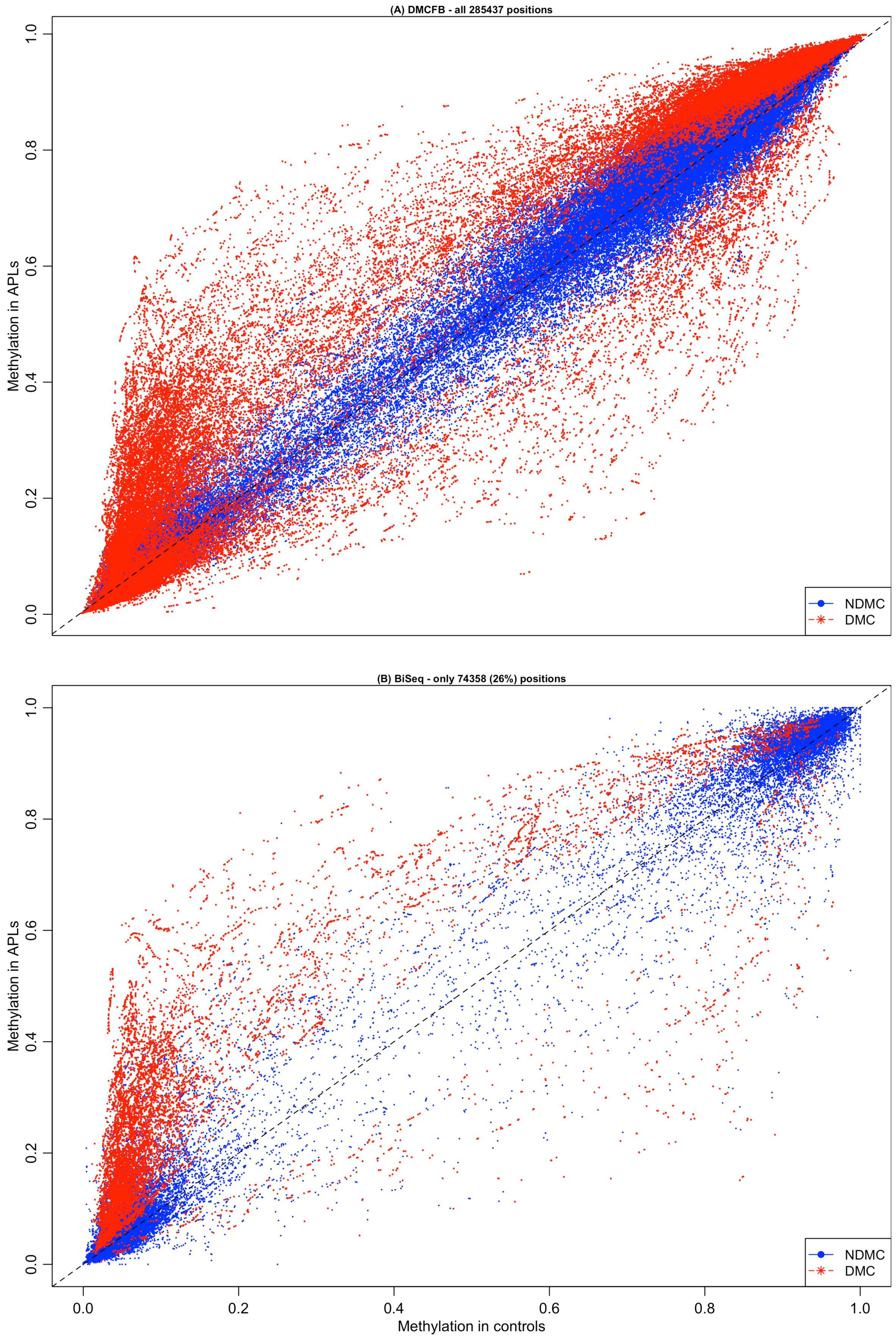


FIGURE S44: *Predicted methylation levels of CpGs for the Control group versus the APL group differentiated for identified* DMCs *(red) and* NDMCs *(blue) by* **DMCFB** *(top graph) and* **BiSeq** *(bottom graph); Chromosome 15.*

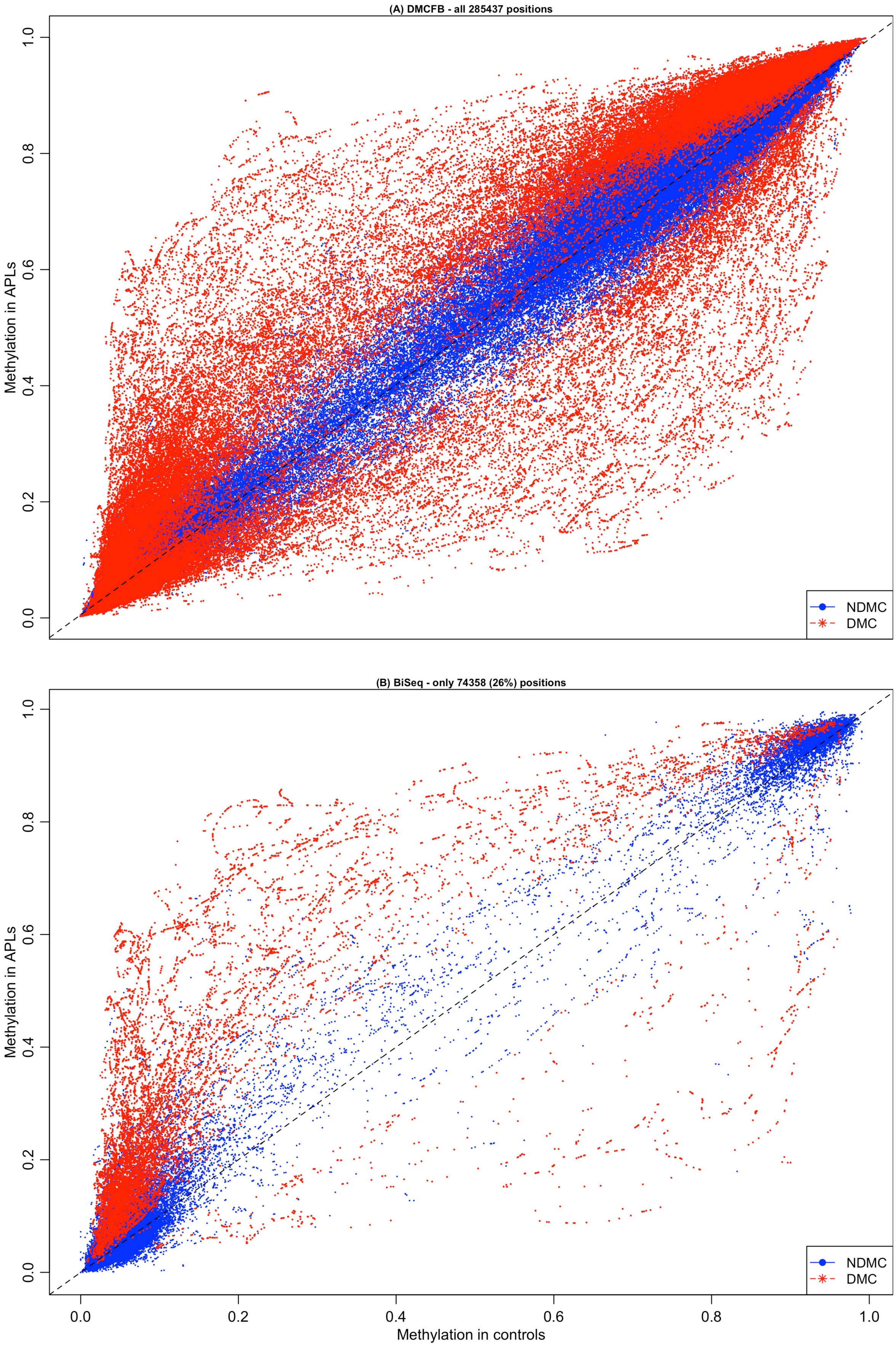


Figure S45: *Predicted methylation levels of CpGs for the Control group versus the APL group differentiated for identified* DMCs *(red) and* NDMCs *(blue) by* `DMCFB` *(top graph) and* `BiSeq` *(bottom graph); Chromosome 17.*

##### *S8.2. Reanalysis of the* APL *data by changing the options in* `BiSeq`

The `BiSeq` pipeline has different options; changing them will result in different percentages of DMCs. In this section, we have changed some of the options to increase the chance of identifying more DMRs leading to more DMCs. Table S11 presents the results. Although the percentage of DMCs has increased in both Chromosome 15 and 17, the agreement between `DMCFB` and `BiSeq` has decreased.

TABLE S11
*Comparing* `DMCFB` *and* `BiSeq` *in identifying* DMCs *in the* APL *data (Default options in* `BiSeq` *are changed to increase the chance of identifying more* DMRs.)
*(A) Chromosome 15*

| | | BiSeq | | | | | |
|---|---|---|---|---|---|---|---|
| **All CpGs** | | NDMC | % of 2x2 table | % of `BiSeq` NDMC | DMC | % of 2x2 table | % of `BiSeq` DMC |
| `DMCFB` | NDMC[1] | 172,510 | 60.44 | | 9,274 | 3.25 | |
| | DMC | 83,212 | 29.15 | 32.54 | 20,441 | 7.16 | 68.79 |
| **CpG Island** | | | | | | | |
| `DMCFB` | NDMC | 24,621 | 32.83 | | 7,479 | 9.97 | |
| | DMC | 27,088 | 36.12 | 52.39 | 15,799 | 21.07 | 67.87 |
| **CpG Shore** | | | | | | | |
| `DMCFB` | NDMC | 13,814 | 49.09 | | 1,225 | 4.35 | |
| | DMC | 9,709 | 34.50 | 41.27 | 3,393 | 12.06 | 73.47 |
| **CpG Desert** | | | | | | | |
| `DMCFB` | NDMC | 134,075 | 73.54 | | 570 | 0.31 | |
| | DMC | 46,415 | 25.46 | 25.72 | 1,249 | 0.69 | 68.66 |

*(B) Chromosome 17*

| | | BiSeq | | | | | |
|---|---|---|---|---|---|---|---|
| **All CpGs** | | NDMC | % of 2x2 table | % of `BiSeq` NDMC | DMC | % of 2x2 table | % of `BiSeq` DMC |
| `DMCFB` | NDMC | 307,858 | 60.32 | | 12,995 | 2.55 | |
| | DMC | 160,568 | 31.46 | 34.28 | 28,965 | 5.68 | 69.03 |
| **CpG Island** | | | | | | | |
| `DMCFB` | NDMC | 61,298 | 44.14 | | 9,862 | 7.10 | |
| | DMC | 49,450 | 35.61 | 44.65 | 18,258 | 13.15 | 64.93 |
| **CpG Shore** | | | | | | | |
| `DMCFB` | NDMC | 47,219 | 55.11 | | 2,309 | 2.70 | |
| | DMC | 29,188 | 34.07 | 38.20 | 6,959 | 8.12 | 75.09 |
| **CpG Desert** | | | | | | | |
| `DMCFB` | NDMC | 199,341 | 69.74 | | 824 | 0.29 | |
| | DMC | 81,930 | 28.66 | 29.13 | 3,748 | 1.31 | 81.98 |

[1] Not a DMC

##### *S8.3. Reanalysis of the* APL *data using* DMCHMM

In this section, we analyze the APL data using DMCHMM and compare the results with those in Table 1(A and B). The statistical model considered in DMCHMM is an HMM with

$$\begin{array}{lll}
\text{Initial probability} & : & p(X_1 = k) = \pi_k, \quad k \in \{0, \ldots, K+1\}, \\
\text{Transition probability} & : & p(X_t = k | X_{t-1} = k') = p_{k',k}, \quad k', k \in \{0, \ldots, K+1\}, \quad X_0 \equiv X_1 \\
\text{Emission probability} & : & Y_t | X_t = k \overset{iid}{\sim} \text{Bin}\left(n_t, \beta_k\right), \quad k \in \{0, \ldots, K+1\},
\end{array}$$

in the first and second step, the Bayesian model

$$p(\boldsymbol{y}|\boldsymbol{x}; \boldsymbol{\beta})p(\boldsymbol{\beta}; \boldsymbol{x}, \boldsymbol{z}, \boldsymbol{\gamma})p(\boldsymbol{x}; \boldsymbol{\pi}, \boldsymbol{P})$$

in the second step, and the logistic model

$$\log\{\beta_{ti}/(1-\beta_{ti})\} = \boldsymbol{Z}_i \boldsymbol{\gamma}_t + \epsilon_{ti}, \quad i = 1, \ldots, 34, \tag{S8.1}$$

in the third step, where $\boldsymbol{Z}_i$ is a vector of categorical variables that represents grouping information (i.e., Control and APL), and $\epsilon_{t,i}$ is a random variable term representing incidental variation for sample $i$ at position $t$. Note that $t = 1, \ldots, T$, where $T = 285,437$ and $510,386$ for Chromosome 15 and 17, respectively

TABLE S12

*The percentage of CpGs identified as* DMC *by* DMCFB *and* DMCHMM *displayed; Chromosomes 15 and 17.*

| Chromosome | DMCFB | DMCHMM |
|---|---|---|
| 15 | 36.31 | 17.36 |
| 17 | 37.14 | 9.43 |

TABLE S13

*The number of CpGs identified as* DMC *by only* DMCFB*, only* DMCHMM *and both methods at the same time, separated for CpG Islands and Shores; Chromosomes 15 and 17.*

| Chromosome | Location | DMCFB | DMCHMM | Both |
|---|---|---|---|---|
| 15 | Islands | 594 | 764 | 594 |
| | Shores | 823 | 1452 | 823 |
| 17 | Islands | 1171 | 1574 | 1171 |
| | Shores | 1931 | 3183 | 1931 |

Table S14

*Counts of Shore regions are shown, categorized by whether the adjacent Island contains at least one* DMC *and whether the Shore (in the same direction) contains at least one* DMC. *Counts are shown* `DMCFB` *and* `DMCHMM`*; Chromosomes 15 and 17.*

| | # of Shores | | | With DMC | |
|---|---|---|---|---|---|
| Chromosome | `DMCFB` | `DMCHMM` | Stream | Island | Shore |
| 15 | 372 | 702 | Up | ✓ | ✓ |
| | 363 | 709 | Down | ✓ | ✓ |
| | 222 | 62 | Up | ✓ | ✗ |
| | 231 | 55 | Down | ✓ | ✗ |
| | 38 | 21 | Up | ✗ | ✓ |
| | 50 | 20 | Down | ✗ | ✓ |
| | 160 | 7 | Up | ✗ | ✗ |
| | 148 | 8 | Down | ✗ | ✗ |
| 17 | 770 | 1535 | Up | ✓ | ✓ |
| | 789 | 1541 | Down | ✓ | ✓ |
| | 401 | 39 | Up | ✓ | ✗ |
| | 382 | 33 | Down | ✓ | ✗ |
| | 195 | 52 | Up | ✗ | ✓ |
| | 177 | 55 | Down | ✗ | ✓ |
| | 268 | 8 | Up | ✗ | ✗ |
| | 286 | 5 | Down | ✗ | ✗ |

TABLE S15
*The number of CpGs identified as* DMC *and* NDMC *by* DMCFB *versus* DMCHMM*, classified by CpG Islands, Shores and Deserts, and by the direction of methylation; Chromosomes 15 and 17.*

| | | | Chromosome | 15 | | | 17 | | |
|---|---|---|---|---|---|---|---|---|---|
| | | | Direction DMCHMM | Equal | Hyper | Hypo | Equal | Hyper | Hypo |
| DMCFB | DMCHMM | Type | DMCFB | | | | | | |
| DMC | DMC | Island | Equal | 0 | 0 | 0 | 0 | 0 | 0 |
| | | | Hyper | 0 | 7299 | 0 | 0 | 4343 | 0 |
| | | | Hypo | 0 | 1974 | 1 | 0 | 2306 | 63 |
| | | Shore | Equal | 0 | 0 | 0 | 0 | 0 | 0 |
| | | | Hyper | 0 | 3045 | 0 | 0 | 3232 | 16 |
| | | | Hypo | 0 | 537 | 51 | 0 | 378 | 264 |
| | | Desert | Equal | 0 | 0 | 0 | 0 | 0 | 0 |
| | | | Hyper | 0 | 6612 | 0 | 0 | 6953 | 109 |
| | | | Hypo | 0 | 812 | 0 | 0 | 1383 | 154 |
| | NDMC | Island | Equal | 0 | 0 | 0 | 0 | 0 | 0 |
| | | | Hyper | 87 | 8543 | 1484 | 130 | 17047 | 4168 |
| | | | Hypo | 249 | 16405 | 6845 | 532 | 24999 | 14120 |
| | | Shore | Equal | 0 | 0 | 0 | 0 | 0 | 0 |
| | | | Hyper | 19 | 4559 | 1115 | 173 | 15446 | 5279 |
| | | | Hypo | 49 | 2352 | 1375 | 94 | 6971 | 4294 |
| | | Desert | Equal | 0 | 0 | 0 | 0 | 0 | 0 |
| | | | Hyper | 357 | 25421 | 8095 | 821 | 41198 | 21870 |
| | | | Hypo | 97 | 3840 | 2430 | 110 | 6324 | 6756 |
| NDMC | DMC | Island | Equal | 0 | 4651 | 10 | 0 | 5462 | 12 |
| | | | Hyper | 0 | 0 | 0 | 0 | 0 | 0 |
| | | | Hypo | 0 | 0 | 0 | 0 | 0 | 0 |
| | | Shore | Equal | 0 | 2801 | 90 | 0 | 4389 | 30 |
| | | | Hyper | 0 | 0 | 0 | 0 | 0 | 0 |
| | | | Hypo | 0 | 0 | 0 | 0 | 0 | 0 |
| | | Desert | Equal | 0 | 21605 | 57 | 0 | 18703 | 313 |
| | | | Hyper | 0 | 0 | 0 | 0 | 0 | 0 |
| | | | Hypo | 0 | 0 | 0 | 0 | 0 | 0 |
| | NDMC | Island | Equal | 336 | 20513 | 6590 | 798 | 46277 | 18611 |
| | | | Hyper | 0 | 0 | 0 | 0 | 0 | 0 |
| | | | Hypo | 0 | 0 | 0 | 0 | 0 | 0 |
| | | Shore | Equal | 102 | 8361 | 3685 | 470 | 30161 | 14479 |
| | | | Hyper | 0 | 0 | 0 | 0 | 0 | 0 |
| | | | Hypo | 0 | 0 | 0 | 0 | 0 | 0 |
| | | Desert | Equal | 1267 | 83068 | 28648 | 2540 | 110857 | 67752 |
| | | | Hyper | 0 | 0 | 0 | 0 | 0 | 0 |
| | | | Hypo | 0 | 0 | 0 | 0 | 0 | 0 |

TABLE S16

*Comparing DMCFB and DMCHMM in identifying DMCs in the APL data.*
*(A) Chromosome 15*

| | | DMCHMM | | | | | |
|---|---|---|---|---|---|---|---|
| **All CpGs** | | NDMC | % of 2x2 table | % of DMCHMM NDMC | DMC | % of 2x2 table | % of DMCHMM DMC |
| DMCFB | NDMC[1] | 152,570 | 53.45 | | 29,214 | 10.23 | |
| | DMC | 83,322 | 29.19 | 35.32 | 20,331 | 7.12 | 41.03 |
| **CpG Island** | | | | | | | |
| DMCFB | NDMC | 27,439 | 36.59 | | 4,661 | 6.22 | |
| | DMC | 33,613 | 44.83 | 55.06 | 9,274 | 12.37 | 66.55 |
| **CpG Shore** | | | | | | | |
| DMCFB | NDMC | 12,148 | 43.17 | | 2,891 | 10.27 | |
| | DMC | 9,469 | 33.65 | 43.80 | 3,633 | 12.91 | 55.69 |
| **CpG Desert** | | | | | | | |
| DMCFB | NDMC | 112,983 | 61.97 | | 21,662 | 11.88 | |
| | DMC | 40,240 | 22.07 | 26.26 | 7,424 | 4.07 | 25.52 |

*(B) Chromosome 17*

| | | DMCHMM | | | | | |
|---|---|---|---|---|---|---|---|
| **All CpGs** | | NDMC | % of 2x2 table | % of DMCHMM NDMC | DMC | % of 2x2 table | % of DMCHMM DMC |
| DMCFB | NDMC | 291,944 | 57.20 | | 28,909 | 5.66 | |
| | DMC | 170,332 | 33.37 | 36.85 | 19,201 | 3.76 | 39.91 |
| **CpG Island** | | | | | | | |
| DMCFB | NDMC | 65,686 | 47.30 | | 5,474 | 3.94 | |
| | DMC | 60,996 | 43.92 | 48.15 | 6,712 | 4.83 | 55.08 |
| **CpG Shore** | | | | | | | |
| DMCFB | NDMC | 45,109 | 52.65 | | 4,419 | 5.16 | |
| | DMC | 32,257 | 37.65 | 41.69 | 3,890 | 4.54 | 46.82 |
| **CpG Desert** | | | | | | | |
| DMCFB | NDMC | 181,149 | 63.37 | | 19,016 | 6.65 | |
| | DMC | 77,079 | 26.97 | 29.85 | 8,599 | 3.01 | 31.14 |

[1]Not a DMC

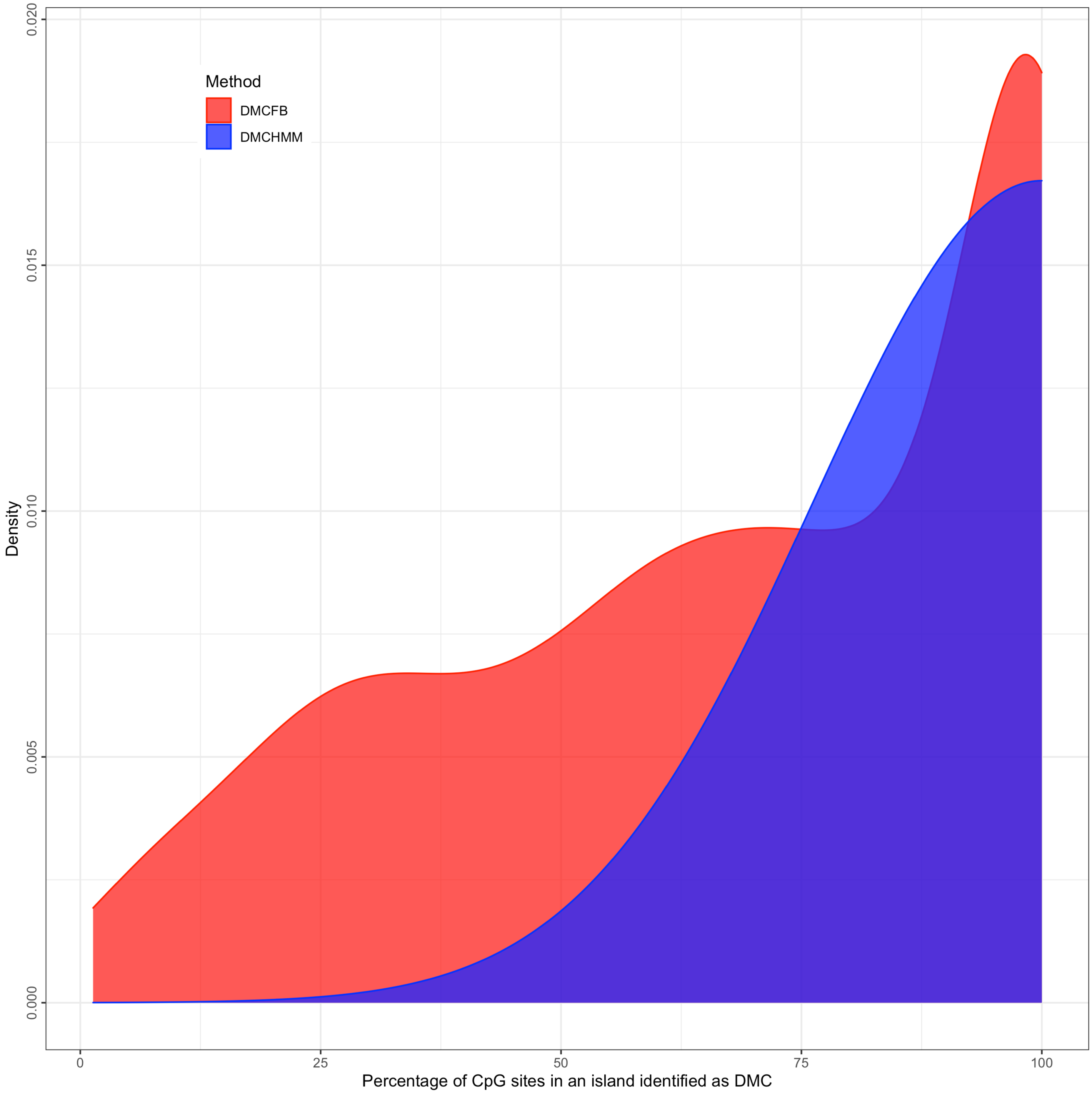


FIGURE S46: *The density of percentage of CpGs in an Island identified as* DMC *by* `DMCFB` *(red) and* `DMCHMM` *(blue); Chromosome 15.*

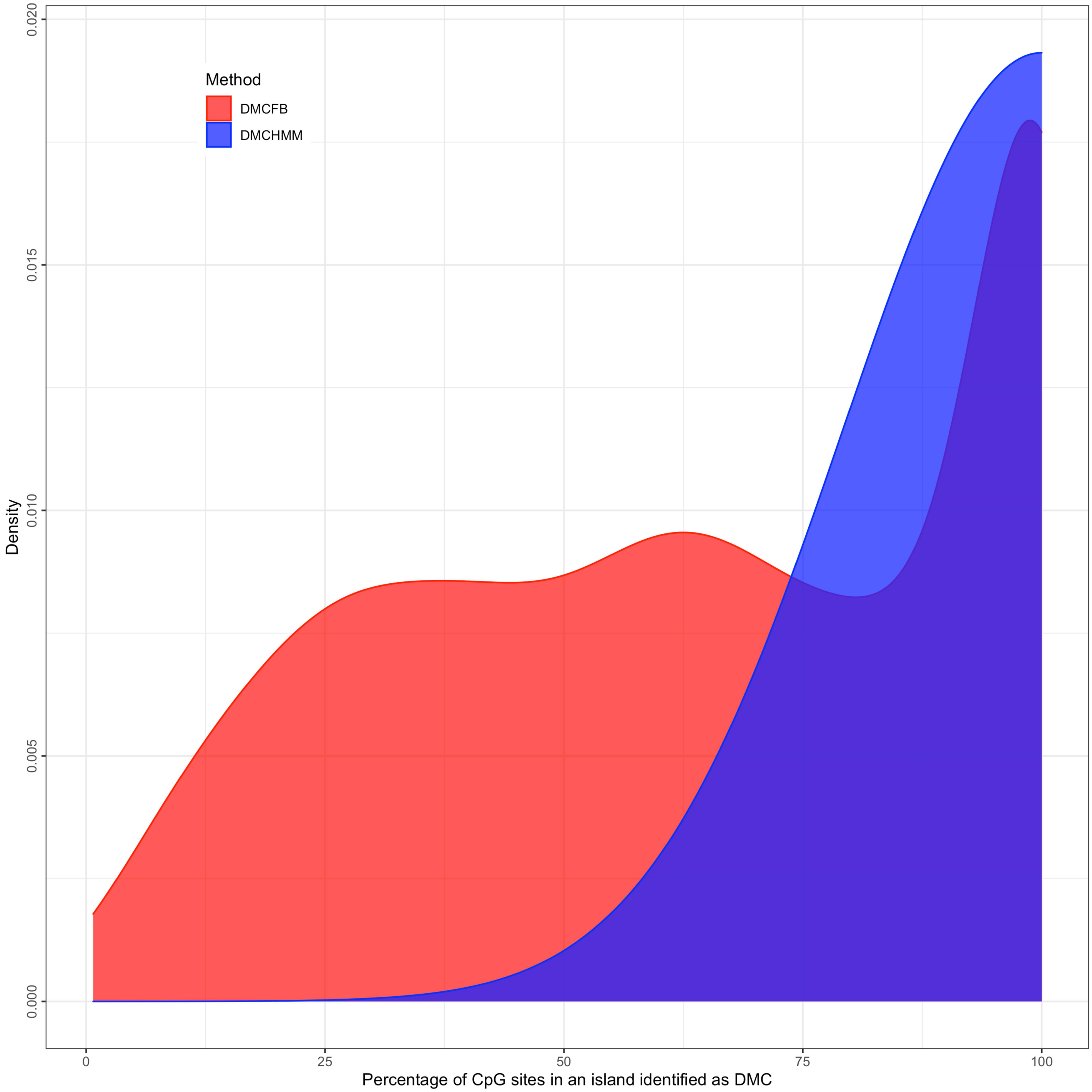


FIGURE S47: *The density of percentage of CpGs in an Island identified as* DMC *by* `DMCFB` *(red) and* `DMCHMM` *(blue); Chromosome 17.*

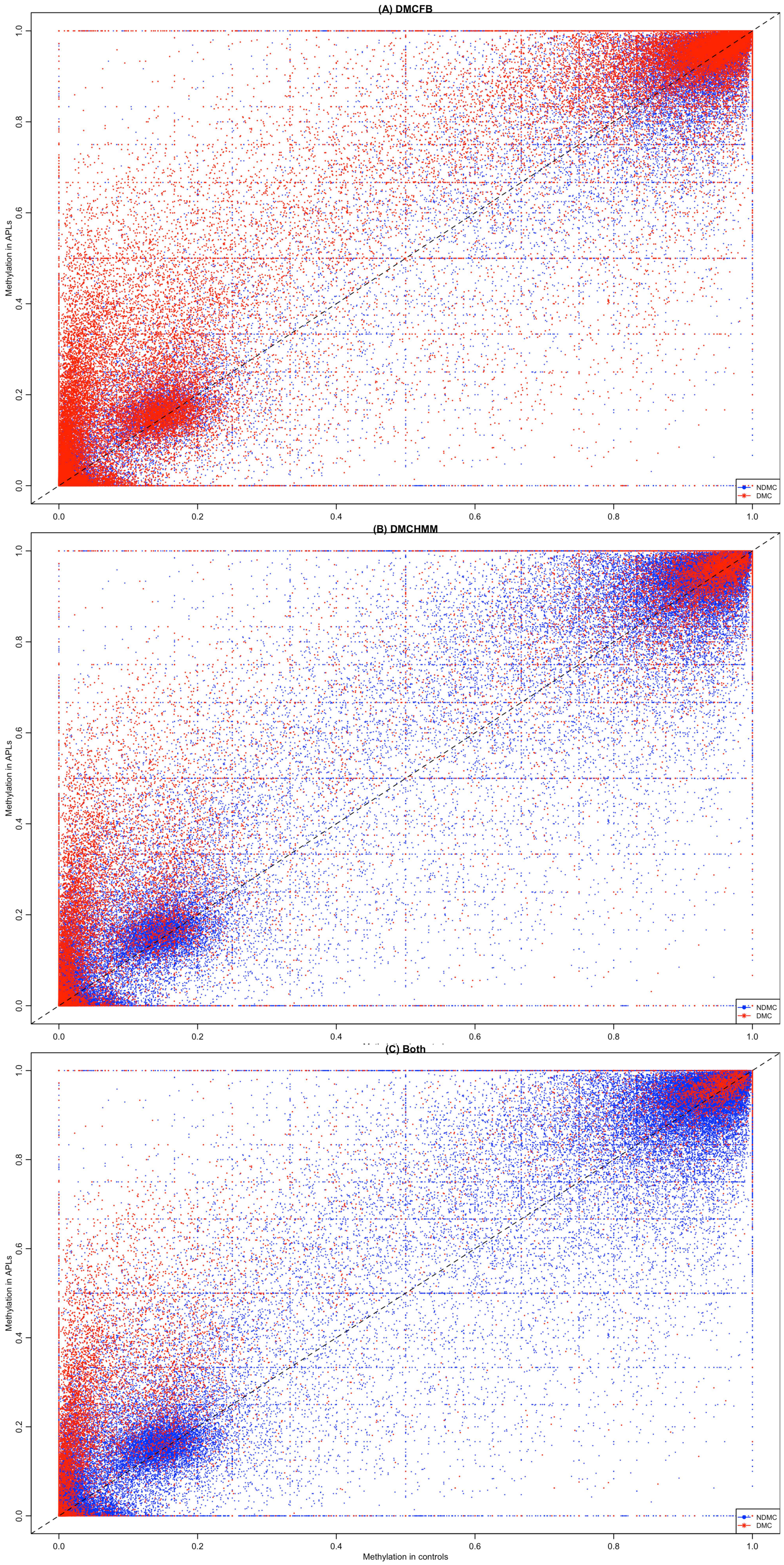


FIGURE S48: *Raw methylation levels of CpGs for the Control group versus the APL group differentiated for identified* DMCs *(red) and* NDMCs *(blue) by* ***DMCFB*** *(top graph),* ***DMCHMM*** *(middle graph) and both (bottom graph); Chromosome 15.*

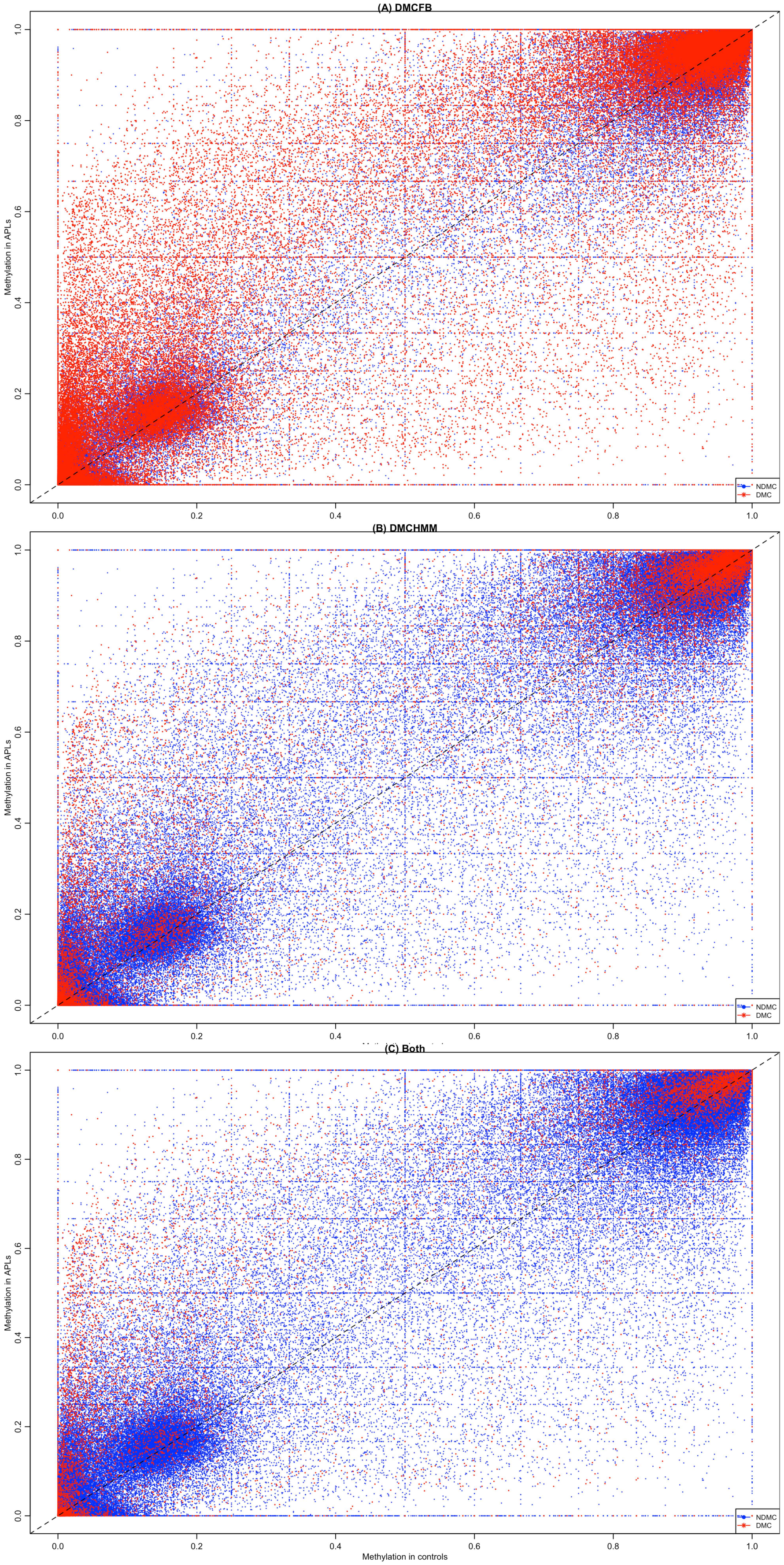


Figure S49: *Raw methylation levels of CpGs for the Control group versus the APL group differentiated for identified* DMCs *(red) and* NDMCs *(blue) by* `DMCFB` *(top graph),* `DMCHMM` *(middle graph) and both (bottom graph); Chromosome 17.*

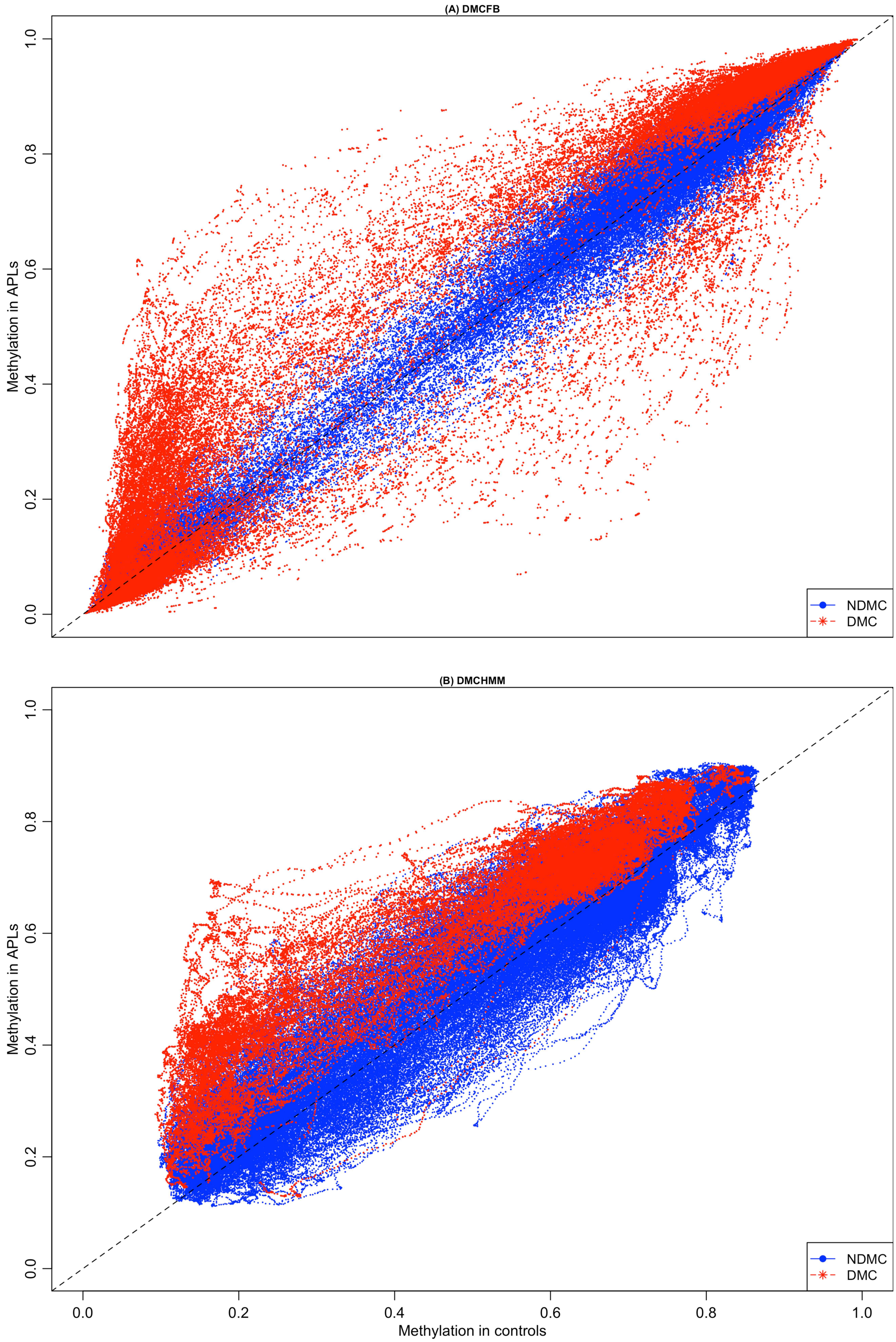


Figure S50: *Predicted methylation levels of CpGs for the Control group versus the APL group differentiated for identified* DMCs *(red) and* NDMCs *(blue) by* `DMCFB` *(top graph) and* `DMCHMM` *(bottom graph); Chromosome 15.*

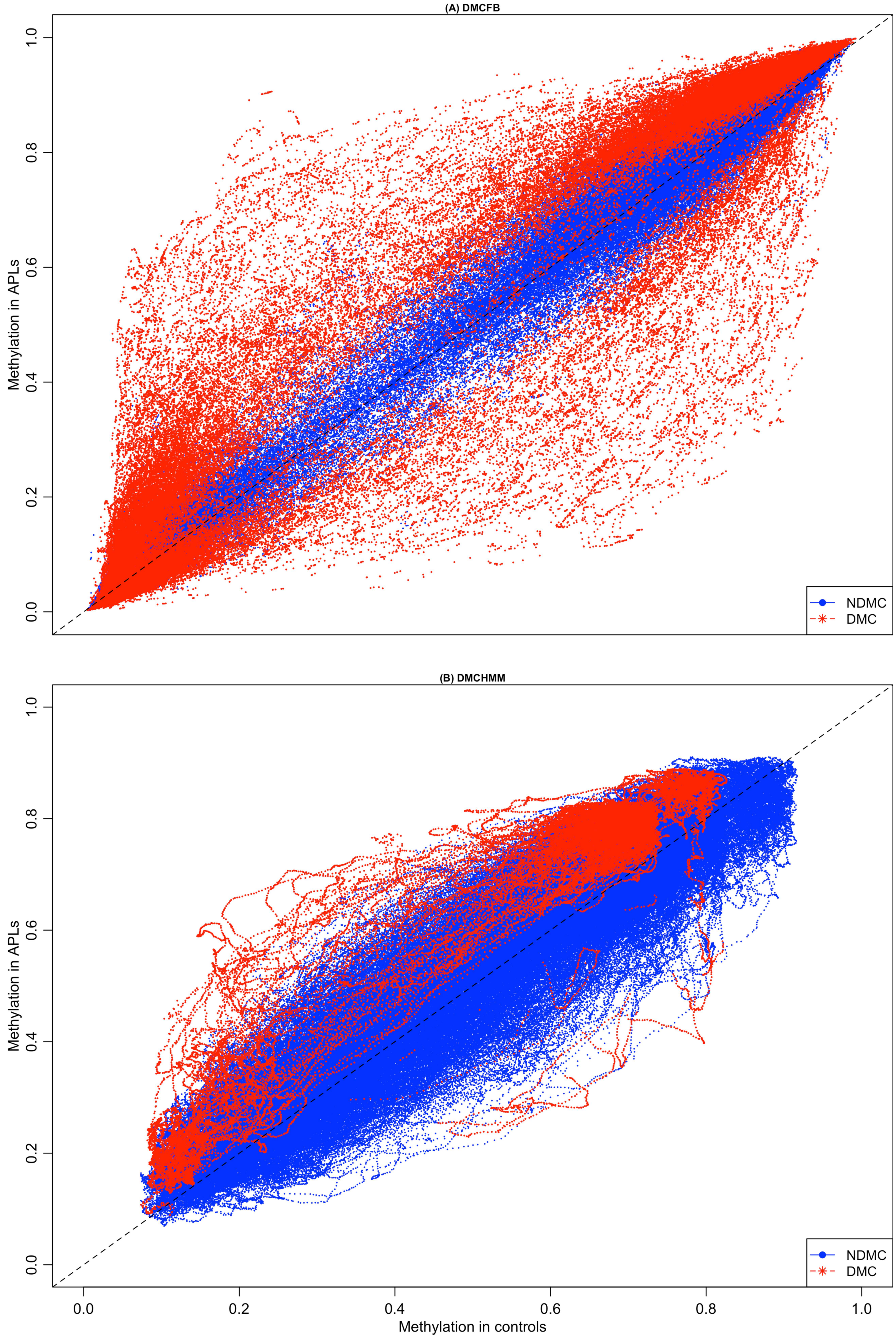


FIGURE S51: *Predicted methylation levels of CpGs for the Control group versus the APL group differentiated for identified* DMCs *(red) and* NDMCs *(blue) by* `DMCFB` *(top graph) and* `DMCHMM` *(bottom graph); Chromosome 17.*
